# Peroxisome quality control is gated by a dynamic ATF6α–ABCD3 tether

**DOI:** 10.64898/2026.10.05.756567

**Authors:** Chari M. Noddings, Rukmini Mukherjee, Carsten Peukert, Thomas G. Laughlin, Jing Wang, Benjamin T. Walters, Tsan-Wen Lu, Tristan Croll, Deniz Eismann, Shelly Harel, Angela Xu, Kibeom Kim, Meghna Gupta, James J. Crawford, Christopher P. Arthur, Daniel Itzhak, Mauro Costa-Mattioli, Joachim Rudolph, Avi Ashkenazi, Peter Walter

## Abstract

The endoplasmic reticulum (ER) coordinates protein folding and lipid biosynthesis, while peroxisomes govern fatty-acid catabolism and redox balance. However, little is known about the mechanisms by which these organelles communicate. Here we show that activating transcription factor 6 (ATF6α), a cytoprotective ER-resident sensor of the unfolded protein response (UPR)^1–4^, forms a molecular tether that zippers the ER and peroxisomes together at a 23-nm intermembrane distance. ATF6α achieves this by inserting a short segment of its unstructured N-terminal cytosolic tail into the solvated transmembrane cavity of the peroxisomal ABC-type lipid transporter ABCD3, trapping it in an off-state. This interaction is reinforced by Ceapin, a previously identified small molecule molecular-glue ligand, which promotes ER–peroxisome contact formation^5–7^. The junctions serve as a scaffold that can initiate peroxisome turnover by p62-triggered autophagy (pexophagy), as enhanced lipid processing resulting from organelle zippering leads to peroxisome damage due to excessive reactive oxygen species (ROS) accumulation. These findings define ATF6α as a non-transcriptional regulator of organelle quality control and provide a template for engineering Ceapin analogues against ATF6α/ABCD3-dependent colorectal and hepatocellular cancers and related diseases.

## Introduction

ATF6 senses endoplasmic reticulum (ER) stress and, together with IRE1 and PERK, helps coordinate the unfolded protein response (UPR)^8–10^. Of the three UPR branches, ATF6 remains the least well understood. ATF6 is a type II single-pass transmembrane protein, with an N-terminal cytosolic bZIP transcription factor domain, preceded by an intrinsically disorder tail, and a stress-sensing ER-lumenal domain^2,11^. ATF6 has two paralogs, ATF6α and ATF6β, which are both ubiquitously expressed, but the role of ATF6β in ER stress resolution is unclear. ATF6α is cytoprotective, its transcriptional targets mostly consist of genes encoding components of the ER’s protein folding machinery^12–14^. Aberrant ATF6α activation can drive cancer progression and chemoresistance, marking it as a potential therapeutic target^15–18^. However, ATF6α has long been considered ‘undruggable’ due to its lack of identifiable binding sites. To activate, ATF6α must exit the ER by vesicular trafficking to the Golgi, where its bZIP module is proteolytically released from the membrane by the S1 and S2 proteases before entering the nucleus to drive target-gene transcription^1–4^. We previously identified small molecule inhibitors of ATF6α through chemical library screening^5,6^, termed Ceapins—from the Irish word for “trap”— for their ability to retain ATF6α in the ER and prevent its trafficking to the Golgi apparatus. Ceapin A7, the most potent Ceapin analog, is highly specific to trap ATF6α in the ER and has no effect on its paralog, ATF6β^6^.

A genome-wide CRISPR interference screen subsequently identified the peroxisomal ABC transporter ABCD3 as the target of Ceapin A7, revealing that the compound traps ATF6α by tethering its intrinsically disordered N-terminus (ATF6α 2–90) to ABCD3 at the ER–peroxisome interface^7^. We previously characterized this interaction as “neomorphic”, induced by the compound rather than occurring spontaneously. In addition to the Ceapin-stabilized ATF6α•ABCD3 tether, VAP family proteins have been shown to form ER–peroxisome membrane contact sites to regulate lipid shuttling and peroxisome positioning^19–21^. Endogenous depletion of VAP proteins does not affect the ATF6α•ABCD3•Ceapin A7 interaction, confirming this ER– peroxisome contact site is distinct from the VAP-mediated contact^7^.

Peroxisomes are essential organelles that carry out β-oxidation of long-chain and branched-chain fatty acids, perform biosynthesis of critical lipid molecules, and maintain cellular redox homeostasis^22,23^. Lipid substrates are imported into the peroxisome by ATP-binding cassette (ABC) transporters: ABCD1, ABCD2, and ABCD3, of which ABCD3 is the most abundant. These transporters import fatty acid-coenzyme A thioesters into the peroxisome to be metabolized^24^. Each transporter functions as an obligate homodimer with a transmembrane domain (TMD) and nucleotide binding domain (NBD)^24,25^. ABCD3 is the most highly and ubiquitously expressed of the three transporters^26^ and is necessary for peroxisomal import of long-chain fatty acids, branched-chain fatty acids, dicarboxylic acids, and bile acid intermediates^27–31^. Characteristic of an ABC transporter, substrate translocation into the peroxisome is coupled to ATP binding and hydrolysis, which drives conformational changes of ABCD3, as recently revealed by cryogenic electron microscopy (cryo-EM)^32,33^. In the apo state, ABCD3 adopts an inward-facing cytosol-open conformation that binds lipid substrates from the cytosol. Upon ATP binding, the NBDs dimerize, forming the outward-facing open conformation with the transmembrane cavity open toward the peroxisome lumen (hereafter, ‘peroxisome-open conformation’), allowing release of the lipid substrate into the peroxisome. ATP hydrolysis destabilizes NBD association and resets the transporter to the cytosol-open apo conformation, restarting the cycle.

ATF6α has not previously been implicated in peroxisomal biology, but its function extends beyond transcriptional control of proteostasis genes to lipid metabolism and stress adaptation more broadly^18,34–37^. ATF6α activation is linked to lipid sensing in the ER membrane^34^ and ATF6α upregulates transcription of peroxisomal lipid metabolism genes (e.g., FAR1, PPARα, ACOX1)^13,14,38–40^. During ischemia–reperfusion injury *in vivo*, ATF6α transcriptionally activates antioxidant genes, including peroxisome-localized catalase and peroxiredoxins^35^. These findings point to a potential functional link between ATF6α signaling and peroxisomal function, but the precise mechanistic connection has not been established.

Cells balance peroxisome quality and quantity through the interplay between peroxisome biogenesis and autophagic removal. Pexophagy, a form of selective autophagy, eliminates peroxisomes in response to oxidative stress, nutrient deprivation, or organelle dysfunction^41–44^. E3 ligases ubiquitinate resident membrane proteins on damaged peroxisomes, which are recognized by autophagy receptors such as p62/SQSTM1 and NBR1^45–47^, linking ubiquitylated cargo to the autophagic machinery, promoting selective sequestration and lysosomal degradation of peroxisomes. The ER also plays a central role in autophagy, including in selective pathways such as reticulophagy and ER-phagy, in which ER membranes contribute to the recognition and turnover of cellular components^48,49^. Whether and how ER-peroxisome contact sites and metabolic regulation are coupled to the initiation of pexophagy is largely unknown. Here, we show that ATF6α forms a direct, non-transcriptional link between ER–peroxisome contact sites, lipid metabolism, and organelle quality control—an unexpected scaffolding role that places ATF6α at the center of peroxisomal homeostasis.

## Results

### Ceapin A7 stabilizes the inter-organelle tether formed by ATF6α and ABCD3

GFP-tagged ATF6α (GFP-ATF6α) localizes to the ER in untreated cells, but it rapidly redistributes into puncta upon Ceapin A7 treatment (**Fig. 1A**), consistent with previous results^5,6^. We previously showed that these puncta mark sites where ER-localized ATF6α engages peroxisomal ABCD3 to form an inter-organelle junction^7^. To visualize the tether between ATF6α and ABCD3, we performed *in situ* cryo-CLEM (correlative light and electron microscopy), which uses fluorescence imaging to locate a region of interest before imaging that same region by electron microscopy. To this end, we thinned vitrified U2OS cells into lamellae using cryo-FIB (focused ion beam milling, which carves frozen cells into thin electron-transparent sections) and localized GFP-ATF6α puncta by cryo-fluorescence microscopy, then imaged these regions by cryo-ET (electron tomography), reconstructing a three-dimensional volume from a tilt series of two-dimensional images (**Fig. 1B**). We collected 13 tomograms from Ceapin A7-treated cells containing GFP-ATF6α puncta and 4 tomograms from untreated controls (**Fig. 1C, Extended Data Fig. 1A**). At these puncta, we consistently observed clusters of peroxisomes enmeshed in the ER network, with the ER membrane extensively wrapped around peroxisomes at a uniform intermembrane distance of 22.6 ± 3.8 nm (**Extended Data Fig. 1B**). Ribosomes stud the ER membrane proximal to the extended contact zones but are excluded from ER-peroxisome junctions (**Fig. 1C**, top). In control cells, no peroxisomes clustered or formed extended contact zones with the ER. Interestingly, peroxisomes in tomograms from Ceapin A7-treated cells were on average ∼2-times larger in cross section area compared to control cells (**Extended Data Fig. 1C**). This indicates swelling, possibly caused by peroxisomal dysfunction or damage^29,50,51^, as discussed below. We did not discern any ultrastructural intermembrane features at the ER-peroxisome contact sites, likely due to the relatively small size of the ATF6α•ABCD3•Ceapin A7 tethering complex (∼150 kDa).

**Figure 1:**
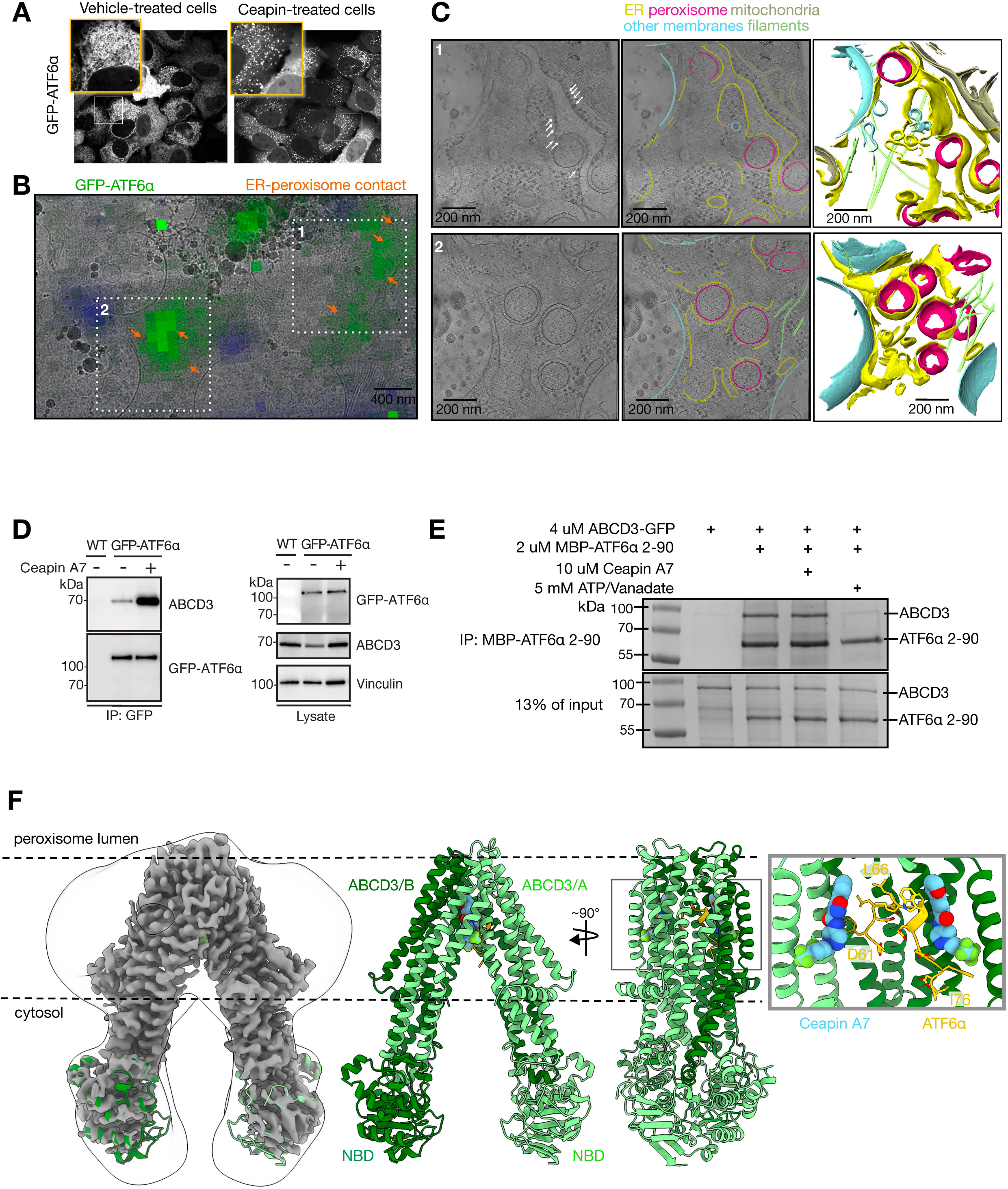
ATF6α and ABCD3 form an inter-organelle tether, stabilized by Ceapin A7. **a**, Light microscopy of U2OS cells expressing GFP-ATF6α with vehicle treatment (left) or 10 μM Ceapin A7 treatment (right) after 1 hour. Ceapin-treated cells show clear GFP-ATF6α puncta. **b**, Correlation of GFP-ATF6α signal (green) with TEM image of U2OS cells treated with 10 μM Ceapin A7 for 1 hour and thinned to ∼200 nm with cryo-FIB. White boxes indicate areas where corresponding tomograms were acquired in **c**. Orange arrows indicate ER-peroxisome contact sites. Blue signal indicates lipid droplets. **c**, Left panel, Z-slices of two representative tomograms corresponding to white box in panel **b** (n = 12). White arrows indicate ribosomes studded along the ER. Middle panel, annotated tomograms corresponding to left panel. ER (yellow), peroxisomes (pink), mitochondria (tan), filaments (green), and other non-ER membranes (blue). Right panel, 3D rendered image of structures in the corresponding tomograms. ER (yellow), peroxisomes (pink), mitochondria (tan), filaments (green), and other non-ER membranes (blue). **d**, Western blot of GFP-ATF6α immunoprecipitation from U2OS cells with or without 10 μM Ceapin A7 treatment for 4 hours. **e**, Coomassie-stained SDS-PAGE from *in vitro* immunoprecipitation of 2 μM purified MBP-ATF6α 2-90 with 4 μM purified ABCD3 in the presence of 10 uM Ceapin A7 or 5 mM ATP/Vanadate. **f**, Left, 2.9 Å cryo-EM map of the ABCD3•ATF6α•Ceapin A7 complex. Black outline is an overlay of a 20 Å lowpass-filtered map showing the boundary of the detergent micelle. Dashed black line indicates where the peroxisome membrane would surround ABCD3. Middle, atomic model of the ABCD3•ATF6α•Ceapin A7 complex corresponding to cryo-EM map. ABCD3 promoter A (light green), ABCD3 promoter B (dark green), ATF6α (yellow), and Ceapin A7 (blue). Color scheme is maintained throughout. Nucleotide binding domains (NBDs) of ABCD3 are indicated. Right, 90-degree rotation from middle and central slice through structure to show inside of the ABCD3 chamber containing ATF6α 61-76 peptide and the two symmetrically bound Ceapin A7 molecules (gray box). Right panel, detailed view of the corresponding gray box. Side chains of ATF6α 61-76 are shown along with volume filling representation of the two Ceapin A7 molecules.

To gain mechanistic insight into how the ATF6α•ABCD3•Ceapin A7 complex assembles, we investigated its formation *in cellula* and *in vitro*. Using U2OS cells expressing GFP-ATF6α, we performed GFP pulldowns in the presence or absence of Ceapin A7 and immunoblotted for ABCD3 (**Fig. 1D**). Contrary to our previous studies^7^, we found that ATF6α coprecipitated ABCD3, even in the absence of Ceapin A7. In addition, we found that the ATF6α•ABCD3 interaction increased markedly upon Ceapin A7 addition, indicating that Ceapin A7 enhances (or stabilizes) a weak (or transient) interaction between ATF6α and endogenous ABCD3.

To understand how Ceapin A7 stabilizes the ATF6α•ABCD3 complex, we reconstituted the assembly *in vitro* with purified components. Briefly, we purified full-length, human ABCD3 for structural and biochemical studies (**Extended Data Fig. 2A**), and an MBP-tagged fragment of human ATF6α comprising residues 2-90 that, as previously shown, is necessary and sufficient for binding to ABCD3^7^. In line with the *in cellula* data, ABCD3 bound MBP-ATF6α^2–90^ in the absence of Ceapin A7 but addition of Ceapin A7 showed no further stabilizing effect (**Fig. 1E**). We reasoned that endogenous ABCD3 may dynamically cycle through distinct conformational states, only one of which is ATF6α-binding competent, while purified ABCD3 *in vitro* may preferentially populate a particular conformational state suitable for ATF6α binding. To test this possibility, we stabilized ABCD3 in its peroxisome-open conformation with orthovanadate^52^, mimicking the transition state of ATP hydrolysis. This treatment abolished ATF6α•ABCD3 binding, indicating that ATF6α binds specifically to the opposite, cytosol-open conformation of ABCD3 (**Fig. 1E, right lane**).

### Ceapin A7 directly binds ABCD3, stabilizing a pentameric complex with ATF6α

We next prepared ATF6α^2–90^•ABCD3•Ceapin A7 and ABCD3•Ceapin A7 complexes for single-particle cryo-EM, which reconstructs three-dimensional structures by averaging images of individual vitrified particles, and obtained 2.9 and 3.1 Å resolution structures of the respective complexes (**Fig. 1F, Extended Data Fig. 2B, 3A-B**, **Table 1**). The structures show ABCD3 as a two-fold symmetric homodimer that adopts a cytosol-open conformation, with no nucleotide bound to the NBDs. Within the cytosol-open solvated inner chamber formed by the transmembrane domain, we observed clear density for two Ceapin A7 molecules, bound symmetrically, one to each ABCD3 protomer, contacting transmembrane helices 4-6 and each burying a surface area of ∼660 Å^2^ (**Fig. 2A**). Ceapin A7 occupies a solvent-exposed binding pocket **(Fig. 2B)**, which extensively overlaps with that used for binding of the coenzyme A moiety of ABCD3 lipid transport substrates (**Fig. 2C**)^32,33^. However, we note that ABCD3 with Ceapin A7 bound in the substrate-binding pocket adopts a widened open angle, characteristic of the apo-state^33^ (PDB: 9NGM) rather than the substrate-bound state^32^ (PDB: 8Z9X) (**Extended Data Fig. 3C-D**).

**Figure 2:**
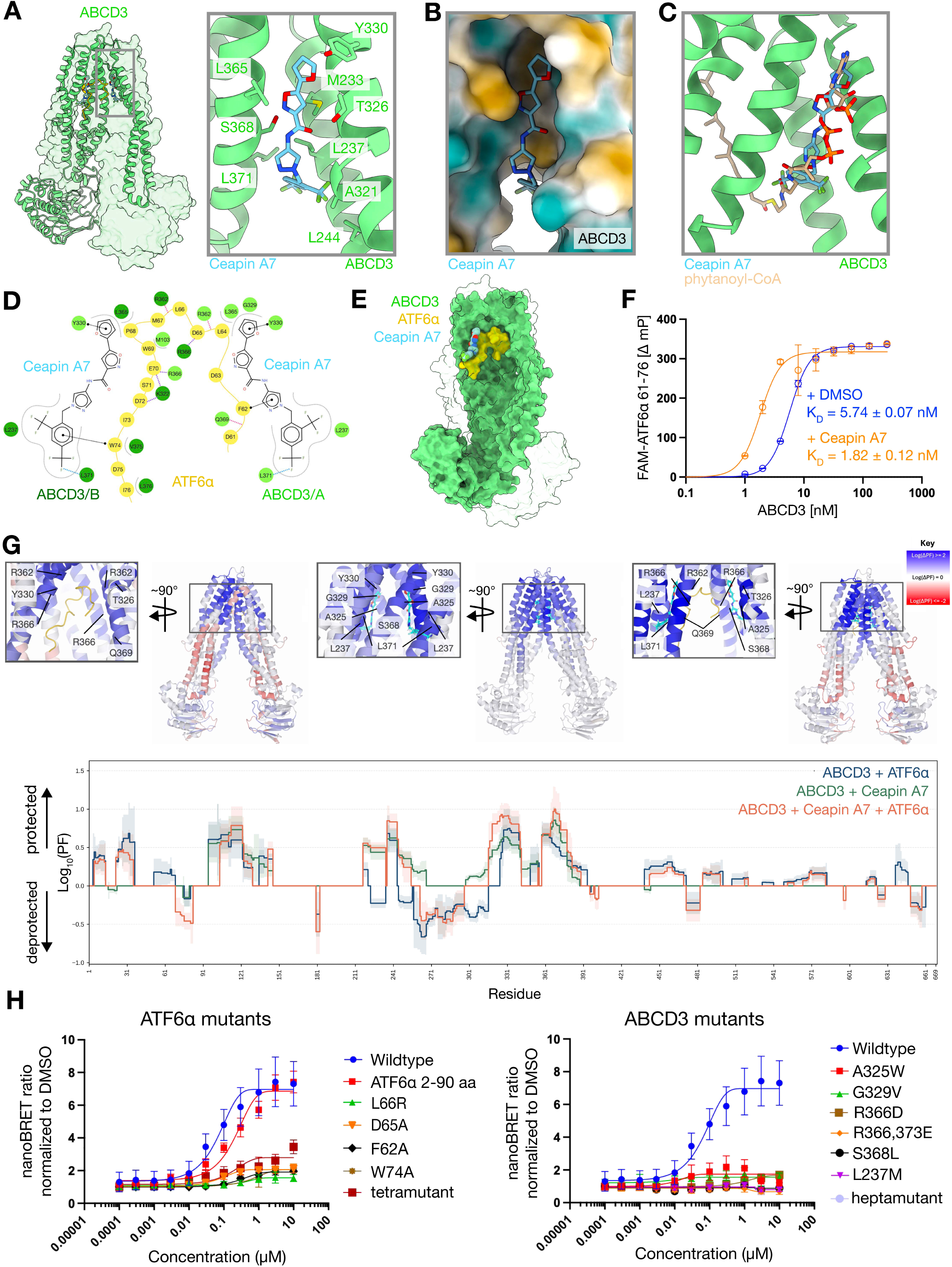
ATF6α directly binds ABCD3. **a**, ABCD3•ATF6α•Ceapin A7 structure with one ABCD3 protomer shown in cartoon representation (green) and one protomer shown in surface view (green, transparent). ATF6α (yellow) and Ceapin A7 (blue). Ceapin A7 binds in a pocket formed by 3 ABCD3 helices (gray box). Gray box shows a detailed view of one of the Ceapin A7 molecules bound to ABCD3. ABCD3 side chains interacting with Ceapin A7 are shown and labeled. **b**, Ceapin A7 binds within a solvated pocket in ABCD3. ABCD3 is shown in surface representation colored by hydrophobicity (yellow-more hydrophobic, blue-more hydrophilic). **c**, Ceapin A7 binding site overlaps with the ABCD3 substrate binding site. Position of the ABCD3 substrate, phytanoyl-CoA (tan) from PDB 8Z9X is overlayed onto the ABCD3•ATF6α•Ceapin A7 structure. Ceapin A7 overlaps with the CoA portion of phytanoyl-CoA. **d**, 2D interaction representation between each symmetrically bound Ceapin A7 molecule with ABCD3 promoter A (light green), ABCD3 promoter B (dark green), and ATF6α (yellow). Interactions are marked as follows: hydrophobic (dotted gray line), backbone halogen bond (dotted blue line), π-stacking (straight, dotted black line), salt bridge (dotted purple line), backbone hydrogen bond (dotted pink line). Adapted from PoseView and Protein-Ligand Interaction Profiler. **e**, ATF6α (yellow, surface representation) completely fills the space within the ABCD3 transmembrane cavity (one ABCD3 protomer, green, surface representation; other ABCD3 protomer transparent). A second ATF6α cannot be accommodated. **f**, Fluorescence polarization of 0.1 nM fluorescent ATF6α 61-76 peptide (FAM-ATF6α 61-76) binding to purified full-length ABCD3 *in vitro* with 25 uM Ceapin A7 (orange line, K_D_ = 1.82 ± 0.12 nM) or with DMSO vehicle (blue line, K_D_ = 5.74 ± 0.07 nM). Graph shows mean ± SD, n = 3 replicates per condition. **g**, HDX–MS analysis of the ABCD3-Ceapin A7-ATF6α complex used to identify changes in ABCD3 dynamics associated with ATF6α and Ceapin A7 binding. **Top**, differential protection factors are mapped onto the ABCD3 homodimer for ABCD3 in the presence of ATF6α, Ceapin A7, or both Ceapin A7 and ATF6α. Side views and corresponding views rotated by 90° highlight the transmembrane cavity and the proposed ternary-complex interface. The ATF6α segment and Ceapin A7 are shown in yellow and cyan, respectively, and selected ABCD3 residues within the affected region are labeled. Structural coloring represents the direction and magnitude of the protection factor change according to the blue–white–red scale shown, with white indicating no detectable change. Bottom, residue-resolved protection factor profiles are shown for ABCD3 plus ATF6α (blue), ABCD3 plus Ceapin A7 (green), and the ATF6α•ABCD3•Ceapin A7 ternary complex (orange), relative to ABCD3 alone. Lines show significant weighted log10 protection factors. Shaded areas indicate ± weighted log10(PF) uncertainty calculated from the total weighted variance of contributing peptides. **h**, NanoBRET validation of the ATF6α–ABCD3 interaction interface. Ceapin A7-dependent proximity between ATF6α and ABCD3 was measured using NanoBRET, with the NanoBRET ratio normalized to the DMSO vehicle control. Left, concentration–response curves are shown for wild-type ATF6α, the ATF6α(2–90) fragment, the individual ATF6α mutants, and the combined tetramutant, each tested with wild-type ABCD3. Wild-type ATF6α and ATF6α(2– 90) produced a concentration-dependent increase in NanoBRET signal, whereas the individual interface mutations and tetramutant reduced the maximal response. Right, corresponding concentration–response curves are shown for wild-type ABCD3, the indicated ABCD3 interface mutants, and the combined hexamutant tested with wild-type ATF6α. Mutations in the ABCD3 transmembrane interface strongly attenuated or abolished the Ceapin A7-dependent NanoBRET response. Symbols represent mean ± SEM from 4 independent experiments, and solid lines represent nonlinear concentration–response fits.

The ATF6α^2–90^•ABCD3•Ceapin A7 and the ABCD3•Ceapin A7 structures are nearly identical (RMSD of 1.32 Å, Ceapin A7•ABCD3 Cα 57-650), apart from the extra density for ATF6α, which we observed sandwiched between the two Ceapin A7 molecules (**Fig. 1F**). We assigned this density to a single loop of ATF6α amino acids 61-76, supported by AlphaFold modeling, map features, conservation, peptide-binding data, and structure-guided mutagenesis (see **Methods, Extended Data Fig. 4**). The resolved ATF6α(61-76) peptide contacts both ABCD3 protomers and both Ceapins, burying a surface area of ∼1300 Å^2^, and resulting in a pentameric complex (ATF6α_1_•ABCD3_2_•Ceapin A7_2_) in which ATF6α breaks the two-fold symmetry (**Fig. 2D**). ATF6α(61-76) makes a tight turn to fit into the ‘ceiling’ of the ABCD3 chamber, making contacts with the chamber’s top and sides through hydrophobic and hydrophilic interactions. The dimensions of the chamber are incompatible with accommodation of a second ATF6α peptide in a similar binding mode (**Fig. 2E**). ATF6α makes multiple backbone intra-molecular hydrogen bonds, stabilizing the tight turn (**Extended Data Fig. 5A**). The hydrophobic interactions consist of ATF6α L66, which sits at the apex of the ABCD3 chamber in between the two ABCD3 protomers, and ATF6α M67 and W69, which occupy a hydrophobic pocket within the ABCD3 chamber (**Extended Data Fig. 5B**). The remainder of the ATF6α loop is predominantly negatively charged and complements the positively charged inner chamber of ABCD3, stabilized by multiple salt bridges (**Extended Data Fig. 5C,D**). We validated the existence of two of the salt bridges (ABCD3(R366)-ATF6α(D65) and ABCD3(R366)-ATF6α(E70)) by charge-swapping, mutating ABCD3(R366) to glutamate and ATF6α(D65 and E70) to arginine. We used the formation of GFP-ATF6α puncta in cells expressing these mutants to assess ABCD3-ATF6α binding. We observed that ABCD3^R366E^ did not bind wild-type ATF6α and ATF6α^D65R,E70R^ did not bind wild-type ABCD3, but co-expression of both charge-swapped mutants to restore the salt bridge rescued binding, validating our structural model (**Extended Data Fig. 5E**). Finally, we quantitatively assessed the binding of this region of ATF6α to ABCD3 using a synthesized fluorescent peptide, FAM-ATF6α(61-76). The ATF6α peptide bound ABCD3 with high affinity as assessed by fluorescence polarization (K_D_ = 5.74 ± 0.07 nM), and as expected, the active analog, Ceapin A7, improved binding (by ∼3-fold; K_D_ = 1.82 ± 0.12 nM) (**Fig. 2F**).

Hydrogen/deuterium exchange followed by mass spectrometry (HDX-MS; slower deuterium uptake indicates buried/rigid regions, whereas faster uptake indicates increased exposure/flexibility) provided orthogonal support for the cryo-EM model of the ATF6α•ABCD3•Ceapin A7 pentameric complex (**Fig. 2G**). Relative to apo-ABCD3, ATF6α binding induced broad, bidirectional HDX changes across ABCD3, including strong protection in residues ∼20–30, ∼91–120, ∼230–240, ∼331–350, and ∼361–371. These protected regions are consistent with contacts observed in the cryo-EM model and support engagement of ATF6α within the ABCD3 transmembrane cavity. Additionally, increased exchange was observed over an extended region of the transmembrane domain, approximately spanning residues 240–320, suggesting ATF6α binding also remodels ABCD3, possibly by promoting local helix destabilization or transient opening of the cavity. This is consistent with the cryo-EM structure, in which only a short ATF6α segment is resolved at the apex of the transmembrane cavity, while the remaining disordered region is expected to occupy the chamber.

Unlike ATF6α, Ceapin A7 alone produced predominantly protective effects over a more compact region of ABCD3 centered on the compound/substrate-binding pocket, with the largest protection factors peaking in regions approximating those observed for ATF6α. Thus, ATF6α and Ceapin A7 affect overlapping regions of the ABCD3 cavity, but with distinct long-range consequences. When ATF6α and Ceapin A7 were both present, ATF6α-associated deprotection was reduced and the HDX pattern shifted toward protection, consistent with Ceapin A7 stabilizing the ATF6α-engaged, cytosol-open conformation of ABCD3. Comparison of the pentameric complex with the sum of the individual ATF6α- and Ceapin A7-bound states revealed non-additive behavior, including regions of apparent potentiation as well as regions in which the individual effects were not retained in the combined state. Together, these data support a molecular-glue model in which ATF6α has basal affinity for ABCD3, while Ceapin A7 reinforces interactions within the substrate-binding pocket to stabilize the ATF6α-ABCD3 interaction.

To quantify Ceapin A7-dependent complex formation and monitor the association between ABCD3 and ATF6α, we established a live-cell nanoBRET (bioluminescence resonance energy transfer) assay, which reports drug-induced nanoscale proximity between protein partners as a surrogate for pentameric complex assembly (**Fig. 2H**). Ceapin A7 produced a clear dose-dependent increase in nanoBRET signal for the wild-type ABCD3-ATF6α pair, both with full-length ATF6α and with the ATF6α(2-90) fragment, indicating that the N-terminal segment is sufficient to support ligand-induced binding. In contrast, mutations in ATF6α (F62A, L66R, D65A, W74A) or in ABCD3 (L237M, A325W, G239V, R366D, R366E/R373E, S368L) that disrupt the interface defined by the cryo-EM structures strongly attenuated or eliminated the nanoBRET signal. This loss is consistent with a failure to form or stabilize the pentameric complex and mirrors the loss of Ceapin A7-dependent GFP-ATF6α puncta formation observed with the same ABCD3 mutants (**Extended Data Fig. 5F**). We note that the related peroxisome lipid transporters, ABCD1 and ABCD2, contain a tryptophan at position 325, which based on our structure is predicted to clash with Ceapin A7 (**Extended Data Fig. 5G**). The observed lack of GFP-ATF6α puncta formation with the ABCD3(A325W) mutant confirms this prediction and suggests that Ceapin A7 binding is specific for ABCD3.

### Ceapin A7 inhibits ABCD3 in a substrate-competitive mode

As a complementary assay to quantitatively assess binding between Ceapin A7 and ABCD3, we developed a fluorescence polarization anisotropy binding assay, which tracks the binding of a small fluorescent ligand to a larger protein, using a fluorescent Ceapin A7 analog, FAM-Ceapin A7 (**Fig. 3A**). FAM-Ceapin A7 bound ABCD3 with a K_D_ of 168 ± 8.40 nM, confirming that Ceapin A7 binds ABCD3 in the absence of ATF6α, and with a Hill coefficient of 1.18, indicating slight cooperativity. The apparent K_D_ falls in the same range as the cellular IC_50_ reported for Ceapin A7 previously (167 versus 600 nM, respectively)^6^. In the structure, the ATF6α density mostly contacts ABCD3 with limited interactions with the two Ceapin A7 molecules. Consistent with this, we failed to detect measurable affinity between FAM-Ceapin A7 and purified ATF6α 2-90 **(Extended Data Fig. 5H**).

**Figure 3:**
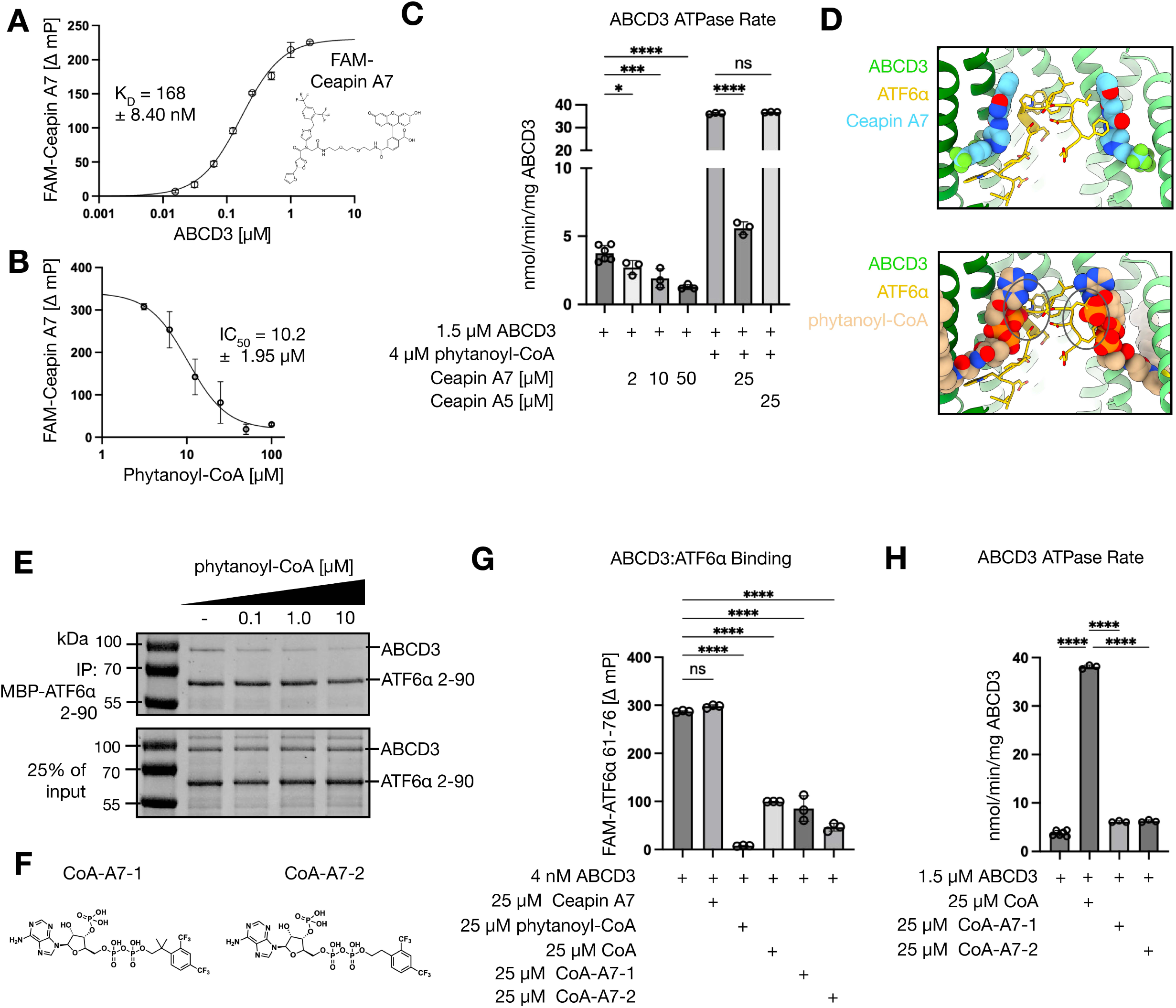
Ceapin A7 is a substrate-competitive inhibitor of ABCD3. **a**, Fluorescence polarization of 10 nM fluorescent Ceapin A7 (FAM-Ceapin A7) binding to purified full-length ABCD3 *in vitro* (K_D_ = 168 ± 8.40 nM). Graph shows mean ± SD, n = 3 replicates per condition. **b**, Fluorescence polarization of 10 nM fluorescent Ceapin A7 (FAM-Ceapin A7) binding to 1 μM purified full-length ABCD3 *in vitro* with titration of ABCD3 substrate, phytanoyl-CoA (IC_50_ = 10.2 ± 1.95 μM). Graph shows mean ± SD, n = 3 replicates per condition. **c**, Measurement of ATPase rate for purified full-length ABCD3 *in vitro* with addition of ABCD3 substrate (phytanoyl-CoA), Ceapin A7, or an inactive Ceapin analog, Ceapin A5, as a control. Graph shows mean ± SD, n = 3-6 replicates per condition. Statistical significance was evaluated using a one-way ANOVA with *post-hoc* Bartlett’s test. \**P* ≤ 0.05, \*\**P* ≤ 0.01, \*\*\**P* ≤ 0.001, \*\*\*\**P* ≤ 0.0001. **d**, Top panel, view inside the ABCD3 chamber of the ATF6α• ABCD3•Ceapin A7 structure. ABCD3 promoter A (light green), ABCD3 promoter B (dark green), ATF6α (yellow), and Ceapin A7 (blue). Side chains of ATF6α 61-76 are shown along with volume filling representation of the two Ceapin A7 molecules. ATF6α 61-76 makes interactions with both Ceapin A7 molecules. Bottom panel, ABCD3 substrate, phytanoyl-CoA (tan) from PDB 8Z9X overlayed onto the ATF6α• ABCD3•Ceapin A7 structure. Phytanoyl-CoA clashes at multiple points with ATF6α (black circles). **e**, Coomassie-stained SDS-PAGE from *in vitro* immunoprecipitation of 2 μM purified MBP-ATF6α 2-90 with 2 μM purified ABCD3 in the presence of increasing phytanoyl-CoA concentrations. **f**, Custom synthesized coenzyme A (CoA)-Ceapin A7 (A7) hybrid molecules. **g**, Fluorescence polarization of 0.1 nM fluorescent ATF6α 61-76 peptide (FAM-ATF6α 61-76) binding to 4 nM purified full-length ABCD3 *in vitro* with 25 uM Ceapin A7, phytanoyl-CoA (ABCD3 substrate), CoA, CoA-A7-1, or CoA-A7-2. Graph shows mean ± SD, n = 3 replicates per condition. Statistical significance was evaluated using a one-way ANOVA with *post-hoc* Dunnett’s test. \**P* ≤ 0.05, \*\**P* ≤ 0.01, \*\*\**P* ≤ 0.001, \*\*\*\**P* ≤ 0.0001. **h**, Measurement of ATPase rate for purified full-length ABCD3 *in vitro* with addition of 25 uM ABCD3 CoA, CoA-A7-1, or CoA-A7-2. Graph shows mean ± SD, n = 3-6 replicates per condition. Statistical significance was evaluated using a one-way ANOVA with *post-hoc* Sidak’s test. \**P* ≤ 0.05, \*\**P* ≤ 0.01, \*\*\**P* ≤ 0.001, \*\*\*\**P* ≤ 0.0001.

We next added increasing concentrations of the ABCD3 transport substrate phytanoyl-CoA to the FAM-Ceapin A7•ABCD3 binding assay. Phytanoyl-CoA competed with Ceapin A7 (IC_50_ = 10.2 ± 1.95 μM) (**Fig. 3B**). Thus, binding of Ceapin A7 and substrate by ABCD3 are mutually exclusive events, as predicted by our structures. Furthermore, addition of orthovanadate, which stabilizes ABCD3 in its peroxisome-open conformation^52^, fully inhibited FAM-Ceapin A7 binding to ABCD3 (**Extended Data Fig. 5I**). We conclude that Ceapin A7 binds selectively to the cytosol-open conformation, matching the widened, apo-conformation of ABCD3 that ATF6α binds with high affinity.

Ceapin A7 binds in the substrate-binding pocket yet does not induce the narrowed cytosol-open conformation, as a transport substrate does. We therefore asked whether Ceapin A7 and transport substrates affect ABCD3’s ATPase rate similarly or distinctly (**Fig. 3C**). Consistent with previous reports^33^, purified ABCD3 displayed a low basal ATP hydrolysis rate (3.75 ± 0.23 nmol ADP/min/mg) that was stimulated ∼10-fold by phytanoyl-CoA addition of (36.3 ± 0.2 nmolADP/min/mg). By contrast, addition of Ceapin A7 inhibited both the basal ATP hydrolysis and its substrate-dependent stimulation. The inactive Ceapin analog, Ceapin A5, which lacks the phenyl-trifluoromethyl group^6^, had no effect. Thus, despite occupying the same binding site, Ceapin A7 and the transport substrate phytanoyl-CoA have the opposite effect on ABCD3’s ATP hydrolysis rate. We reason that Ceapin A7 stabilizes the widened cytosol-open apo-conformation, which impedes the association of ABCD3’s NBDs, a prerequisite for ATP hydrolysis.

Docking the phytanoyl-CoA•ABCD3 structure onto the ATF6α•ABCD3•Ceapin A7 structure revealed that the bulky coenzyme A phosphate groups of both symmetrically bound transport substrates would sterically clash with ATF6α (**Fig. 3D**), suggesting that ABCD3 transport substrates inhibit ATF6α binding to ABCD3. To test this notion, we titrated the phytanoyl-CoA concentration and assessed ATF6α•ABCD3 complex formation in our *in vitro* pull-down assay. Indeed, phytanoyl-CoA addition displaced ATF6α from ABCD3 in a concentration dependent manner (**Fig. 3E**). Thus, Ceapin A7 stabilizes ATF6α binding to ABCD3 i) by inhibiting binding of a competing fatty acyl-CoA transport substrate, and ii) by contributing to the binding energy through direct contacts with ATF6α.

Given the opposing effects of Ceapin A7 and transport substrates on ABCD3’s ATP hydrolysis rate, we asked which ligand features distinguish substrate-induced ATPase activation from Ceapin A7-mediated inhibition. We reasoned the phenyl-trifluoromethyl group of Ceapin A7 accounts for its ability to stabilize the apo-like conformation of ABCD3, given the lack of ATPase inhibition by Ceapin A5, analysis of the Ceapin A7•ABCD3 interaction, and the finding this group is essential for Ceapin A7’s potency^6^. We tested the properties of two custom-synthesized hybrid compounds (“CoA-A7-1 and CoA-A7-2”), in which we appended the phenyl-trifluoromethyl group to the end of coenzyme A (**Fig. 3F**). Applying the fluorescence polarization binding assay with FAM-ATF6α(61-76), we assessed how the hybrid compounds affect complex formation. Addition of phytanoyl-CoA completely abolished binding, while CoA alone significantly reduced binding, both as expected from the predicted steric clashes of ATF6α with CoA’s bulky phosphate groups. Addition of CoA-A7-1 or CoA-A7-2 also reduced ATF6α-ABCD3 binding significantly, to a slightly greater extent than CoA alone (**Fig. 3G**), demonstrating that the hybrid molecules have the same disruptive effect as CoA on ATF6α-ABCD3 binding. By contrast, in the ATPase assay, CoA-A7-1 or CoA-A7-2 marginally potentiated hydrolysis (∼1.5-fold), compared to the more pronounced inhibition observed for CoA (∼10-fold) (**Fig. 3H**). These data are consistent with the possibility that Ceapin A7’s phenyl-trifluoromethyl group blocks ABCD3 from reaching its ATPase-active conformation and that the phosphate groups on CoA inhibit ATF6α binding to ABCD3.

### Endogenous ATF6α•ABCD3 complexes generate dynamic ER–peroxisome contact sites, stabilized upon Ceapin A7 treatment

Having defined the structural basis of the ATF6α•ABCD3 interaction and its stabilization by Ceapin A7, we next aimed to decipher the functional significance of this interaction. We asked whether ATF6α, expressed at physiological levels, associates with peroxisomes. To this end, we engineered U2OS cells in which endogenous ATF6α, expressed from its own promoter, was marked at its N-terminus with a Halo tag. Consistent with the findings for overexpressed GFP-ATF6α (**Fig. 1D)**, immunoprecipitation of Halo-ATF6α revealed a basal interaction with ABCD3, which was significantly enhanced by treatment with Ceapin A7 (**Fig. 4A**). We next labeled peroxisomes with a fluorescent lipid (Peroxi-SPY555). Halo-ATF6α displayed a predominantly ER-like distribution, with ∼6% of the protein forming puncta. A fraction of these puncta (∼33±2%, i.e. ∼2% of the total Halo-ATF6) colocalized with peroxisomes. Upon Ceapin A7 treatment, the abundance of ATF6α puncta markedly increased (51.4% of total Halo-ATF6α), and more than half of the puncta (∼62±2%) overlapped with peroxisomes (**Fig. 4B and 4C**). Immunostaining for the peroxisomal marker PEX14 similarly revealed minimal colocalization of endogenous Halo-ATF6α with PEX14 under basal conditions, which however was markedly enhanced upon Ceapin A7 treatment. Quantitative analysis determining the Manders overlap coefficient (M1) confirmed that the fraction of Halo-ATF6α puncta associated with peroxisomes increased from 20% under basal conditions to 67% following Ceapin A7 treatment **(Extended Data Fig. 6A)**. These results show that endogenous ATF6α expressed at physiological levels can associate with peroxisomes and that Ceapin A7 substantially enhances this interaction.

**Figure 4:**
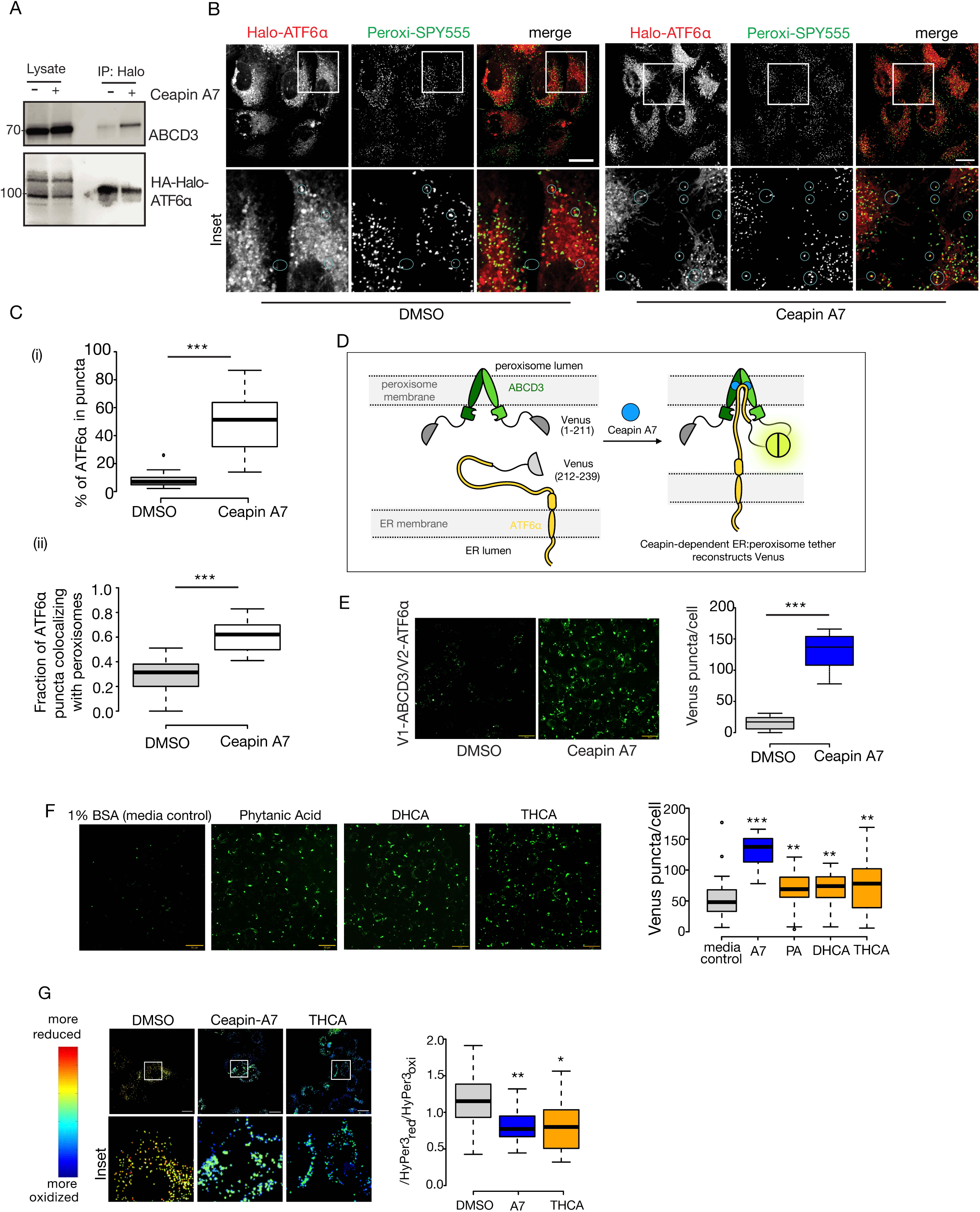
ATF6α–ABCD3 ER–peroxisome tethers are stabilized by Ceapin A7 and peroxisomal lipid substrates. **a**, Immunoprecipitation of endogenously tagged 3×HA–Halo–ATF6α from U2OS cells treated with DMSO or Ceapin A7, followed by immunoblotting for ABCD3. ATF6α is tagged at its endogenous locus, and Ceapin A7 enhances the interaction between ATF6α and ABCD3. **b**, Representative fluorescence images of endogenously tagged 3×HA–Halo–ATF6α labeled with the Halo ligand JFX650 (red) and peroxisomes labeled with Peroxi-SPY555, in cells treated with DMSO or 10μM Ceapin A7 for 4 h. Insets highlight ATF6α puncta colocalizing with peroxisomes. Scale bar, 5 μm. **c**, Quantification of the experiment in **b**: (i) the fraction of total 3×HA–Halo–ATF6α forming puncta and (ii) the fraction of ATF6α puncta overlapping with peroxisomes under DMSO and Ceapin A7 treatment. Center lines indicate medians; box limits represent the 25th and 75th percentiles; whiskers extend 1.5× the interquartile range; outliers are shown as individual points. *n* = 47 (DMSO) and 52 (Ceapin A7) cells from 3 independent experiments. Statistical significance was determined using a two-tailed Student’s *t*-test. \*\*\**P* < 0.001. **d**, Schematic of the split-Venus assay used to detect ATF6α–ABCD3 interactions at ER– pero xisome contact sites. Venus fluorescence is reconstituted upon tether formation. **e**, Representative images of split-Venus fluorescence in cells expressing ATF6α fused to Venus(212–239) and ABCD3 fused to Venus(1–211) under the indicated conditions, demonstrating increased puncta upon 10μM Ceapin A7 treatment for 4 h. Quantification of puncta per cell is shown at right. Center lines indicate medians; box limits represent the 25th and 75th percentiles; whiskers extend 1.5× the interquartile range; outliers are shown as individual points. *n* = 65 cells per condition from 3 independent experiments. Statistical significance was determined using a two-tailed Student’s *t*-test. \*\*\**P* < 0.001. Scale bar, 10 μm. **f**, Quantification of Venus puncta per cell following treatment with10μM peroxisomal lipid substrates (phytanic acid, DHCA, THCA or media control (1% BSA/DMEM) for 4 h, showing that substrate engagement enhances ATF6α–ABCD3 interaction. Center lines indicate medians; box limits represent the 25th and 75th percentiles; whiskers extend 1.5× the interquartile range; outliers are shown as individual points. *n* = 48,65, 62,62 and 48 cells for media control, Ceapin A7, phytanic acid (PA), DHCA, and THCA treatment respectively. Data represents 3 independent experiments. Statistical significance was determined using a two-tailed Student’s *t*-test. **P ≤ 0.01, \*\*\**P* < 0.001. Scale bar, 10 μm. **g**, Representative ratiometric images of peroxisomal HyPer3 in cells treated with the 10 μM Ceapin A7 or 10 μM THCA for 16 h. HyPer3 fluorescence was acquired at dual excitation wavelengths and converted to a ratio (F500/F420), which was pseudo-colored using a rainbow lookup table in Fiji, where blue indicates a more oxidized state. Quantification of peroxisomal redox status is shown at right. Center lines indicate medians; box limits represent the 25th and 75th percentiles; whiskers extend 1.5× the interquartile range; outliers are shown as individual points. *n* = 50 cells per condition. Statistical significance was determined using a two-tailed Student’s *t*-test. \**P* < 0.05 / \*\**P* < 0.01. Scale bar, 5 μm.

### Peroxisomal lipids promote the association of ATF6α with ABCD3

To identify physiological conditions that may modulate the ATF6α•ABCD3 interaction, we adapted the Split-Venus fluorescence complementation system^53^, in which two non-fluorescent fragments of the Venus protein reconstitute and fluoresce only when brought together by their fused binding partners. We tagged the C-terminus of ABCD3 with the long Venus fragment (amino acids 1–211) and the N-terminus of ATF6α with the short fragment (amino acids 212– 239). In this system, conditions that promote ATF6α•ABCD3 engagement should exhibit Venus fluorescence (**Fig. 4D**). Because Venus complementation stabilizes these proximity events, the assay reports cumulative ATF6α•ABCD3 encounters rather than native dwell time. As expected, control cells expressing only one of the constructs showed no detectable fluorescence (**Extended Data Fig. 6B**). By contrast, co-expression of both fragments produced fluorescent puncta, the number of which was significantly enhanced by Ceapin A7 treatment (**Fig. 4E**).

To test which conditions might affect the association between ATF6α and ABCD3, we first exposed cells to the ER stressors—tunicamycin, thapsigargin, and DTT—all of which activate the three branches of the UPR, including ATF6α. Surprisingly, these ER stressors differentially affected the number of Venus puncta: tunicamycin and thapsigargin had no effect, whereas DTT treatment markedly increased puncta numbers. Thus, general UPR activation is not sufficient to promote ATF6α•ABCD3 proximity; the DTT response may instead reflect redox perturbation or another DTT-specific effect. We next tested stressors that disrupt lipid metabolism. Treatment with palmitic acid (a saturated fatty acid that induces ER stress) or oleic acid (which promotes lipid droplet formation) did not alter Venus puncta numbers^54,55^. Similarly, mitochondrial inhibitors (FCCP and oligomycin) and oxidative stress induced by H₂O₂ failed to change the abundance of fluorescent puncta (**Extended Data Fig. 6C, D**).

By contrast, treatment of cells with known ABCD3 transport substrates—including the bile acid biosynthetic intermediates trihydroxy-cholestanoic acid (THCA) and dihydroxy-cholestanoic acid (DHCA)^56^, as well as the branched-chain fatty acid phytanic acid^32^—robustly increased the number of Venus puncta from a mean value of 15 ± 1 per cell under basal conditions to 69 ± 3, 75 ± 6, and 68 ± 3, respectively (**Fig. 4F**). THCA and DHCA undergo peroxisomal β-oxidation to generate mature bile acids, whereas phytanic acid is metabolized via peroxisomal α-oxidation followed by β-oxidation. Their divergent fates after import suggest that it is the engagement of ABCD3 in active transport that promotes formation of the ATF6α•ABCD3 complex, i.e., peroxisomal lipid import activity controls the ER-peroxisome junction mediated by ATF6α and ABCD3. This effect was specific to peroxisomal substrate transport, as other cell stressors did not increase Venus puncta above basal levels, indicating that generic cell stress is not sufficient to drive ATF6α•ABCD3 complex formation. Peroxisomal lipid import activity is therefore a direct regulatory input into the ER–peroxisome junction formed by ATF6α and ABCD3.

### Ceapin A7 and peroxisomal lipids increase peroxisomal ROS

One consequence of lipid import into the peroxisome is lipid oxidation, which generates hydrogen peroxide and increases oxidative burden within the organelle^22^. To evaluate peroxisome redox state, we used a ratiometric fluorescent reporter, HyPer3 (a genetically encoded hydrogen peroxide sensor), directed to peroxisomes via an SKL targeting sequence^57^. Because hydrogen peroxide is one of the end products of peroxisome-specific β-oxidation specific (in contrast to mitochondria, which produce water)^58^, its relative abundance within the peroxisomal lumen serves as a functional proxy for peroxisomal metabolic activity. HyPer3 detects local hydrogen peroxide levels by shifting its excitation spectrum between oxidized and reduced states. Accordingly, the ratio of fluorescence intensities at 500 nm versus 420 nm excitation (F500/F420) provides a quantitative measure of peroxisomal redox status and overall peroxisomal health^59^. Cells treated with Ceapin A7 or THCA displayed a significantly higher proportion of peroxisomes with elevated oxidative burden compared to control cells (**Fig. 4G**; p = 1.85 x 10^-6^, and 0.0035 for Ceapin A7 and THCA, respectively), indicative of peroxisomal dysfunction and consistent with our observations of peroxisome enlargement by cryo-ET after Ceapin A7 treatment (see **Extended Data Fig. 1C**).

### The autophagy adaptor protein p62 is recruited to ATF6α•ABCD3–mediated ER–peroxisome tether sites

One possible fate of damaged peroxisomes is turnover by autophagy. To test this possibility, we performed mass spectrometric analysis of the ATF6α•ABCD3 interactome to ask whether ATF6α•ABCD3 contacts recruit the autophagy machinery. Briefly, we immunoprecipitated the complex using anti-GFP nanobodies that recognize the reconstituted Venus protein generated by Split-Venus complementation. Mass spectrometry revealed a notable enrichment of proteins involved in actin remodeling (**Fig. 5A**). Consistent with this, the tomograms of Ceapin A7-treated cells showed a remarkable abundance of actin fibers at the tether sites (**Fig. 1C**). Disruption of actin polymerization with CK-666 had no effect on puncta formation (**Extended Data Fig. 7A**). The role of actin at ATF6α•ABCD3 tether sites remains unexplored.

**Figure 5:**
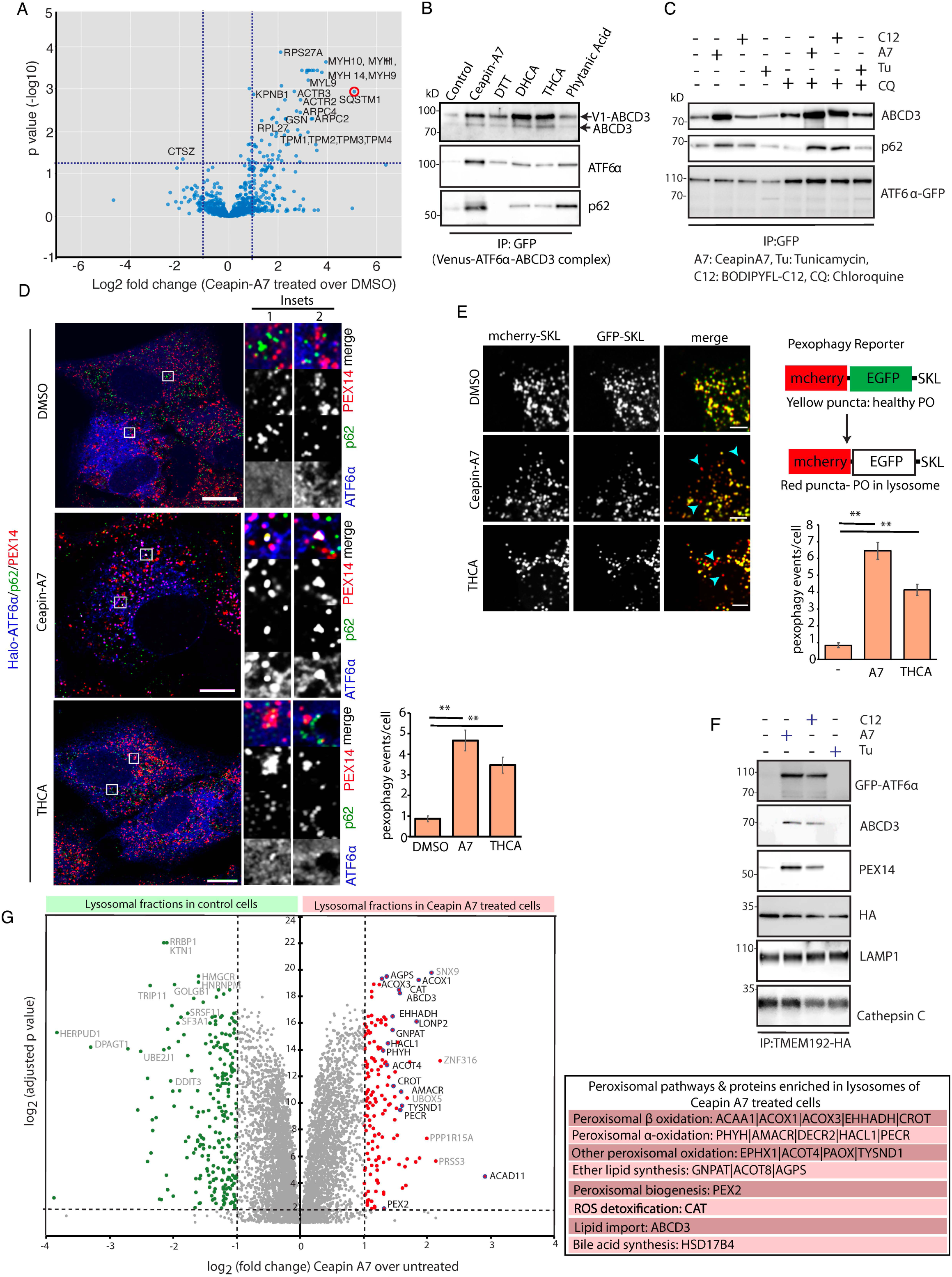
ATF6α–ABCD3 tether sites recruit p62 and undergo pexophagy. **a**, Mass spectrometric analysis of the ATF6α•ABCD3 interactome following immunoprecipitation of the split-Venus complex from cells treated with 10 μM Ceapin A7 for 4 h, revealing enrichment of interacting proteins, including the autophagy adaptor p62. **b**, Immunoblot analysis following immunoprecipitation of the reconstituted Venus-based ATF6α– ABCD3 complex showing association with ABCD3, ATF6α, and p62 under the indicated treatments. Cells were treated with 10 μM Ceapin A7, 10 μM lipid substrates for 4 h, or 1 mM DTT for 1 h. **c**, Immunoblot analysis of GFP pulldowns of ATF6α from cells expressing GFP– ATF6α, showing association with ABCD3 and p62 under the indicated treatments. Cells were treated with 10 μM Ceapin A7, 2.5 μg/mL tunicamycin, or 1 μM BODIPY-C12 for 6 h, in the presence or absence of 50 μM chloroquine (CQ), which was added to block autophagic– lysosomal degradation. Blocking lysosomal degradation enhances the association of p62 with ATF6α. **d**, Representative immunofluorescence images of cells expressing 3×HA–Halo–ATF6α (labeled with JFx650, red) and stained for p62 and the peroxisomal membrane protein PEX14, under the indicated treatments. Cells were treated with 10 μM Ceapin A7 or 10 μM lipid substrates for 16 h. Colocalization of p62 with PEX14 is observed, indicating recruitment of the autophagy adaptor to peroxisomes. Quantification of this pexophagy event is shown at right. Data are presented as mean ± s.e.m.; *n* = 30 cells from 3 independent experiments. Scale bar, 5μm. **e**, Tandem fluorescent pexophagy reporter (GFP–SKL/mCherry–SKL). Red puncta indicate peroxisomes delivered to lysosomes, while yellow puncta represent healthy peroxisomes. Quantification of red puncta shows increased pexophagy upon treatment with 10 μM Ceapin A7 or 10 μM THCA for 16 h. Data are presented as mean ± s.e.m.; *n* = 30 cells from 3 independent experiments. Scale bar, 5μm **f**, Immunoblot analysis of lysosomal fractions isolated by Lyso-IP from cells expressing 3×HA–TMEM192, showing accumulation of ATF6α, ABCD3, and PEX14 in lysosomes under conditions that promote tether formation. Cells were treated with 10 μM Ceapin A7, 1 μM BODIPY-labeled peroxisomal lipid, or 2.5 μg/mL tunicamycin for 6 h. LAMP1 and cathepsin are shown as lysosomal membrane and lumenal markers, respectively, and serve as loading controls. **g**, Volcano plot of proteins identified by mass spectrometry in lysosomes isolated by Lyso-IP from cells expressing 3×HA–TMEM192, comparing control and 10 μM Ceapin A7 treatment for 16 h. Intact lysosomes were immunopurified using anti-HA beads. Proteins enriched upon Ceapin A7 treatment include peroxisomal proteins indicating increased delivery of peroxisomal material to lysosomes. Dashed lines indicate fold-change and statistical significance cutoffs.

In addition to the local engagement of the actin cytoskeleton, the mass spectrometry data indicated a ∼30-fold enrichment of the autophagy adaptor SQSTM1/p62 in Ceapin A7–treated cells as compared to controls (**Fig. 5A**). We validated the association of p62 with the Venus– ATF6α•ABCD3 complex by immunoblotting following immunoprecipitation (**Fig. 5B**). Because p62 links ubiquitylated cargo to selective autophagy pathways, we asked whether blocking lysosomal degradation would influence its association with ATF6α. We first treated GFP-ATF6α– expressing U2OS cells with chloroquine to inhibit autophagic flux, and then immunoprecipitated GFP-ATF6α and immunoblotted for ABCD3 and p62 (**Fig. 5C**). Blocking lysosomal degradation in Ceapin A7–treated cells markedly increased the interaction between ATF6α and p62, consistent with p62 accumulation at ER–peroxisome tether sites. Treating cells with fluorescent BODIPY-C12—a cell permeable fatty acid analog transported into peroxisomes in an ABCD3-dependent manner^60^—also increased the interaction between ATF6α, ABCD3, and p62, supporting a link between ABCD3 mediated lipid-substrate import and p62 recruitment to the ATF6α–ABCD3 complex when peroxisomes are challenged with fatty acid overload. Consistent with the results above (**Extended Data Fig. 6D**), the ER stressor tunicamycin did not enhance this association (**Fig. 5C**).

### Ceapin A7 and peroxisomal lipids promote enhanced pexophagy

Immunofluorescence staining of Halo-ATF6α-expressing U2OS cells treated with Ceapin A7 or THCA using antibodies for p62 and the peroxisomal membrane protein PEX14, revealed strong colocalization between the two proteins (**Fig. 5D, Extended Data Fig. 7B**). To monitor lysosomal delivery of peroxisomal cargo after Ceapin A7 or THCA treatment, we employed a tandem fluorescent reporter^61^, in which GFP and mCherry were fused to the peroxisomal targeting sequence SKL. Under neutral pH, such as within healthy peroxisomes or autophagosomes, both reporter modules fluoresce, producing yellow puncta when visualized by confocal microscopy. By contrast, when peroxisomes reach acidic compartments, such as the lysosome, GFP fluorescence is quenched while mCherry persists, causing these structures to appear as red puncta. Cells treated with Ceapin A7 or peroxisomal lipid substrates exhibited numerous red puncta, consistent with increased pexophagy-associated lysosomal delivery (**Fig. 5E, Extended Data Fig. 7C**).

As complementary test for lysosomal delivery, we enriched lysosomes from cells expressing 3×HA-TMEM192, a lysosomal membrane protein, using organellar immunoprecipitation (Lyso-IP)^62^. This approach isolates intact lysosomes using HA-tagged magnetic beads: we confirmed successful enrichment by the presenece of the lysosomal membrane protein LAMP1 and the lumenal enzyme Cathepsin C (**Extended Data Fig. 8A**). Lysosomal fractions from cells treated with Ceapin A7 or peroxisomal lipid contained ATF6α, ABCD3, and PEX14, supporting incorporation of ATF6α and peroxisomal membrane proteins into lysosomal compartments under tether-stabilizing conditions (**Fig. 5F**). Mass spectrometric analysis of lysosomes showed enrichment of peroxisomal proteins upon Ceapin A7 treatment (**Fig. 5G**). These included ABCD3, as well as proteins of the peroxisomal β-oxidation and ɑ-oxidation pathways, plasmalogen (ether lipid) synthesis, and catalase, which detoxifies peroxisomal ROS. GO term analysis further identified peroxisomal processes as the most enriched pathways in Ceapin A7-treated cells (**Extended Data Fig. 8B**).

### p62 recruitment and pexophagy-driven peroxisome loss require both ATF6α and ABCD3

To determine whether the recruitment of p62 to ATF6α•ABCD3 tether sites requires the assembled complex, we examined its interaction with p62 in dCas9-U2OS cells transduced with sgRNAs targeting either ATF6α or ABCD3. Depletion of either ATF6α or ABCD3 abolished the interaction with p62, which failed to co-immunoprecipitate with the remaining partner (**Fig. 6A**). Given that ubiquitylation of ABCD3 acts as a signal for pexophagy^47,63,64^ and that ABCD3 was detected in lysosomal fractions in our lysosome immunoprecipitation mass spectrometry data, we next examined whether tether formation is associated with increased ubiquitylation of ABCD3. We enriched ubiquitylated proteins from cell lysates, in the presence or absence of Ceapin A7, and immunoblotted for ABCD3. We observed that Ceapin A7 treatment for 6 hours markedly increased ABCD3 ubiquitylation. We found that Ceapin A7-induced ubiquitylation was significantly reduced when ATF6α levels were reduced by CRISPRi, whereas basal levels of ABCD3 ubiquitylation in untreated cells were unaffected (**Fig. 6B**). These findings indicate that ATF6α is specifically required for Ceapin A7-induced ubiquitylation of ABCD3 and support a model in which ATF6α-dependent ABCD3 ubiquitylation leads to the initiation of pexophagy. Finally, to determine whether pexophagy induced by Ceapin A7 or THCA affects peroxisome abundance, we immunostained cells for the peroxisomal marker PEX14 (24 hours after treatment). Both Ceapin A7 and THCA caused a significant reduction in peroxisome number per cell. Notably, this decrease was prevented when either ATF6α or ABCD3 were depleted using sgRNAs (**Fig. 6C, Extended Data Fig. 8C)**.

**Figure 6:**
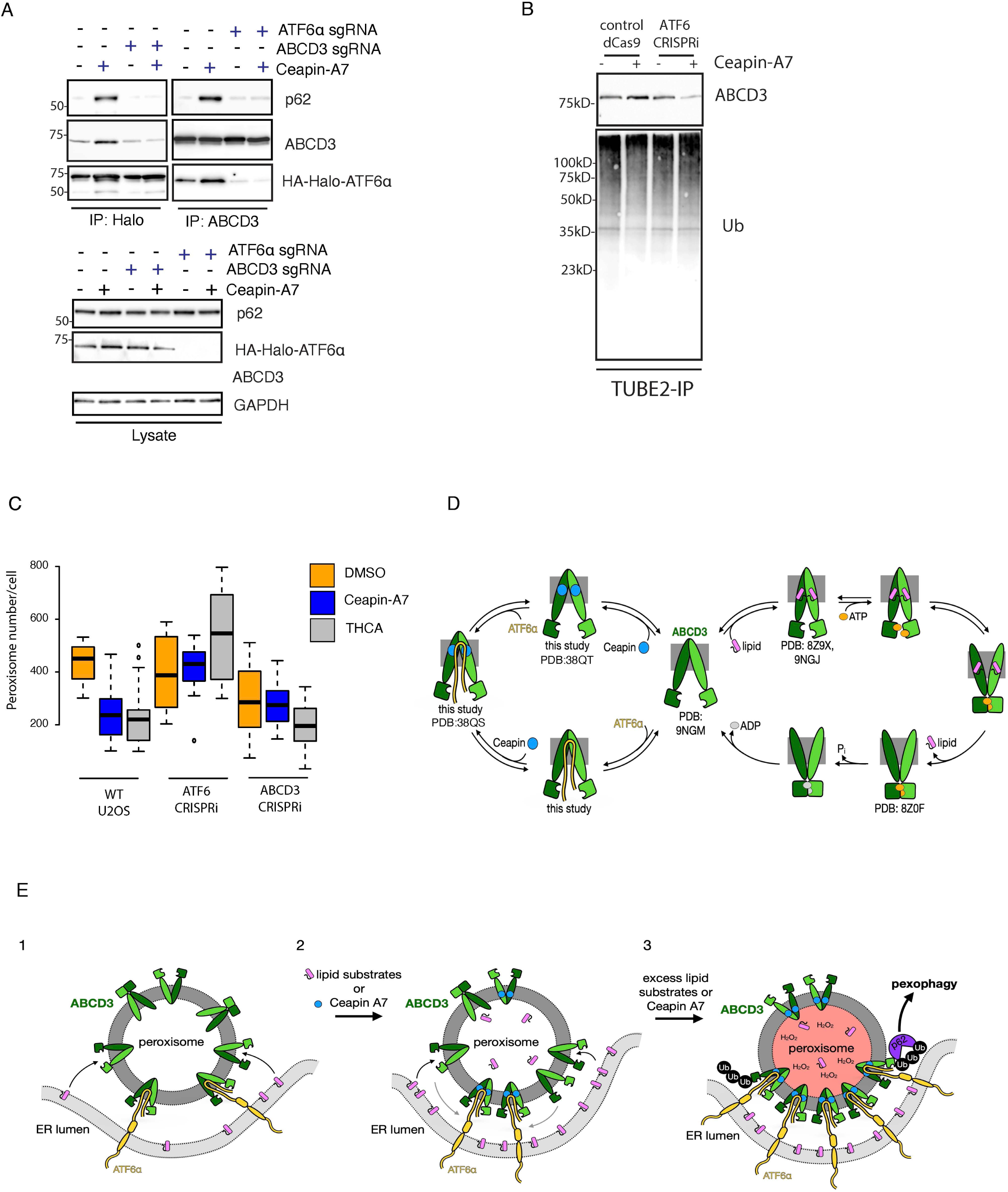
ATF6α and ABCD3 are required for p62 recruitment and pexophagy. **a**, Immunoprecipitation of 3×HA–Halo–ATF6α or ABCD3 from control and CRISPRi-depleted cells (targeting ATF6α or ABCD3), followed by immunoblotting for p62, showing that recruitment of p62 to the complex requires both proteins. Cells were treated with 10 μM Ceapin A7 for 6 h as indicated. **b**, Immunoblot analysis of ubiquitin-enriched fractions following TUBE2 pulldown, showing increased ABCD3 ubiquitylation upon 10 μM Ceapin A7 treatment (6 h), which is reduced upon ATF6α depletion. **c**, Quantification of peroxisome number per cell in wild-type and CRISPRi-depleted cells treated with DMSO, 10 μM Ceapin A7, or THCA for 16 h. Peroxisomes were stained with anti-PEX14. Center lines indicate medians; box limits represent the 25th and 75th percentiles; whiskers extend 1.5× the interquartile range; outliers are shown as individual points. n = 70 cells from 3 independent experiments. Statistical significance was determined using a two-tailed Student’s t-test. *P < 0.05, ***P < 0.001. **d**, Model illustrating how ATF6α and Ceapin A7 integrate within the ABCD3 ATPase cycle. ABCD3 (light and dark green), Ceapin A7 (blue), ATF6α (yellow), acyl-CoA lipid substrate (pink), ATP (orange), ADP (gray). PDBs referenced: 9NGM (ABCD3 apo), 8Z9X/9NGJ (ABCD3:phytanoyl-CoA complex), 8Z0F (ABCD3:ATP). **e**, Model illustrating the proposed mechanism. Panel 1: ATF6α transiently interacts with ABCD3 cycling through conformational states during lipid transport to form ER–peroxisome tethers. Panel 2: Addition of Ceapin A7 or ABCD3 lipid substrates promotes ABCD3:ATF6α complex formation and increased ER•peroxisome contact, nucleating ABCD3:ATF6α foci at the inter-organelle contact point. Panel 3: Treatment with excess Ceapin A7 or ABCD3 lipid substrates results in damaged peroxisomes with increased oxidation, p62 recruitment to the ABCD3:ATF6α complex, ABCD3 ubiquitylation, and pexophagy.

## Discussion

Cells maintain homeostasis not only through the function of individual organelles but through the coordinated direct physical interaction between them. At membrane contact sites, two organelles appose their membranes, exchanging lipids, calcium and other signals that allow one compartment to sense and respond to the state of its neighbor. This form of inter-organelle communication is emerging as a resilience mechanism in its own right: ER–mitochondria contacts relay calcium and lipid transfer that shapes mitochondrial fission and can be directly required for the autophagic clearance of damaged organelles, and mitochondria–lysosome contacts coordinate the timing of fission and mitophagy^65–69^. A loss of this coordination is increasingly linked to aging and disease^70–73^. Peroxisomes, despite their central role in lipid catabolism and redox homeostasis, have comparatively few known contact partners, and no established mechanism for signaling their own damage to the rest of the cell.

Here we identify such a mechanism. We discover a dynamic ER–peroxisome tether formed by the ER stress sensor ATF6α and the peroxisomal lipid transporter ABCD3. This tether forms when a cytosolic unstructured loop of ATF6α transiently inserts deep into the cytosol-exposed substrate-binding pocket of ABCD3, trapping ABCD3 in an off-pathway state (**Fig. 6D**). Importantly, we show that this interaction, which we initially believed to be a neomorphic consequence of treating cells with the ATF6α inhibitor Ceapin A7, occurs even in the absence of Ceapin A7, supporting its physiological relevance (**Fig. 6E**, left). We propose that the resulting organelle proximity serves to facilitate lipid transport from the ER into peroxisomes for catabolic processing, and that the same junction acts as a scaffold that can initiate peroxisome turnover by p62-triggered pexophagy once lipid processing generates excessive ROS (**Fig. 6E**, right). Thus, our work links the cell’s surveillance of lipid and protein homeostasis to ER and peroxisome health in an unexpected way.

### Discovery that an ABC transporter can form a membrane contact site

Our cryo-ET and single-particle cryo-EM data provide a detailed view of the ATF6α•ABCD3 tether at atomic resolution and explain how it is stabilized by Ceapin A7. Due to the dynamic nature of ABC transporters, the endogenous ATF6α•ABCD3 tether forms transiently in cells as ABCD3 undergoes its ATP-coupled conformational cycle and thereby dynamically exposes the cytosol-open conformation conducive to engage with ATF6 (**Fig. 6D**). ATF6α utilizes a unique interaction mode with ABCD3 that is specific and of high affinity: an unstructured region of ATF6α loops into the apex of the solvated transmembrane cavity of ABCD3, making extensive contact only possible in its apo, cytosol-open conformation. Although unprecedented for a membrane contact site, this interaction is reminiscent of ABC transporter structures in which unstructured regulatory domains or viral proteins adopt a helical-hairpin to plug the transmembrane cavity, stabilizing their respective apo, cytosol-open conformations^74–79^. While ATF6α does not adopt any observable secondary structure in the complex, the apparent effect on the ABC transporter is similar: ABCD3 is stabilized in an off-pathway conformation from which the regulator —ATF6α— must be released prior to substrate binding. In this case, the off-pathway state allows ABCD3 to adopt a new function as an ER-peroxisome tether that is responsive to lipid import.

### Ceapin A7’s mechanism of action

We also demonstrate in mechanistic detail how the small molecule Ceapin A7 acts as a molecular glue to stabilize the ATF6α•ABCD3 interaction. Ceapin A7 engages ABCD3 in a unique manner, in which two molecules of Ceapin A7 bind symmetrically to the ABCD3 homodimer, but cement two distinct interfaces formed by each ABCD3 protomer and ATF6α. The bulk of contact is at the Ceapin A7•ABCD3 interface where the Ceapin A7 molecules occupy a solvated pocket within the ABCD3 transmembrane cavity normally used for substrate binding. This binding mode allows Ceapin A7 to stabilize the ATF6α•ABCD3 complex in three ways: 1) lock the cytosol-open conformation of ABCD3, which ATF6α binds specifically, 2) act as a molecular glue to stabilize the ATF6α•ABCD3 interface, and 3) directly inhibit binding of ABCD3 substrates, which would compete with ATF6α binding. To our knowledge, Ceapin A7 is the first substrate-competitive inhibitor of ABCD3 described. As a consequence of this interaction, ATF6α becomes trapped in the ER at the junction sites, which inhibits its activation– the phenotype that was read out in the screen that led to Ceapin’s discovery^6^. Our structure now provides a framework to improve Ceapin’s potency and rationally manipulate its diverse functions, as we demonstrate.

### A new role for ATF6α in regulating lipid metabolism

Our study also establishes a previously unknown “moonlighting” function of ATF6α in regulating lipid metabolism and pexophagy, in which ATF6α forms a reversible tether to peroxisomes via ABCD3 in activity-dependent manner (**Fig. 6E**, left). Given the apparently low cellular stoichiometry of ATF6α *versus* ABCD3 (∼1:18, https://opencell.sf.czbiohub.org/), only a subset of ABCD3 molecules can engage in the ER-peroxisome tether, trapping all ATF6α molecules available but leaving excess ABCD3 available to transport substrates at the ER-peroxisome contact site. We pose that tethering by enhancing the proximity between the respective organelles serves to facilitate transfer of ER-derived lipid substrates—including bile acid intermediates, oxidized lipids, long-chain fatty acids, and dicarboxylic acids—into peroxisomes for degradation and metabolism^56,80^. Cryo-ET reveals the ATF6α•ABCD3 tether mediates an extensive ER-peroxisome contact site in which peroxisomes are partially enveloped by the ER. Although no membrane fusion is observed, the close proximity (∼23 nm) is reminiscent of described membrane contact sites and could allow for non-vesicular transport of these substrates^69,81,82^. Notably, the ATF6α•ABCD3 contact site is distinct from the previously described VAPA/B-mediated ER-peroxisome tether^7^, which also has been suggested to facilitate lipid exchange between the two organelles^21^. These disparate molecular contact sites may form in distinct cellular contexts or to facilitate transfer of distinct lipid substrates.

### How Ceapin A7 and lipid substrates affect the ATF6α•ABCD3 tether

Treatment with Ceapin A7 or addition of ABCD3 lipid transport substrates stabilizes the ATF6α•ABCD3 inter-organelle tether (**Fig. 6E**, middle). Trapped ATF6α•ABCD3 complexes accumulate as the ER-peroxisome junction expands, nucleating additional tethers and producing the observed ATF6α•ABCD3 puncta in cells. Notably, while ABCD3 substrate binding reduced ATF6α•ABCD3 binding *in vitro*, substrate treatment *in cellula* increased puncta formation (**Fig. 3E** and **Fig. 4F**, respectively). In the *in cellula* split-Venus assay, fluorescent complementation effectively trapped ATF6α•ABCD3 interactions (due to the extremely slow dissociation rate of the reconstituting Venus parts), whereas inhibition of lysosomal degradation (e.g., with chloroquine) resulted in accumulation of tethers that are otherwise transient, targeting them for autophagic turnover. Indeed, we showed that these stabilized ER-peroxisome tethers are associated with peroxisomal swelling and increased oxidative stress, hallmarks of organelle dysfunction, underscoring the notion that peroxisomes have become over-loaded under these conditions and require autophagic clearance^29,50,51^.

### A new role of the ATF6α•ABCD3 tether in pexophagy

Mechanistically, we showed that the autophagy adaptor p62 is selectively recruited to the ATF6α•ABCD3 tether, where it promotes pexophagy (**Fig. 6E**, right). Consistent with this observation, Ceapin-induced tether formation enhanced ABCD3 ubiquitylation in an ATF6α-dependent manner. Given that ubiquitylation of ABCD3 has been implicated in p62-mediated peroxisomal degradation^47,63,64^, these findings suggest that ATF6α-dependent tether formation couples lipid import to selective peroxisome turnover. While the ubiquitin machinery responsible for this modification remains undefined, multiple E3 ligases and deubiquitinases have been shown to regulate peroxisome turnover^63,83,84^. These or analogous pathways may act on ABCD3 under lipid stress conditions. ATF6α turnover has been primarily attributed to proteasomal degradation following ER stress^85,86^. However, we detect a small amount of ATF6α in lysosomal fractions, and inhibition of lysosomal degradation increases ATF6α levels. Notably, ER resident proteins were not enriched in lysosomal fractions under these conditions, arguing against bulk ER turnover. Instead, these observations suggest that only a subset of ATF6α associated with the ATF6α•ABCD3 tether is delivered to lysosomes, perhaps in the context of peroxisome-directed autophagy, but the mechanism remains to be determined. We propose that the ATF6α•ABCD3 tether functions as a metabolically responsive quality-control checkpoint that marks dysfunctional peroxisomes for autophagic clearance.

### Context-dependent functions of ATF6α

Beyond the canonical role of ATF6α to remediate protein folding stress in the ER by upregulating components of the ER-lumenal protein folding machinery, ATF6α has been shown to induce antioxidant genes, including catalase and peroxiredoxins, linking ER stress to oxidative stress responses^35^. Although this latter context is distinct from the lipid-driven stress conditions examined here, these findings suggest that ATF6α may regulate redox homeostasis through both transcriptional and organelle-level mechanisms. ATF6α has also been shown to be activated by specific lipid perturbations, including ceramide accumulation in the ER membrane^34^. However, induction of ATF6α under such conditions does not alter ER–peroxisome contacts, indicating that ER stress driven ATF6α activation does not drive tether formation. Instead, our findings suggest that formation of the ATF6α•ABCD3 tether is specifically coupled to the activity state of ABCD3 and peroxisomal lipid import, rather than general ER stress or lipid perturbation. This distinction highlights a context-dependent, non-transcriptional function of ATF6α that is selectively engaged under lipid flux conditions linked to peroxisomal metabolism.

### How Ceapins could be used for therapeutic purposes

Due to its lack of structured domains or identifiable binding sites, ATF6α was regarded as ‘undruggable’. We show that a molecular glue can be used to stabilize protein complexes containing intrinsically disordered proteins, demonstrating a potentially effective strategy to drug such obstinate targets. Elucidation of Ceapin A7’s mechanism of action has led to the discovery of new biology–the ATF6α•ABCD3 inter-organelle tether–and a basis to improve Ceapins for therapeutic use. We find Ceapin A7 has a multitude of diverse functions within the cell that may prove therapeutically beneficial: stabilizing ER-peroxisome contacts, inhibiting ATF6α activation and ABCD3 enzymatic activity, promoting lysosomal degradation of the ATF6α•ABCD3 complex, and inducing pexophagy. Thus, Ceapin A7 has the potential to enter the growing therapeutic space exploiting induced proximity^87^. In line with this idea, ATF6α activation has been shown to drive colorectal cancer (CRC) and hepatocellular carcinoma, as well as contribute to chemoresistance^15–18^. Indeed, Ceapin A7 has been shown to inhibit growth of cultured CRC cell lines and organoids^16^. Furthermore, ABCD3 activity has recently been shown to promote chemoresistance in CRC^88^. Ceapin A7 and its pharmacologically improved analogs ([U.S. Patent No. 12,325,702](https://patents.google.com/patent/US12325702B2/en?oq=US12325702B2)) thus may provide a promising therapeutic strategy against ATF6α/ABCD3-dependent cancers or, more broadly, be employed to improve cell health by promoting turnover of damaged peroxisomes.

## Data Availability

ATF6α•ABCD3•Ceapin A7 and ABCD3•Ceapin A7 cryo-EM density maps and associated atomic coordinates are publicly available through the PDB (38QS, 38QT, respectively) and the EMDB (EMD-79063, EMD-79064, respectively).

## Acknowledgements

We thank Emma McCullagh for helpful discussions and design of fluorescent Ceapin molecules. We thank Melissa Mendez for the design of tiled ATF6α fluorescent peptides. We thank Nico Stuurman for help with light microscopy data collection. We thank the Costa-Mattioli lab and the Walter lab for helpful discussions.

## Author Contributions

C.M.N. designed and performed biochemical experiments, cryo-EM sample preparation, data processing, model building, and data analysis. R.M. designed and performed cell biology and microscopy experiments, data processing and analysis, and mass spectrometry sample preparation and analysis. C.P. designed, performed, and analyzed nanoBRET and HDX-MS experiments under the supervision of J.R. and A.A. T.G.L. collected cryo-EM data and processed cryo-EM and cryo-ET data. J.W. collected cryo-ET data and performed data processing. B.T.W. performed HDX-MS experiments and data processing and analysis. T.L. designed and performed cryo-ET sample preparation. T.C. performed cryo-EM model building. D.E. designed and synthesized CoA hybrid compounds. S.H. generated the endogenous Halo-ATF6α knock-in cell line. A.X. performed mass spectrometry experiments. K.K. analyzed mass spectrometry data. M.G. provided purified ABCD3 for pilot studies and guidance on cryo-EM sample preparation. J.J.C. supervised CoA hybrid compound design and synthesis. C.P.A. collected and processed cryo-EM data. D.I. supervised mass spectrometry experiments. M.C.M., J.R., and A.A. provided guidance on the project and manuscript preparation. C.M.N., R.M., and P.W. conceived the project, interpreted the results, and co-wrote the manuscript. All authors reviewed and edited the manuscript.

## References

1. Yoshida, H., Haze, K., Yanagi, H., Yura, T. & Mori, K. Identification of the cis-acting endoplasmic reticulum stress response element responsible for transcriptional induction of mammalian glucose-regulated proteins. Involvement of basic leucine zipper transcription factors: Involvement of basic leucine zipper transcription factors. J. Biol. Chem. 273, 33741–33749 (1998).

2. Haze, K., Yoshida, H., Yanagi, H., Yura, T. & Mori, K. Mammalian transcription factor ATF6 is synthesized as a transmembrane protein and activated by proteolysis in response to endoplasmic reticulum stress. Mol. Biol. Cell 10, 3787–3799 (1999).

3. Shen, J., Chen, X., Hendershot, L. & Prywes, R. ER stress regulation of ATF6 localization by dissociation of BiP/GRP78 binding and unmasking of Golgi localization signals. Dev. Cell 3, 99– 111 (2002).

4. Ye, J. et al. ER stress induces cleavage of membrane-bound ATF6 by the same proteases that process SREBPs. Mol. Cell 6, 1355–1364 (2000).

5. Gallagher, C. M. & Walter, P. Ceapins inhibit ATF6α signaling by selectively preventing transport of ATF6α to the Golgi apparatus during ER stress. Elife 5, e11880 (2016).

6. Gallagher, C. M. et al. Ceapins are a new class of unfolded protein response inhibitors, selectively targeting the ATF6α branch. Elife 5, e11878 (2016).

7. Torres, S. E. et al. Ceapins block the unfolded protein response sensor ATF6α by inducing a neomorphic inter-organelle tether. Elife 8, e46595 (2019).

8. Walter, P. & Ron, D. The unfolded protein response: from stress pathway to homeostatic regulation. Science 334, 1081–1086 (2011).

9. Hetz, C., Zhang, K. & Kaufman, R. J. Mechanisms, regulation and functions of the unfolded protein response. Nat. Rev. Mol. Cell Biol. 21, 421–438 (2020).

10. Acosta-Alvear, D., Harnoss, J. M., Walter, P. & Ashkenazi, A. Homeostasis control in health and disease by the unfolded protein response. Nat. Rev. Mol. Cell Biol. 26, 193–212 (2025).

11. Chen, X., Shen, J. & Prywes, R. The luminal domain of ATF6 senses endoplasmic reticulum (ER) stress and causes translocation of ATF6 from the ER to the Golgi. J. Biol. Chem. 277, 13045–13052 (2002).

12. Yamamoto, K. et al. Transcriptional induction of mammalian ER quality control proteins is mediated by single or combined action of ATF6alpha and XBP1. Dev. Cell 13, 365–376 (2007).

13. Adachi, Y. et al. ATF6 is a transcription factor specializing in the regulation of quality control proteins in the endoplasmic reticulum. Cell Struct. Funct. 33, 75–89 (2008).

14. Wu, J. et al. ATF6alpha optimizes long-term endoplasmic reticulum function to protect cells from chronic stress. Dev. Cell 13, 351–364 (2007).

15. Schewe, D. M. & Aguirre-Ghiso, J. A. ATF6alpha-Rheb-mTOR signaling promotes survival of dormant tumor cells in vivo. Proc. Natl. Acad. Sci. U. S. A. 105, 10519–10524 (2008).

16. Rodvold, J. J. et al. ATF6 promotes colorectal cancer growth and stemness by regulating the Wnt pathway. Cancer Res. Commun. 4, 2734–2755 (2024).

17. Su, S. et al. ATF6 activation promotes tumorigenesis and drug resistance in diffuse large B-cell lymphoma (DLBCL) by regulating the mTOR/S6K signaling pathway. *Discov*. Oncol. 16, 499 (2025).

18. Li, X. et al. Activated ATF6α is a hepatic tumour driver restricting immunosurveillance. Nature 651, 796–807 (2026).

19. Costello, J. L., Castro, I. G., Schrader, T. A., Islinger, M. & Schrader, M. Peroxisomal ACBD4 interacts with VAPB and promotes ER-peroxisome associations. Cell Cycle 16, 1039–1045 (2017).

20. Costello, J. L. et al. ACBD5 and VAPB mediate membrane associations between peroxisomes and the ER. J. Cell Biol. 216, 331–342 (2017).

21. Hua, R. et al. VAPs and ACBD5 tether peroxisomes to the ER for peroxisome maintenance and lipid homeostasis. J. Cell Biol. 216, 367–377 (2017).

22. Wanders, R. J. A. & Waterham, H. R. Biochemistry of mammalian peroxisomes revisited. Annu. Rev. Biochem. 75, 295–332 (2006).

23. Schrader, M., Costello, J., Godinho, L. F. & Islinger, M. Peroxisome-mitochondria interplay and disease. J. Inherit. Metab. Dis. 38, 681–702 (2015).

24. Morita, M. & Imanaka, T. Peroxisomal ABC transporters: structure, function and role in disease. Biochim. Biophys. Acta 1822, 1387–1396 (2012).

25. van Roermund, C. W. T. et al. The human peroxisomal ABC half transporter ALDP functions as a homodimer and accepts acyl-CoA esters. FASEB J. 22, 4201–4208 (2008).

26. Berger, J. et al. The four murine peroxisomal ABC-transporter genes differ in constitutive, inducible and developmental expression. Eur. J. Biochem. 265, 719–727 (1999).

27. Imanaka, T. et al. Characterization of the 70-kDa peroxisomal membrane protein, an ATP binding cassette transporter. J. Biol. Chem. 274, 11968–11976 (1999).

28. van Roermund, C. W. T., Ijlst, L., Wagemans, T., Wanders, R. J. A. & Waterham, H. R. A role for the human peroxisomal half-transporter ABCD3 in the oxidation of dicarboxylic acids. Biochim. Biophys. Acta 1841, 563–568 (2014).

29. Ferdinandusse, S. et al. A novel bile acid biosynthesis defect due to a deficiency of peroxisomal ABCD3. Hum. Mol. Genet. 24, 361–370 (2015).

30. Violante, S. et al. Peroxisomes can oxidize medium- and long-chain fatty acids through a pathway involving ABCD3 and HSD17B4. FASEB J. 33, 4355–4364 (2019).

31. Ranea-Robles, P. et al. The peroxisomal transporter ABCD3 plays a major role in hepatic dicarboxylic fatty acid metabolism and lipid homeostasis. J. Inherit. Metab. Dis. 44, 1419–1433 (2021).

32. Li, Y. et al. Structural insights into human ABCD3-mediated peroxisomal acyl-CoA translocation. Cell Discov. 10, 92 (2024).

33. Gupta, M. et al. Molecular mechanism of substrate transport by human peroxisomal ABCD3. Proc. Natl. Acad. Sci. U. S. A. 122, e2513928122 (2025).

34. Tam, A. B. et al. The UPR activator ATF6 responds to proteotoxic and lipotoxic stress by distinct mechanisms. Dev. Cell 46, 327–343.e7 (2018).

35. Jin, J.-K. et al. ATF6 decreases myocardial ischemia/reperfusion damage and links ER stress and oxidative stress signaling pathways in the heart. Circ. Res. 120, 862–875 (2017).

36. Blackwood, E. A. et al. ATF6 regulates cardiac hypertrophy by transcriptional induction of the mTORC1 activator, rheb. Circ. Res. 124, 79–93 (2019).

37. Glembotski, C. C., Arrieta, A., Blackwood, E. A. & Stauffer, W. T. ATF6 as a nodal regulator of proteostasis in the heart. Front. Physiol. 11, 267 (2020).

38. Marsh, K. G. et al. The peroxisomal enzyme, FAR1, is induced during ER stress in an ATF6-dependent manner in cardiac myocytes. Am. J. Physiol. Heart Circ. Physiol. 320, H1813–H1821 (2021).

39. Yamamoto, K. et al. Induction of liver steatosis and lipid droplet formation in ATF6alpha-knockout mice burdened with pharmacological endoplasmic reticulum stress. Mol. Biol. Cell 21, 2975–2986 (2010).

40. Chen, X. et al. Hepatic ATF6 increases fatty acid oxidation to attenuate hepatic steatosis in mice through peroxisome proliferator-activated receptor α. Diabetes 65, 1904–1915 (2016).

41. Zhang, J. et al. ATM functions at the peroxisome to induce pexophagy in response to ROS. Nat. Cell Biol. 17, 1259–1269 (2015).

42. Lee, J. N. et al. Catalase inhibition induces pexophagy through ROS accumulation. Biochem. Biophys. Res. Commun. 501, 696–702 (2018).

43. Farré, J.-C., Mahalingam, S. S., Proietto, M. & Subramani, S. Peroxisome biogenesis, membrane contact sites, and quality control. EMBO Rep. 20, e46864 (2019).

44. Bajdzienko, J. & Bremm, A. Mammalian pexophagy at a glance. J. Cell Sci. 137, jcs259775 (2024).

45. Kim, P. K., Hailey, D. W., Mullen, R. T. & Lippincott-Schwartz, J. Ubiquitin signals autophagic degradation of cytosolic proteins and peroxisomes. Proc. Natl. Acad. Sci. U. S. A. 105, 20567– 20574 (2008).

46. Deosaran, E. et al. NBR1 acts as an autophagy receptor for peroxisomes. J. Cell Sci. 126, 939–952 (2013).

47. Jo, D. S. et al. PINK1 and STUB1 pathway orchestrates peroxisomal selective autophagy by PEX13 depletion. Cell Death Differ. 1–13 (2026).

48. Grumati, P., Dikic, I. & Stolz, A. ER-phagy at a glance. J. Cell Sci. 131, jcs217364 (2018).

49. Wilkinson, S. ER-phagy: shaping up and destressing the endoplasmic reticulum. FEBS J. 286, 2645–2663 (2019).

50. Poll-The, B. T. et al. A new peroxisomal disorder with enlarged peroxisomes and a specific deficiency of acyl-CoA oxidase (pseudo-neonatal adrenoleukodystrophy). Am. J. Hum. Genet. 42, 422–434 (1988).

51. Funato, M. et al. Aberrant peroxisome morphology in peroxisomal beta-oxidation enzyme deficiencies. Brain Dev. 28, 287–292 (2006).

52. Thomas, C. & Tampé, R. Structural and mechanistic principles of ABC transporters. Annu. Rev. Biochem. 89, 605–636 (2020).

53. Croucher, D. R. et al. Bimolecular complementation affinity purification (BiCAP) reveals dimer-specific protein interactions for ERBB2 dimers. Sci. Signal. 9, ra69 (2016).

54. Fujimoto, Y. et al. Involvement of ACSL in local synthesis of neutral lipids in cytoplasmic lipid droplets in human hepatocyte HuH7. J. Lipid Res. 48, 1280–1292 (2007).

55. Wei, Y., Wang, D., Topczewski, F. & Pagliassotti, M. J. Saturated fatty acids induce endoplasmic reticulum stress and apoptosis independently of ceramide in liver cells. Am. J. Physiol. Endocrinol. Metab. 291, E275–81 (2006).

56. Ferdinandusse, S., Denis, S., Faust, P. L. & Wanders, R. J. A. Bile acids: the role of peroxisomes. J. Lipid Res. 50, 2139–2147 (2009).

57. Bilan, D. S. et al. HyPer-3: a genetically encoded H(2)O(2) probe with improved performance for ratiometric and fluorescence lifetime imaging. ACS Chem. Biol. 8, 535–542 (2013).

58. Van den Branden, C., Kerckaert, I. & Roels, F. Peroxisomal beta-oxidation from endogenous substrates. Demonstration through H2O2 production in the unanaesthetized mouse. Biochem. J. 218, 697–702 (1984).

59. Kritsiligkou, P. et al. Proteome-wide tagging with an H2O2 biosensor reveals highly localized and dynamic redox microenvironments. Proc. Natl. Acad. Sci. U. S. A. 120, e2314043120 (2023).

60. Korotkova, D. et al. Fluorescent fatty acid conjugates for live cell imaging of peroxisomes. Nat. Commun. 15, 4314 (2024).

61. Zheng, J., Chen, X., Liu, Q., Zhong, G. & Zhuang, M. Ubiquitin ligase MARCH5 localizes to peroxisomes to regulate pexophagy. J. Cell Biol. 221, e202103156 (2022).

62. Abu-Remaileh, M. et al. Lysosomal metabolomics reveals V-ATPase- and mTOR-dependent regulation of amino acid efflux from lysosomes. Science 358, 807–813 (2017).

63. Sargent, G. et al. PEX2 is the E3 ubiquitin ligase required for pexophagy during starvation. J. Cell Biol. 214, 677–690 (2016).

64. Koyano, F. et al. AAA+ ATPase chaperone p97/VCPFAF2 governs basal pexophagy. Nat. Commun. 15, 9347 (2024).

65. Ma, X. et al. Mitochondria-lysosome-related organelles mediate mitochondrial clearance during cellular dedifferentiation. Cell Rep. 42, 113291 (2023).

66. Nakamura, K. et al. Mitochondrial complexity is regulated at ER-mitochondria contact sites via PDZD8-FKBP8 tethering. Nat. Commun. 16, 3401 (2025).

67. Hirabayashi, Y. et al. ER-mitochondria tethering by PDZD8 regulates Ca2+ dynamics in mammalian neurons. Science 358, 623–630 (2017).

68. John Peter, A. T., Petrungaro, C., Peter, M. & Kornmann, B. METALIC reveals interorganelle lipid flux in live cells by enzymatic mass tagging. Nat. Cell Biol. 24, 996–1004 (2022).

69. Kornmann, B. et al. An ER-mitochondria tethering complex revealed by a synthetic biology screen. Science 325, 477–481 (2009).

70. Obara, C. J. et al. Motion of VAPB molecules reveals ER-mitochondria contact site subdomains. Nature 626, 169–176 (2024).

71. Sun, S. et al. TBC1D15-Drp1 interaction-mediated mitochondrial homeostasis confers cardioprotection against myocardial ischemia/reperfusion injury. Metabolism 134, 155239 (2022).

72. Prinz, W. A., Toulmay, A. & Balla, T. The functional universe of membrane contact sites. Nat. Rev. Mol. Cell Biol. 21, 7–24 (2020).

73. Wang, X. et al. The bridge-like lipid transport protein VPS13C/PARK23 mediates ER-lysosome contacts following lysosome damage. Nat. Cell Biol. 27, 776–789 (2025).

74. Oldham, M. L., Grigorieff, N. & Chen, J. Structure of the transporter associated with antigen processing trapped by herpes simplex virus. Elife 5, e21829 (2016).

75. Oldham, M. L. et al. A mechanism of viral immune evasion revealed by cryo-EM analysis of the TAP transporter. Nature 529, 537–540 (2016).

76. Khandelwal, N. K. & Tomasiak, T. M. Structural basis for autoinhibition by the dephosphorylated regulatory domain of Ycf1. Nat. Commun. 15, 2389 (2024).

77. Mazza, T. et al. Structural basis for the modulation of MRP2 activity by phosphorylation and drugs. Nat. Commun. 15, 1983 (2024).

78. Koide, E., Pietz, H. L., Beltran, J. & Chen, J. Structural basis for the transport and regulation mechanism of the multidrug resistance-associated protein 2. Nat. Commun. 16, 484 (2025).

79. Huang, Y. et al. Inhibition and transport mechanisms of the ABC transporter hMRP5. Nat. Commun. 15, 4811 (2024).

80. Kawaguchi, K. & Morita, M. ABC transporter subfamily D: Distinct differences in behavior between ABCD1-3 and ABCD4 in subcellular localization, function, and human disease. Biomed Res. Int. 2016, 6786245 (2016).

81. Scorrano, L. et al. Coming together to define membrane contact sites. Nat. Commun. 10, 1287 (2019).

82. Phillips, M. J. & Voeltz, G. K. Structure and function of ER membrane contact sites with other organelles. Nat. Rev. Mol. Cell Biol. 17, 69–82 (2016).

83. Riccio, V. et al. Deubiquitinating enzyme USP30 maintains basal peroxisome abundance by regulating pexophagy. J. Cell Biol. 218, 798–807 (2019).

84. Marcassa, E. et al. New aspects of USP30 biology in the regulation of pexophagy. Autophagy 15, 1634–1637 (2019).

85. Thuerauf, D. J., Morrison, L. E., Hoover, H. & Glembotski, C. C. Coordination of ATF6-mediated transcription and ATF6 degradation by a domain that is shared with the viral transcription factor, VP16. J. Biol. Chem. 277, 20734–20739 (2002).

86. Hong, M., Li, M., Mao, C. & Lee, A. S. Endoplasmic reticulum stress triggers an acute proteasome-dependent degradation of ATF6. J. Cell. Biochem. 92, 723–732 (2004).

87. King, E. A., Meyers, M. & Nomura, D. K. Induced proximity-based therapeutic modalities. Nat. Rev. Drug Discov. 25, 175–203 (2026).

88. Yin, J. et al. Peroxisomal membrane protein PMP70 confers drug resistance in colorectal cancer. Cell Death Dis. 16, 293 (2025).

