## Supplementary Data for "Peroxisome quality control is gated by a dynamic ATF6α–ABCD3 tether"

###### **This PDF file includes:**

Extended Data Figures 1-8

Extended Data Figure Legends

Table 1

Extended Data Figure 1: Tomography of ATF6α-GFP puncta

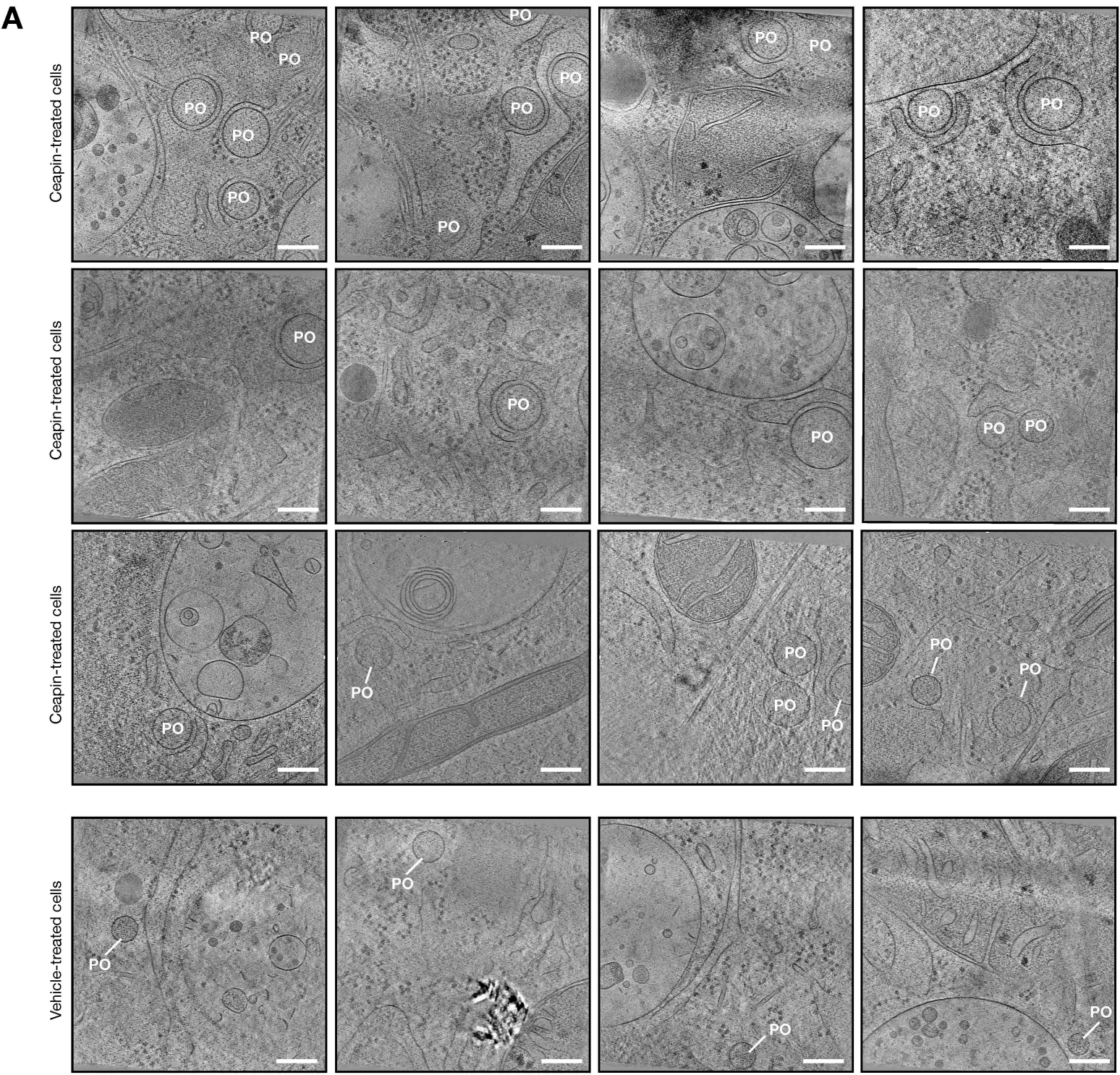

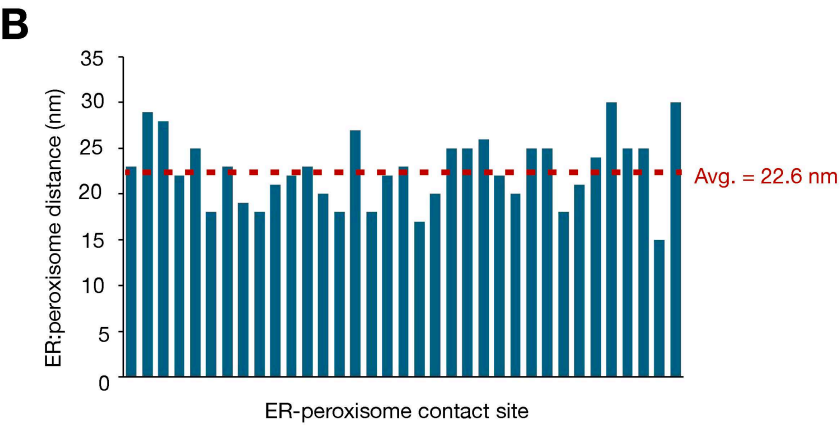

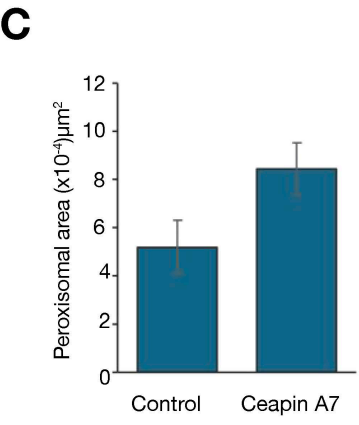

### Extended Data Figure 2: Cryo-EM of ATF6α•ABCD3•Ceapin A7 complex

**A**

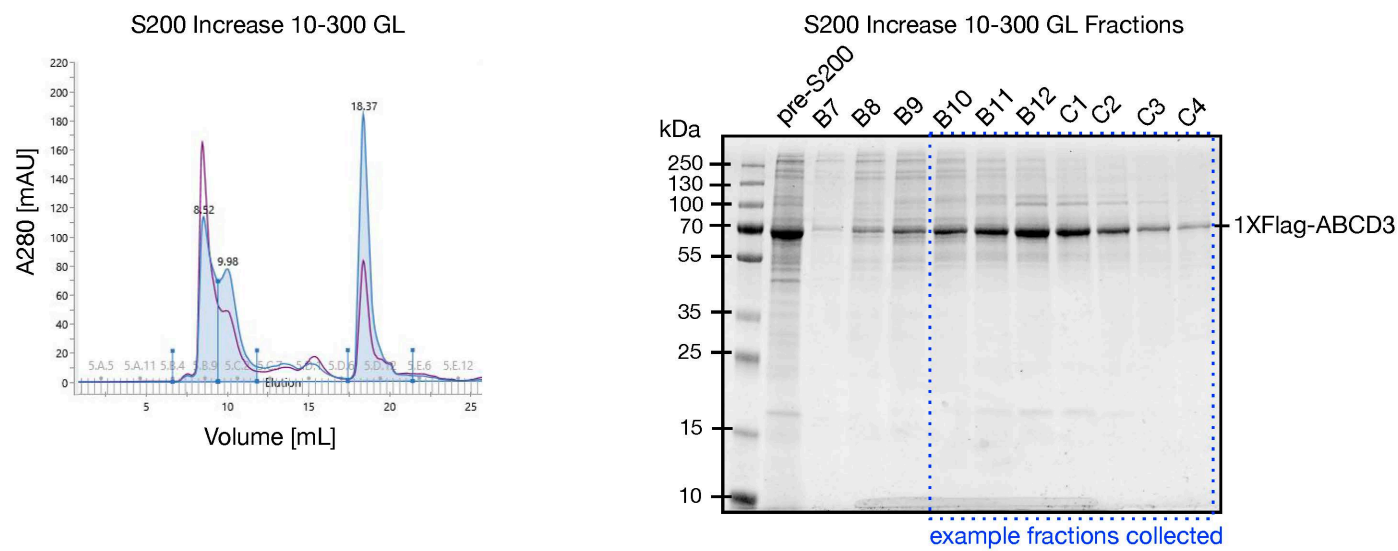

**B**

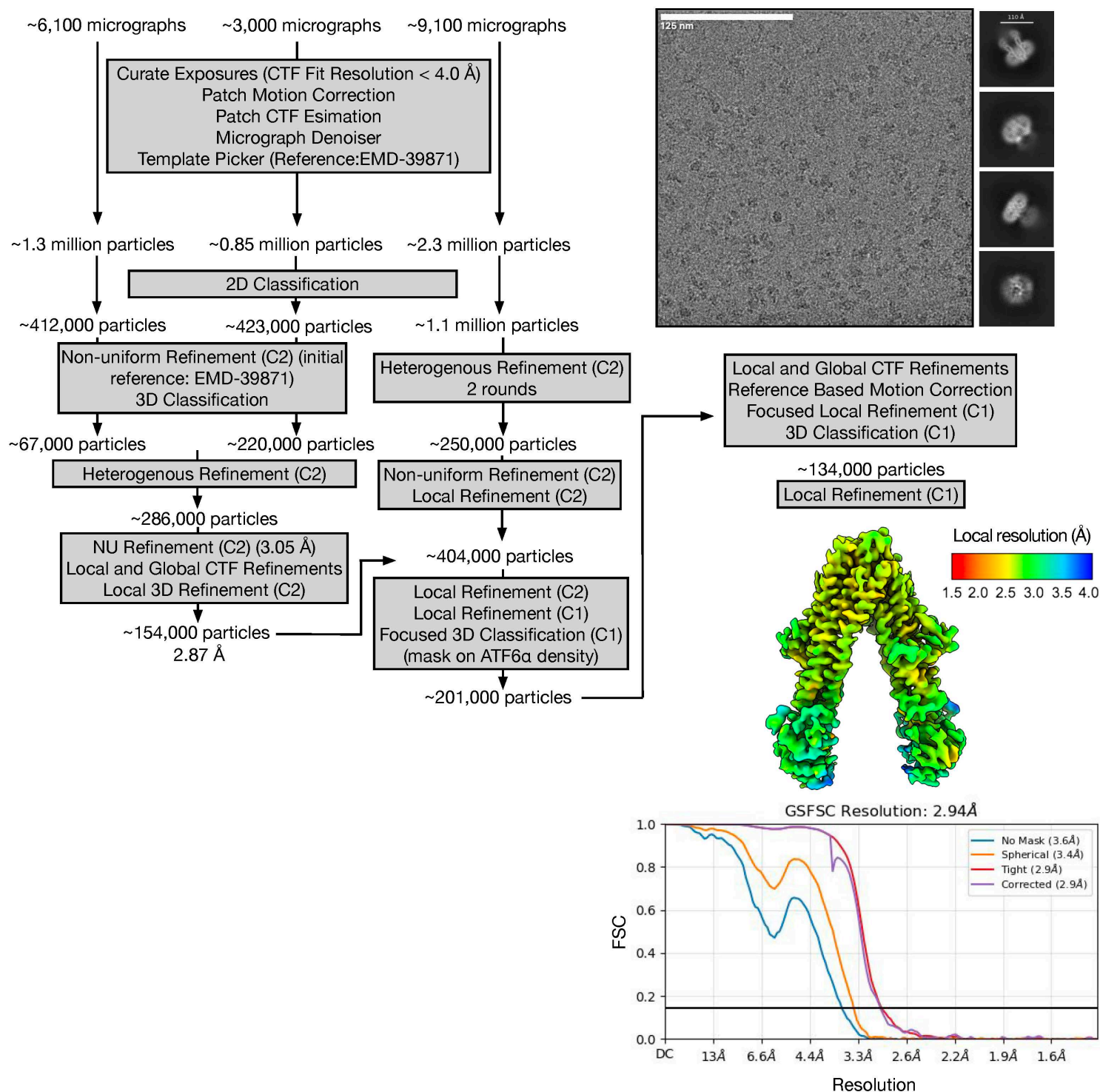

Extended Data Figure 3: Cryo-EM of ABCD3•Ceapin A7 complex

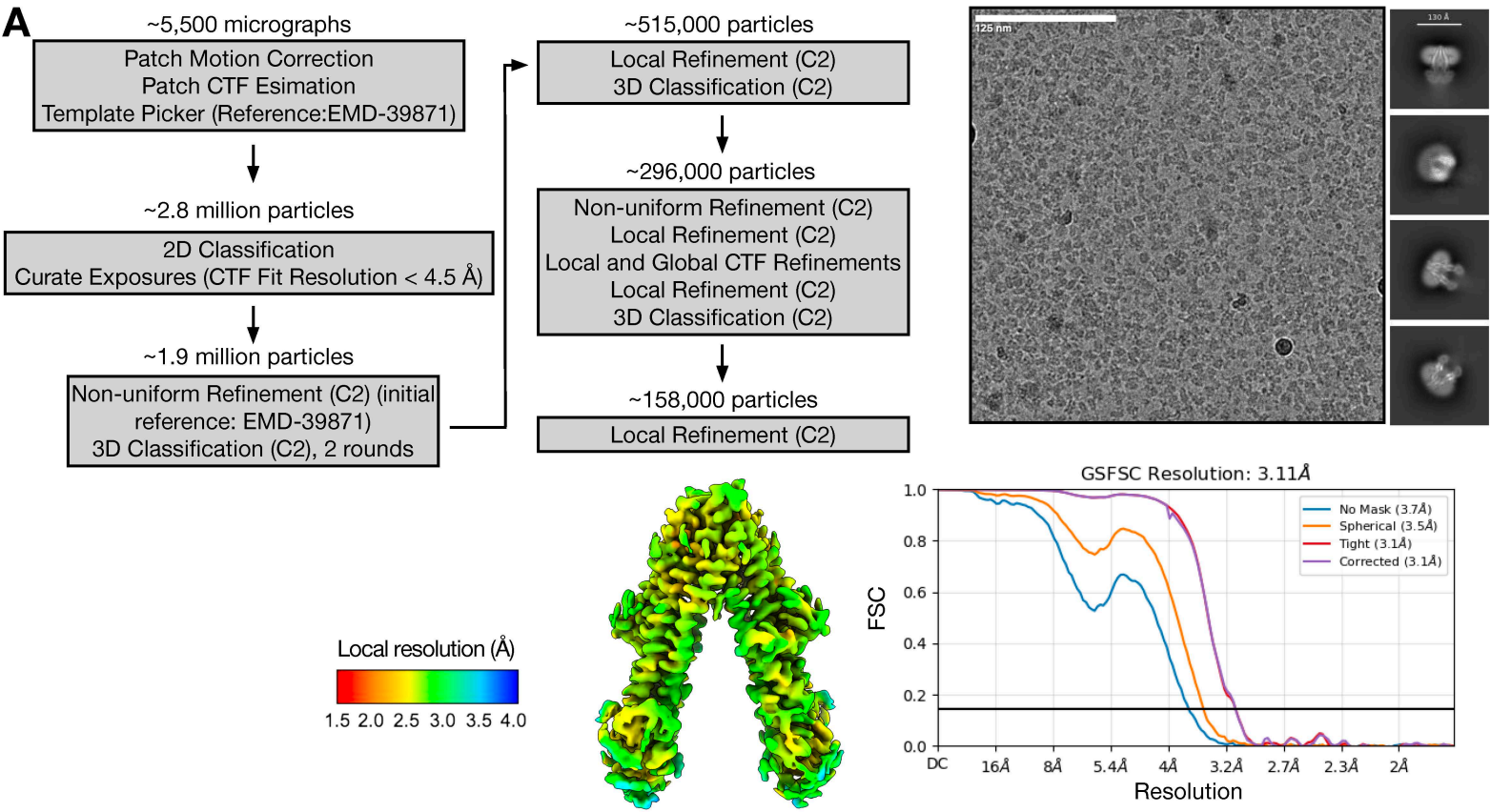

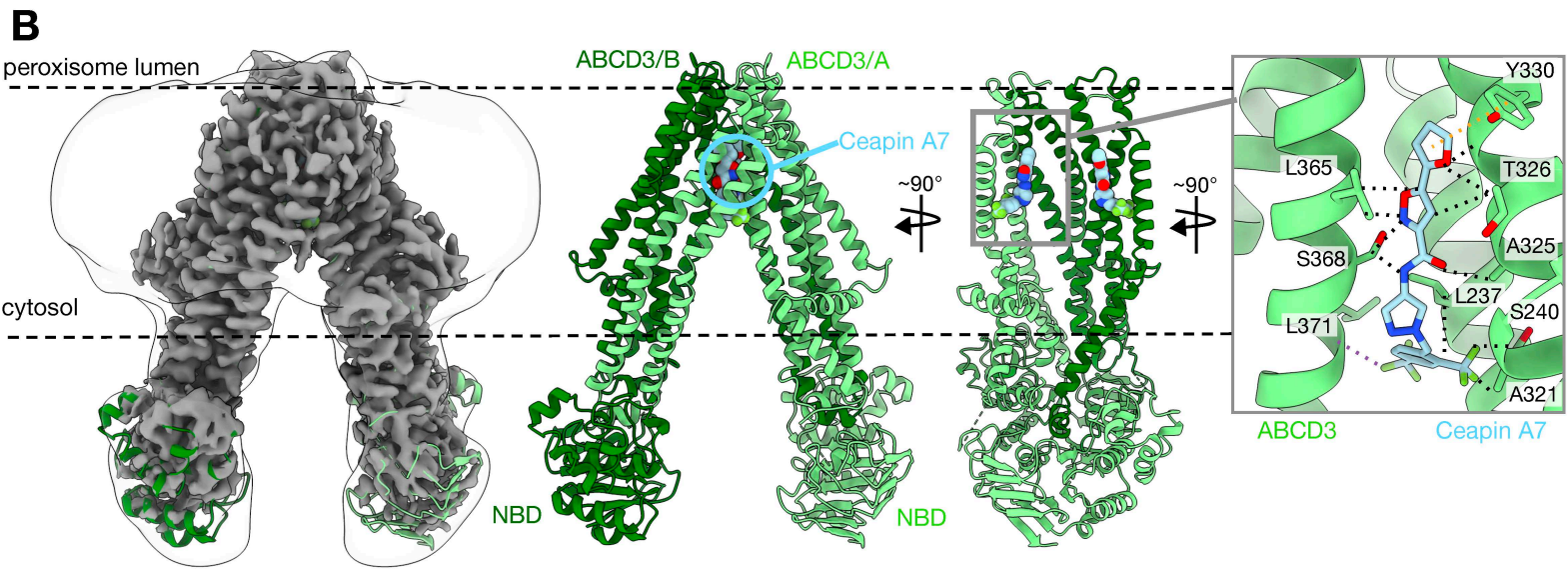

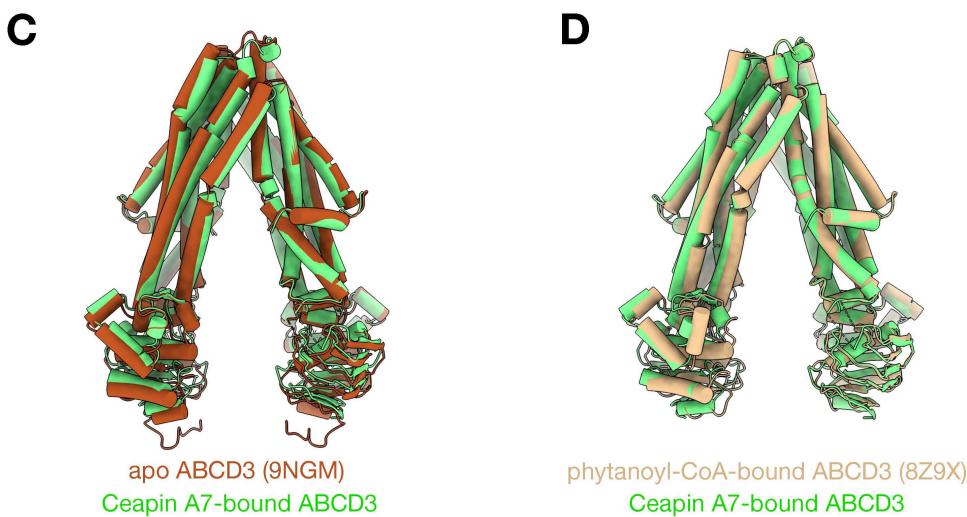

Extended Data Figure 4: Assigning the ATF6α register

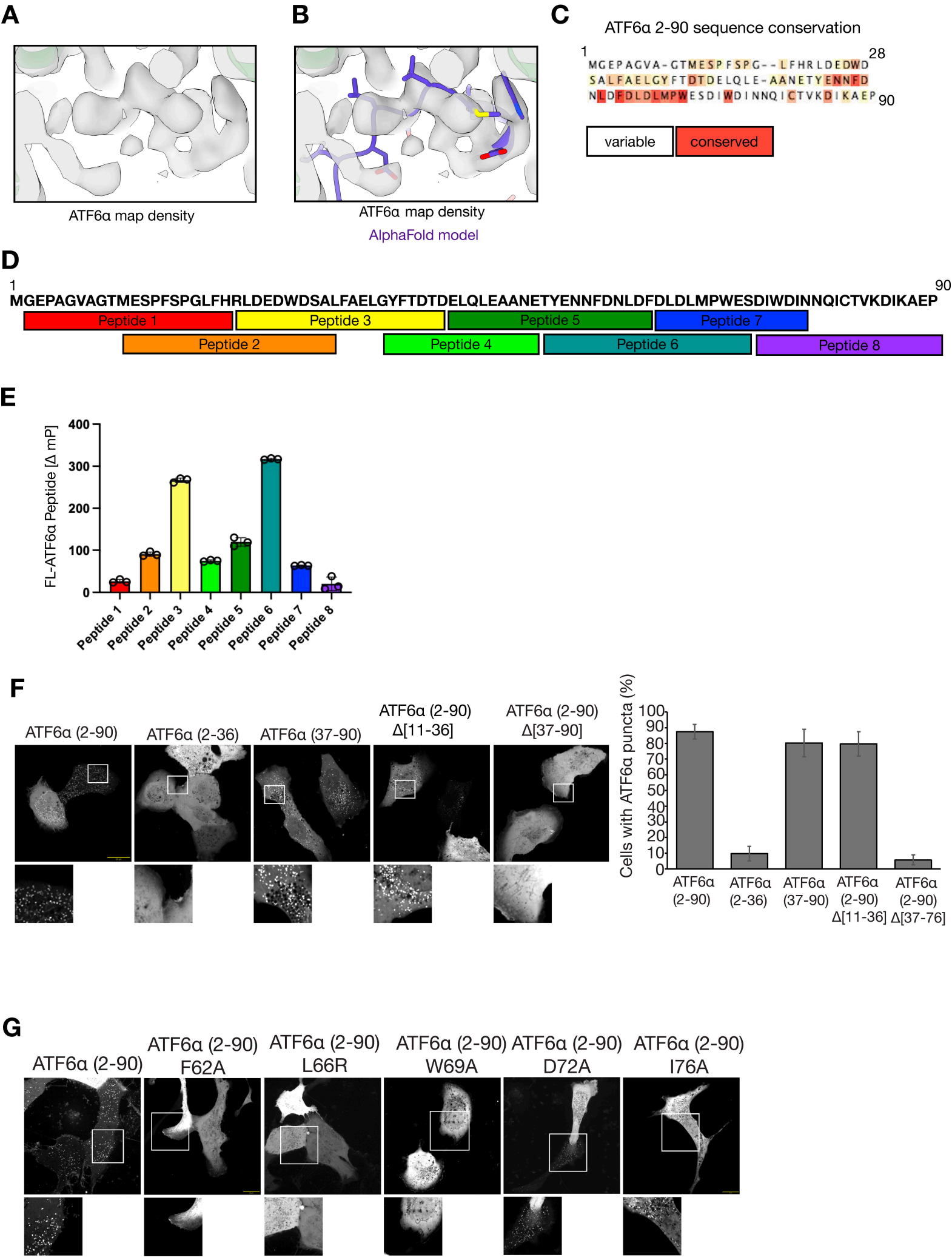

Extended Data Figure 5: Analysis of ATF6α•ABCD3•Ceapin A7 complex

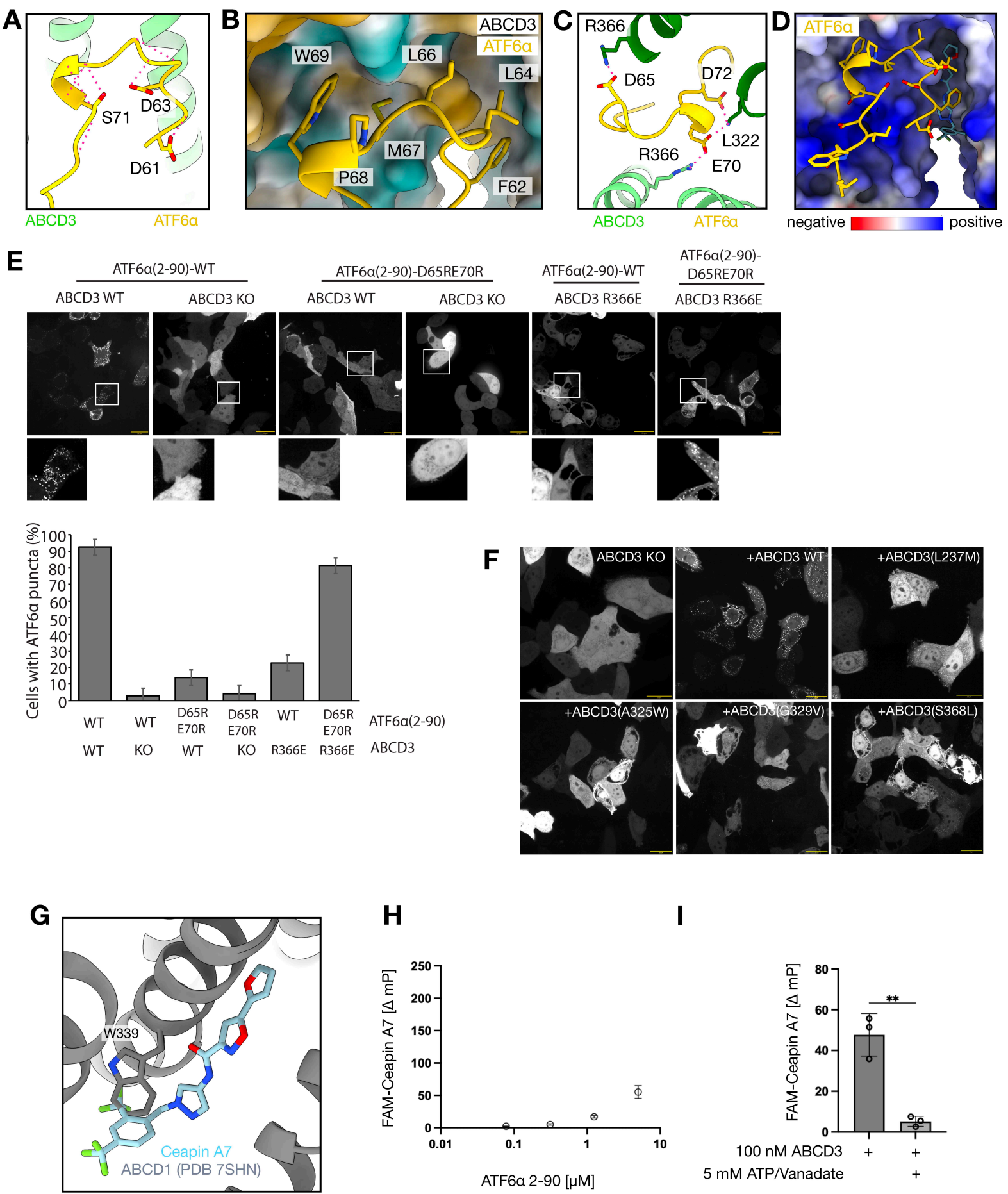

Extended Data Figure 6

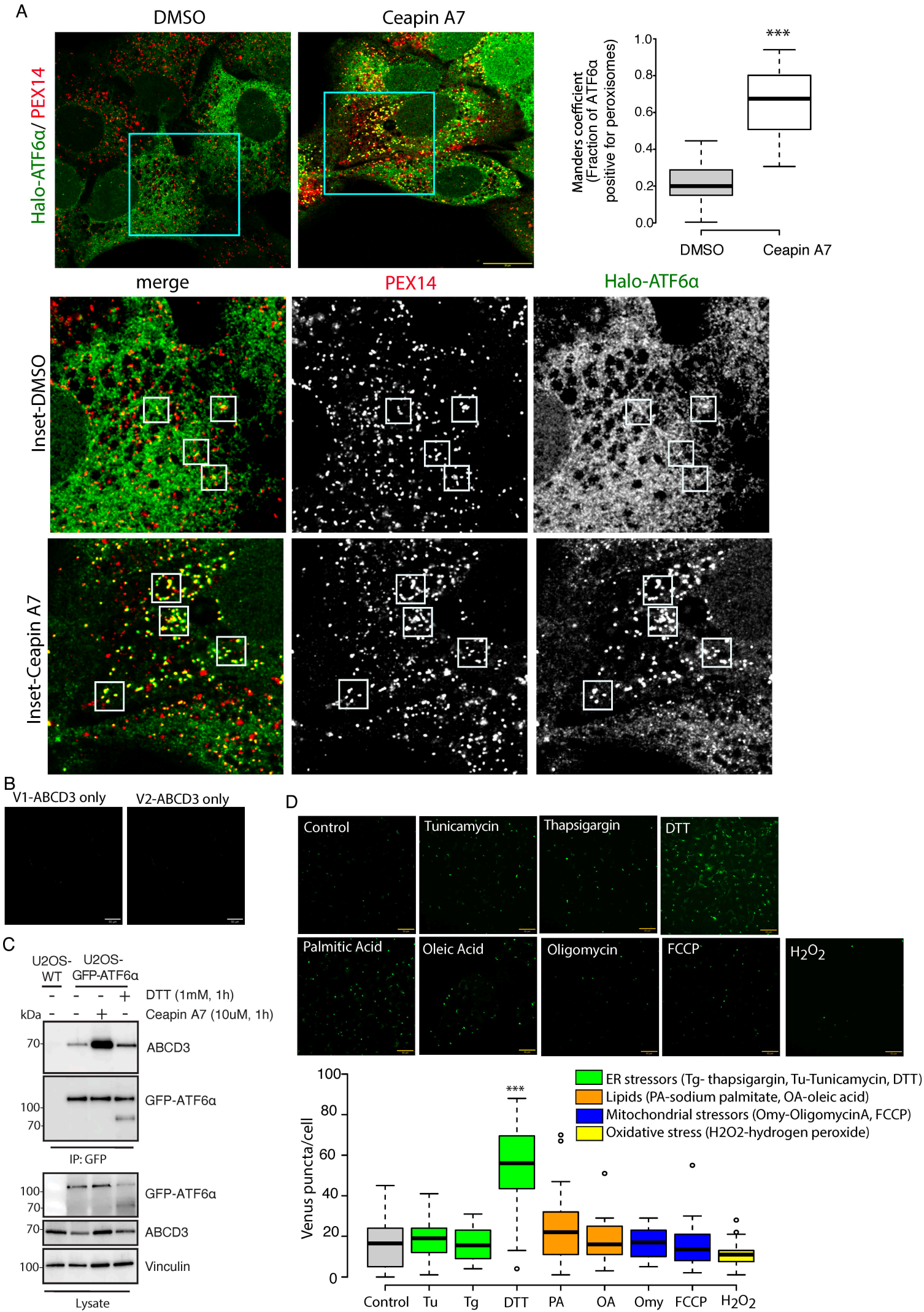

### Extended Data Figure 7

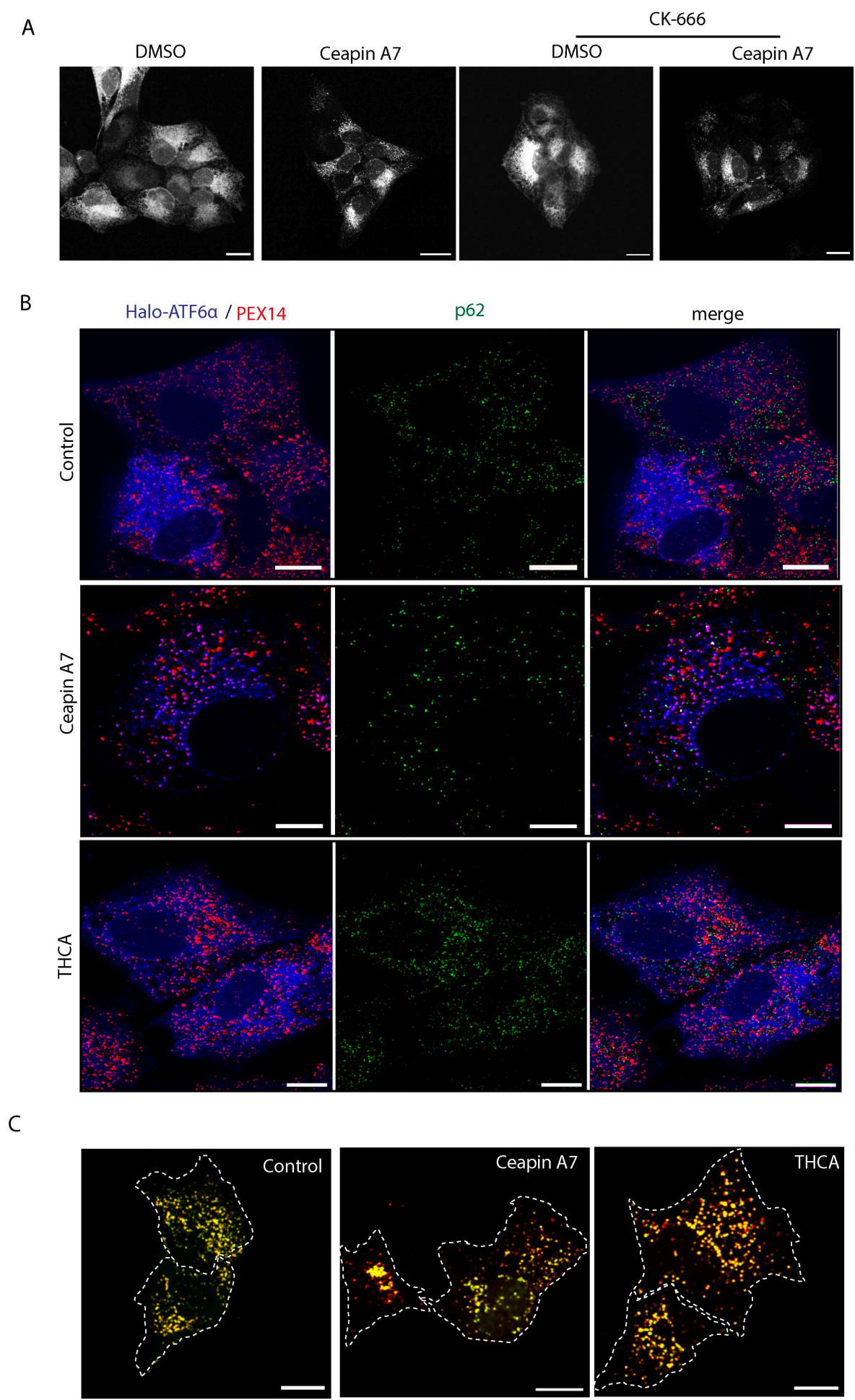

Extended Data Figure 8

A

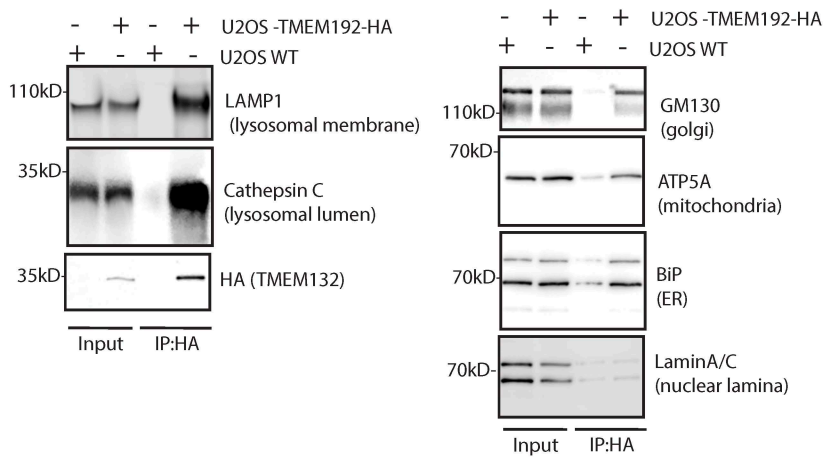

B

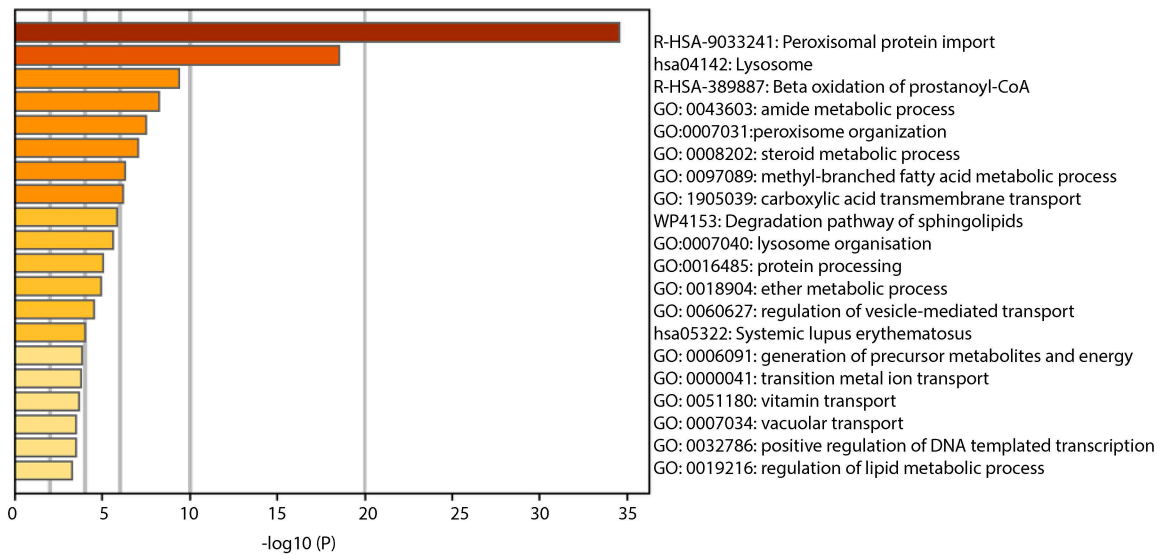

C

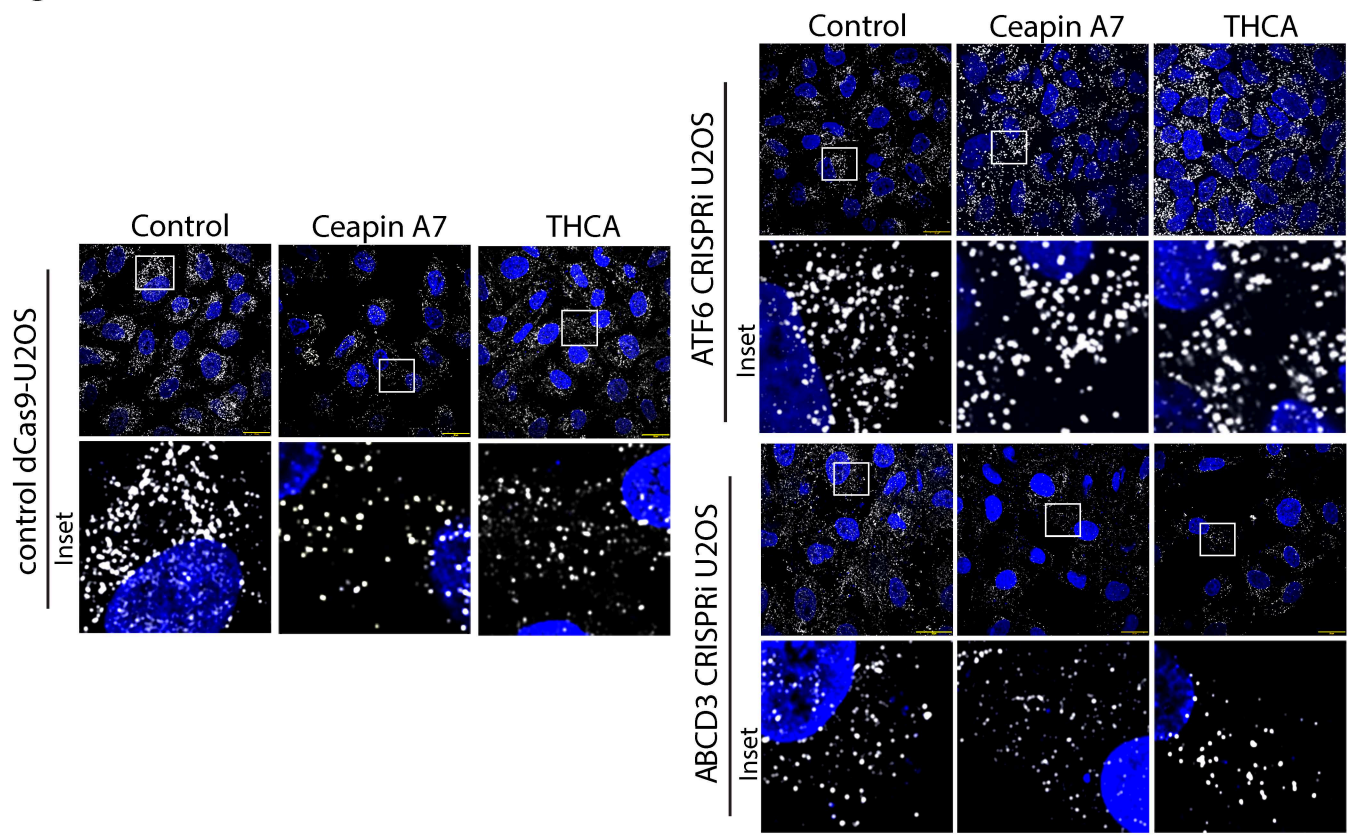

#### Extended Data Figure Legends

##### Extended Data Figure 1: Tomography of GFP-ATF6α puncta

**a**, Top three rows, Z-slice of tomograms collected at sites of GFP-ATF6α signal from Ceapin A7-treated cells ($n = 12$). Scale bar, 200 nm. The two tomograms at the top left are also shown in **Fig. 1C** as representative tomograms. Bottom row, Z-slice of tomograms collected at sites of GFP-ATF6α signal containing peroxisomes from vehicle-treated cells ($n = 4$). PO, peroxisomes. **b**, Measurement of ER-peroxisome distance from tomograms collected on Ceapin A7-treated cells. $n = 35$. **c**, Measurement of peroxisome cross-sectional area measured from tomograms collected on vehicle-treated or Ceapin A7-treated cells.

##### Extended Data Figure 2: Cryo-EM of ATF6α•ABCD3•Ceapin A7 complex

**a**, Left panel, representative size exclusion chromatography (SEC) of purified, detergent-extracted 1XFlag-ABCD3. Right, Coomassie-stained SDS-PAGE of SEC fractions corresponding to SEC trace in left panel. Peak corresponding to fractions B10-C4 were collected and used for structural analysis. **b**, Cryo-EM data processing pipeline for the ATF6α•ABCD3•Ceapin A7 complex performed in CryoSparc. Three separate datasets were pooled during processing. Representative micrograph and 2D classes are shown (top right). Final ATF6α•ABCD3•Ceapin A7 density map is shown with local resolution estimates. Gold-standard Fourier shell correlation (GSFSC) curves of the final 3D reconstruction are shown with FSC value of 0.143 (bottom right).

##### Extended Data Figure 3: Cryo-EM of ABCD3•Ceapin A7 complex

**a**, Cryo-EM data processing pipeline for the ABCD3•Ceapin A7 complex performed in CryoSparc. Representative micrograph and 2D classes are shown (top right). Final ABCD3•Ceapin A7 density map is shown with local resolution estimates. Gold-standard Fourier shell correlation (GSFSC) curves of the final 3D reconstruction are shown with FSC value of 0.143 (bottom right). **b**, Left, 3.1 Å cryo-EM map of the ABCD3•Ceapin A7 complex. Black outline is an overlay of a 20 Å lowpass-filtered map showing the boundary of the detergent micelle. Dashed black line indicates where the peroxisome membrane would surround ABCD3. Middle, atomic model of the ABCD3•Ceapin A7 complex corresponding to cryo-EM map. ABCD3 promoter A (light green), ABCD3 promoter B (dark green), and Ceapin A7 (blue). Color scheme is maintained throughout. Nucleotide binding domains (NBDs) of ABCD3 are indicated. Right, 90-degree rotation from middle and central slice through structure to show the inside of the ABCD3 chamber containing the two symmetrically bound Ceapin A7 molecules (gray box). Right panel, detailed view of the corresponding gray box showing the Ceapin A7 binding pocket. ABCD3 residues interacting with Ceapin A7 are shown. Interactions are marked as follows: contact site (dotted black line), backbone halogen bond (dotted purple line), and π-stacking (dotted orange line). **c**, Overlay of the apo-ABCD3 structure (brown) (PDB: 9NGM) and the ABCD3•Ceapin A7 structure (green). **d**, Overlay of phytanoyl-CoA bound ABCD3 structure (tan) (PDB: 8Z9X) and the ABCD3•Ceapin A7 structure (green).

##### Extended Data Figure 4: Assigning the ATF6α register

**a**, Cryo-EM density map for the ATF6α•ABCD3•Ceapin A7 complex, focused on the ATF6α map density. **b**, AlphaFold modeling was performed on the ATF6α (2-90)•ABCD3 complex and the top ranked result was rigidly fit into ATF6α•ABCD3•Ceapin A7 map density. ATF6α from the AlphaFold model (purple) fit fairly well with the density map. **c**, ATF6α (2-90) sequence conservation calculated from sequence alignments with human ATF6α (2-90) and homologs from six different species. Uniprot accession codes for sequences in the alignment: P18850 (*H. sapiens*), Q20435 (*C. elegans*), G3V909 (*R. norvegicus*), F6VAN0 (*M. musculus*), Q7K2V1 (*D. melanogaster*), H2Q0H3 (*P. troglodytes*), H0VGM7 (*C. porcellus*). Red highlight is more conserved, white highlight is variable. Alignments and conservation made using Clustal Omega and JalView. **d**, ATF6α peptides tiled across amino acids 2-90 and used for binding studies with ABCD3. **e**, Fluorescence polarization of 10 nM fluorescent ATF6α peptide (corresponding to panel **d**) binding to 100 nM purified, full-length ABCD3 *in vitro*. Graph shows mean ± SD, n = 3 replicates per condition. **f**, Live-cell microscopy images of GFP-ATF6α (2-90) and truncation mutants expressed in U2OS cells. All cells were treated with 10 μM Ceapin A7. GFP-ATF6α puncta indicate positive Ceapin A7 induced ATF6α•ABCD3 complex formation. Right, quantification of percentage of cells with GFP-ATF6α puncta for each condition. **g**, Live-cell microscopy images of GFP-ATF6α (2-90) and point mutants expressed in U2OS cells. All cells were treated with 10 μM Ceapin A7. GFP-ATF6α puncta indicate positive Ceapin A7 induced ATF6α•ABCD3 complex formation.

###### Extended Data Figure 5: Analysis of the ATF6α•ABCD3•Ceapin A7 complex

**a**, View of the ATF6α•ABCD3•Ceapin A7 complex focused on the ATF6α peptide (yellow). Dashed pink lines indicate intra-molecular hydrogen bonds, stabilizing the tight turn of ATF6α within the ABCD3 transmembrane cavity. **b**, View of the ATF6α•ABCD3•Ceapin A7 focused on ATF6α (yellow) binding the ABCD3 transmembrane cavity (surface view, colored by hydrophobicity). **c**, View of the ATF6α•ABCD3•Ceapin A7 complex focused on the key salt bridges (pink dashes) formed between ATF6α (yellow) and ABCD3 (light and dark green). **d**, View of the ATF6α•ABCD3•Ceapin A7 focused on ATF6α (yellow) binding the ABCD3 transmembrane cavity (surface view, colored by Coulombic electrostatic potential. Ceapin A7 (light blue). **e**, Live-cell microscopy images of GFP-ATF6α (2-90) point mutants expressed in HeLa cells. The GFP-ATF6α (2-90) point mutants were designed to disrupt key salt bridges stabilizing the ATF6α•ABCD3 interaction. GFP-ATF6α (2-90) were expressed in wildtype HeLa cells and ABCD3 knockout HeLa cells as a control. All cells were treated with 10 μM Ceapin A7. GFP-ATF6α puncta indicate positive Ceapin A7 induced ATF6α•ABCD3 complex formation. Bottom, quantification of percentage of cells with GFP-ATF6α puncta for each condition. **f**, Live-cell microscopy images of GFP-ATF6α (2-90) expressed in HeLa cells with various ABCD3 point mutants expressed on an ABCD3 knockout background. All cells were treated with 10 μM Ceapin A7. GFP-ATF6α puncta indicate positive Ceapin A7 induced ATF6α•ABCD3 complex formation. **g**, Structure of ABCD1 (gray) (PDB: 7SHN) overlaid with the Ceapin A7-bound ABCD3 (Ceapin, blue) structure to indicate where ABCD1 Trp339 sterically clashes with Ceapin A7. **h**, Fluorescence polarization of 10 nM fluorescent Ceapin A7 (FAM-Ceapin A7) binding to purified ATF6α (2-90) *in vitro*. Graph shows mean ± SD, n = 3 replicates per condition. **i**, Fluorescence polarization of 10 nM fluorescent Ceapin A7 (FAM-Ceapin A7) binding to 100 nM purified, full-length ABCD3 *in vitro* with or without 5 mM ATP and vanadate. Graph shows mean ± SD, n = 3 replicates per condition. Statistical significance was evaluated with an unpaired two-tailed *t*-test. \*$P \leq 0.05$, \*\*$P \leq 0.01$, \*\*\*$P \leq 0.001$, \*\*\*\*$P \leq 0.0001$.

###### Extended Data Figure 6

**a**, Immunofluorescence staining of endogenously tagged Halo-ATF6α (green) and PEX14 (red) in U2OS cells treated with DMSO (control) or Ceapin A7. Mander's coefficient to measure fraction of Halo-ATF6α colocalizing with PEX14 is shown for each condition. Data represents 3 independent experiments. Statistical significance was determined using a two-tailed Student's *t*-test. \*\*\*$P < 0.001$. Scale bar, 10 μm. Blue squares indicate zoomed views shown in insets, white boxes indicate colocalization between ER and peroxisomes. **b**, Microscopy images of cells expressing either Venus(fragment V1)-ABCD3 or Venus(fragment V2)-ATF6α showing no fluorescence is detected when only one of the constructs is expressed. **c**, Immunoprecipitation of GFP-ATF6α from U2OS cells treated with 1 mM DTT (4 hours) or 10 μM Ceapin A7 (1 hour) followed by immunoblotting for ABCD3, showing that the ATF6α•ABCD3 interaction strongly increases with Ceapin A7 treatment, but also moderately increases with DTT treatment. The image is the same as shown in **Fig. 1D**. **d**, Quantification of Venus puncta per cell following treatment with ER stressors (tunicamycin, thapsigargin, and DTT), lipids (palmitic acid, oleic acid), mitochondrial inhibitors (oligomycin, FCCP), or oxidative stressors (H<sub>2</sub>O<sub>2</sub>) showing that only DTT treatment enhances ATF6α–ABCD3 interaction. Center lines indicate medians; box limits represent the 25th and 75th percentiles; whiskers extend 1.5× the interquartile range; outliers are shown as individual points. Data represents 3 independent experiments. Statistical significance was determined using a two-tailed Student's *t*-test. \*\*$P \leq 0.01$, \*\*\*$P < 0.001$. Scale bar, 10 μm.

###### Extended Data Figure 7

**a**, GFP-ATF6α distribution in cells treated with Ceapin A7 and CK-666 for 2h. **b**, Immunofluorescence staining of endogenously tagged Halo-ATF6α (shown in blue), PEX14 (red), and p62 (green) in U2OS cells, treated with Ceapin A7 or THCA for 16h. These are the split channels corresponding to microscopy images shown in **Fig. 5D**. Scale bar, 5 μm. **c**, Zoomed out images corresponding to **Fig. 5E**. Tandem fluorescent pexophagy reporter (GFP–SKL/mCherry–SKL) in cells treated with Ceapin A7 or THCA for 16h. Red puncta indicate peroxisomes delivered to lysosomes, while yellow puncta represent healthy peroxisomes.

###### Extended Data Figure 8

**a**, Immunoprecipitation of 3XHA-TMEM192 to isolate lysosomes (Lyso-IP) and immunoblot of lysosomal proteins (LAMP1, lysosomal membrane; Cathepsin C, lysosomal lumen) or Golgi resident proteins (GM130), mitochondrial proteins (ATP51), ER proteins (BiP), or nuclear proteins (Lamin A/C), showing enrichment of lysosomes. **b**, GO term analysis from lysosomal IP-MS upon Ceapin A7 treatment. Peroxisomal processes are the most enriched pathways. **c**, Immunofluorescence staining of PEX14 labelled peroxisomes (white) in cells showing reduced peroxisome number with Ceapin A7 or THCA treatment for 16h. The nucleus is marked with DAPI staining (blue). sgRNA depletion of ATF6α (bottom left) or ABCD3 (bottom right) abolishes this effect.

**Table 1: Cryo-EM data collection, refinement and validation statistics**

|  | ATF6α:ABCD3:Ceapin A7<br>(EMD-79063)<br>(PDB 38QS) | ABCD3:Ceapin A7<br>(EMD-79064)<br>(PDB 38QT) |
| --- | --- | --- |
| **Data collection and processing** |  |  |
| Magnification | 165,000x | 130,000x |
| Voltage (kV) | 300 | 200 |
| Electron exposure (e-/Å<sup>2</sup>) | 60 | 60 |
| Defocus range (μm) | 0.5 – 2.2 | 0.5 – 2.5 |
| Pixel size (Å) | 0.728 | 0.8936 |
| Symmetry imposed | C1 | C2 |
| Initial particle images (no.) | 4,433,880 | 2,783,203 |
| Final particle images (no.) | 134,334 | 158,549 |
| Map resolution (Å) | 2.93 | 3.10 |
| FSC threshold | 0.143 | 0.143 |
| Map resolution range (Å) | 1.87 – 3.87 | 1.93 – 6.88 |
| **Refinement** |  |  |
| Initial model used | AlphaFold2 | AlphaFold2 |
| Model resolution (Å) | 3.10 | 3.33 |
| FSC threshold | 0.5 | 0.5 |
| Map sharpening *B* factor (Å<sup>2</sup>) | -70 or -83.96 | -60 or -104.62 |
| Model composition |  |  |
| Non-hydrogen atoms | 9576 | 9434 |
| Protein residues | 1170 | 1154 |
| Ligands | 2 | 2 |
| *B* factors (Å<sup>2</sup>) (min, medium, max) |  |  |
| Protein | ABCD3/A- 41.8; 104.4; 187.4<br>ABCD3/B- 41.9; 105.7; 186.4<br>ATF6α/C- 71.8; 92.0; 125.0 | ABCD3/A- 39.2; 93.8; 156.8<br>ABCD3/B- 39.2; 93.8; 157.5 |
| Ligand | Ceapin A7/X- 60.2; 66.9; 75.4<br>Ceapin A7/Y- 66.4; 74.8; 82.7<br>All- 41.8; 104.2; 187.4 | Ceapin A7/X- 56.6; 68.5; 72.6<br>Ceapin A7/Y- 56.4; 69.1; 73.3<br>All- 39.2; 93.6; 157.5 |
| R.m.s. deviations |  |  |
| Bond lengths (Å) | 0.0088 | 0.0080 |
| Bond angles (°) | 1.57 | 1.33 |
| Validation |  |  |
| MolProbity score | 0.75 | 0.678 |
| Clashscore | 0.78 | 0.95 |
| Poor rotamers (%) | 0.58 | 0.20 % |
| Ramachandran plot |  |  |
| Favored (%) | 98.10% | 98.34 % |
| Allowed (%) | 1.72% | 1.48% |
| Disallowed (%) | 0.17% | 0.17 % |
