## Supplementary Methods for "Peroxisome quality control is gated by a dynamic ATF6α–ABCD3 tether"

##### **This PDF file includes:**

Materials and Methods

References for Materials and Methods

### Materials and Methods

#### Cell culture

Human bone osteosarcoma (U2OS) cells (ATCC HTB-96) were obtained from the American Type Culture Collection. All cells were grown in DMEM with high glucose (Sigma D5796) supplemented with 10% FBS (Life technologies # 10082147), 2 mM L-glutamine (Sigma G2150). Cells were incubated at 37°C, 5% CO<sub>2</sub>. For cells expressing GFP-ATF6α, growth media was supplemented with 500 µg/mL G418 (Roche 04 727 878 001). All cell lines tested negative for mycoplasma contamination.

#### Plasmids and Generation of Stable Cell lines

U2OS cells stably expressing GFP-ATF6α were purchased from Thermo Fisher Scientific (Cat. No. 084\_01).

The tandem fluorescence pexophagy reporter was generated by VectorBuilder by cloning the coding sequence of mCherry-GFP fused to a C-terminal type 1 peroxisomal targeting signal (SKL) into the lentiviral vector pLV[Exp]-Puro-CMV. The mCherry-GFP coding sequence was derived from Addgene plasmid #170446.

The peroxisomal redox reporter HyPer3-SKL was generated in the same pLV[Exp]-Puro-CMV lentiviral backbone. The HyPer3 coding sequence was obtained from Addgene plasmid #42131 and fused to a C-terminal SKL targeting sequence.

For split-Venus interaction studies, the N-terminal fragment of Venus (amino acids 1–211; VN) was fused to the C terminus of ABCD3, while the C-terminal fragment of Venus (amino acids 212–239; VC) was fused to the N terminus of ATF6α. Both constructs were synthesized and cloned by VectorBuilder into the lentiviral vector pLV[Exp]-CMV. The ABCD3-VN construct contained a puromycin resistance cassette, whereas the VC-ATF6α construct contained a hygromycin resistance cassette to enable sequential selection.

For lysosome immunoprecipitation experiments, the TMEM192-3×HA construct (Addgene plasmid #102930) was used.

For generation of stable cell lines, lentiviral particles were produced in Lenti-X HEK293T cells by transient transfection of the transfer plasmid together with the packaging plasmids psPAX2 (Addgene #12260) and pMD2.G (Addgene #12259) using the VirusGEN® LV Transfection Kit (Mirus Bio # MIR6760) according to the manufacturer's instructions. Viral supernatants were collected 48 h after transfection, filtered through a 0.45-µm filter, and used to transduce target cells in the presence of 8 µg/ml polybrene. 100 µl of lentiviral supernatant was added for transduction of U2OS cells per well of a 6-well plate. Culture medium was replaced 24 h after transduction, and cells were subsequently selected with 1 µg/ml puromycin and/or 200 µg/ml hygromycin B for 7 days. Following antibiotic selection, fluorescent cells were enriched by fluorescence-activated cell sorting (FACS) to generate stable polyclonal populations.

#### Generation of HA-Halo-ATF6 U2OS cell line

The guide RNA (CACCGGAGAAGGAACTTGTGAAATG; AAACCATTTCACAAGTTCCTTCTCC) was designed using the online tool Benchling (https://www.benchling.com), procured as annealed oligonucleotides from IDT, and cloned into the SpCas9 px330 plasmid (Addgene plasmid #42230). The HA-Halo-ATF6<sup>EN</sup> homology repair (HR) plasmid was designed with 0.6-kb homology arms and synthesized by Twist Bioscience. A glycine-serine (GS) linker and BamHI and EcoRI sites were added between the two homology arms to allow insertion or replacement of tags in future experiments. The coding sequence of the

HaloTag, including a small epitope tag (HA-tag), was integrated between the homology arms. The PAM sequence of the guide was mutated in the homology arms to prevent re-editing of the plasmid template once inside cells.

To generate U2OS endogenous knock-in cell lines, $2 \times 10^5$ U2OS cells were washed with PBS and resuspended in 20 μL of Lonza SE Cell Line solution (Lonza # V4XC-1032), mixed with 1 μg of plasmid DNA (split evenly between the guide and HR plasmids) and 0.69 μL of 0.69 mM HDR enhancer solution (IDT #10007910). The reaction mix was transferred into one well of a 16-well Nucleocuvette® Strip (provided with the Lonza kit) and nucleofected using the Lonza 4D-Nucleofector® (program CM-104). Cells were loaded with 200nM Janelia Fluor® 646 HaloTag® Ligand (Promega #GA1120) prior to sorting them on the BD FACS Aria™ Fusion Flow Cytometer, using WT cells for negative gating. Cells with high fluorescence for Halotag were expanded. Knock-in was confirmed by amplifying the target locus from total DNA by PCR (forward primer: 5'-CGCCTTCCCCATAAAACAGC-3'; reverse primer: 5'-AGATCACCCGGTACTTCCCC-3'). The resulting amplicons were Sanger sequenced (Quintara Biosciences), and knock-in was confirmed by BLAST analysis. Finally, the expression of HA-Halo-ATF6 was confirmed by western blotting using antibodies against HA and HaloTag.

#### Protein expression and purification

For *in vitro* pulldown experiments, human ABCD3 (Uniprot ID: P28288) was cloned in pFastBac1 vector with a cleavable C-terminal 10XHis tag and eGFP for baculovirus-mediated expression in Gibco Sf9 cells (Thermo Scientific #11496015) derived from *Spodoptera frugiperda*. To prepare recombinant Bacmid, baculovirus multiplicity of infection (MOI) of 1:1000 (v/v) in 0.5 L Sf9 cells for 72 hours at 27°C. The cells were harvested by centrifugation at 2000 rpm, 5 min, 4°C and then resuspended in Lysis Buffer (50 mM Tris pH 7.5, 100 mM NaCl, 100 mM MgCl<sub>2</sub>, 10% glycerol) supplemented with protease inhibitor cocktail (Roche). Cells were incubated on ice for 30 minutes and then lysed by sonication and centrifuged at 5000 x g, 4°C, for 15 minutes. The supernatant was collected and centrifuged at 100,000 x g, 4°C for 60 min. The pellet was solubilized in 15 mL of Lysis Buffer containing 300 mM NaCl and 1% w/v lauryl maltose neopentyl glycol (LMNG) (Anatrace): 0.1% w/v cholesteryl hemisuccinate (CHS) (Anatrace) overnight. Then it was clarified by centrifugation at 20,300 x g for 30 minutes and the supernatant was collected. Ni-NTA resin was pre-equilibrated with Binding Buffer (50 mM Tris pH 7.5, 300 mM NaCl, 100 mM MgCl<sub>2</sub>, 10% glycerol, 0.02% GDN), and then incubated with the supernatant (supplemented with 5 mM imidazole) for 2 hours at 4°C on a rotator. The resin was washed in the column with Binding Buffer, Wash Buffer (50 mM Tris pH 7.5, 300 mM NaCl, 100 mM MgCl<sub>2</sub>, 10% glycerol, 0.02% GDN, and 10 mM imidazole). Protein was eluted with 250 mM imidazole and then concentrated in an ultra-filtration tube. Protein was further purified by size exclusion chromatography on a Superdex 200 increase column (GE) with Storage Buffer (20 mM HEPES pH 7.5, 300 mM NaCl, 2 mM MgCl<sub>2</sub>, 2% glycerol, 0.02% GDN). The later half of the major peak was collected (**Extended Data Fig. 2A**). The protein was concentrated in an ultra-filtration tube, flash frozen, and stored at -80°C.

For *in vitro* binding experiments and ATPase assays, human ABCD3 (Uniprot ID: P28288) was cloned in pFastBac1 vector with a N-terminal 1XFlag tag for baculovirus-mediated expression in Gibco High Five cells (Thermo Scientific, #B85502) derived from *Trichoplusia ni*. Protein was purified as above, except with Anti-DYKDDDDK Affinity Resin instead of Ni-NTA. Resin was pre-equilibrated with Binding Buffer (50 mM Tris pH 7.5, 300 mM NaCl, 100 mM MgCl<sub>2</sub>, 10%

glycerol, 0.02% GDN), and then incubated with the supernatant for 2 hours at 4°C on a rotator. The resin was washed in the column with Binding Buffer, ATP Wash Buffer (50 mM Tris pH 7.5, 300 mM NaCl, 100 mM MgCl<sub>2</sub>, 5 mM ATP, 10% glycerol, and 0.02% GDN), then Binding Buffer again. Protein was eluted with 250 ug/ml 3X DYKDDDDK Peptide (Pierce) and then concentrated in an ultra-filtration tube. Protein was further purified by size exclusion chromatograph on a Superdex 200 increase column (GE) with Storage Buffer (20 mM HEPES pH 7.5, 150 mM NaCl, 2% glycerol, 0.02% GDN). The protein was concentrated in an ultra-filtration tube, flash frozen, and stored at -80°C.

For cryo-EM, human ABCD3 (Uniprot ID: P28288) was cloned by Twist into the pTwist CMV BG WPRE Neo plasmid with a N-terminal 1XFlag tag for transient mammalian expression in Expi293F cells (Gibco, #A14527). 0.8 L Expi293F cells were transfected according to Expi293 protocol (Gibco) and grown at 37°C for 48 hours. The cells were harvested by centrifugation and resuspended in 60 mL Lysis Buffer (50 mM Tris pH 7.5, 150 mM NaCl, 20% glycerol, complete EDTA-free protease inhibitor cocktail (Roche)) and then dounce homogenized (30 strokes pestle A, 30 strokes pestle B). 1% w/v lauryl maltose neopentyl glycol (LMNG) (Anatrace): 0.1% w/v cholesteryl hemisuccinate (CHS) (Anatrace) was added to the solution and incubated with stirring for 2 hours at 4°C. Then it was clarified by centrifugation at 111,818 x g for 1 hour at 4°C and the supernatant was collected. Anti-DYKDDDDK Affinity Resin (Pierce) was pre-equilibrated with Lysis Buffer and then incubated with the supernatant at 4°C on a rotator overnight. The resin was washed in the column with Wash Buffer (50 mM Tris pH 7.5, 150 mM NaCl, 10% glycerol, 0.02% GDN). Protein was eluted with 600 ug/mL 3X DYKDDDDK Peptide (Pierce) and then concentrated in an ultra-filtration tube. Protein was further purified by size exclusion chromatograph on a Superdex 10-300 increase column (GE) with Storage Buffer (25 mM HEPES pH 7.5, 150 mM NaCl, 0.02% GDN). The protein was concentrated in an ultra-filtration tube, flash frozen, and stored at -80°C.

Human ATF6α (amino acids 2-90) (Uniprot ID:P18850) was cloned into pET302 with an N-terminal 3xFLAG tag and cleavable C-terminal maltose binding protein and 6xHis tag for expression in BLR (DE3) *E. coli* (Novagen). For pulldown experiments, cells were grown in LB media at 37°C until OD<sub>600</sub> reached 0.7 and then induced with 1 mM IPTG for 3 hours at 25°C. Cells were harvested and resuspended in Lysis Buffer (20 mM HEPES pH 7.0, 100 mM KCl, 10% glycerol, 0.1% Triton x-100, 5 mM BME, and complete EDTA-free protease inhibitor cocktail (Roche)). Cells were lysed by sonication at 4°C and lysate was clarified at 20,000 x g for 30 minutes at 4°C. The clarified supernatant was supplemented with 25 mM imidazole and loaded onto a HisTrap FF (5 mL) column, washed with Binding Buffer (50 mM HEPES pH 7.0, 300 mM NaCl, 10% glycerol, 25 mM imidazole, and 1 mM TCEP). Protein was eluted with 500 mM imidazole. Protein was loaded onto a HiPrep 26/10 desalting column equilibrated with Buffer A (50 mM HEPES pH 7.5, 100 mM NaCl, 10% glycerol, and 1 mM DTT) to remove imidazole. Protein was then purified by anion exchange chromatography using a HiTrap Capto Q column equilibrated with Buffer A. Protein was eluted with a linear gradient from 100 mM to 1 M NaCl. Protein eluted between 400-600 mM NaCl gradient. The protein was concentrated in an ultra-filtration tube, flash frozen, and stored at -80°C.

For cryo-EM and fluorescence polarization assays, human ATF6α (amino acids 2-90) was purified as above, but the plasmid lacked the C-terminal 6xHis tag. Protein was expressed in BL21 (DE3)

*E. coli*, cells were grown in LB media at 37°C until OD600 reached 0.6-0.8 and then induced with 0.5 mM IPTG for 16 hours at 16°C. Cells were harvested and resuspended in Lysis Buffer (30 mM Tris pH 7.5, 400 mM KCl, 10% glycerol, 5 mM BME, and complete EDTA-free protease inhibitor cocktail (Roche)). Cells were lysed using an Emulsiflex-C3 (Avestin) and lysate was clarified at 30,000 x g for 45 minutes at 4°C. Amylose resin (NEB #E8021) was pre-equilibrated with Lysis Buffer and then incubated with the clarified supernatant at 4°C on a rotator for 1 hour. The resin was washed in the column with High Salt Buffer (30 mM Tris pH 7.5, 400 mM KCl, 5% glycerol, 5 mM BME) and then Low Salt Buffer (30 mM Tris pH 7.5, 100 mM KCl, 5% glycerol, 5 mM BME). Protein was eluted with 10 mM maltose and then concentrated in an ultra-filtration tube. For cryo-EM, the C-terminal MBP tag was removed with TEV protease (GenScript) by overnight incubation at 4°C. Protein was then purified by anion exchange chromatography using a MonoQ 10-100 GL (Cytiva) column equilibrated with Buffer A (50 mM Tris pH 7.5, 100 mM KCl, 5% glycerol, 1 mM DTT). Protein was concentrated in an ultra-filtration tube and further purified by SEC using a S200 10-300 GL (Cytiva) column equilibrated with Storage Buffer (50 mM HEPES 7.5, 100 mM KCl, 5% glycerol, 0.5 mM TCEP). The eluted protein was concentrated in an ultra-filtration tube, flash frozen, and stored at -80°C.

#### **CRISPR interference-mediated depletion of ATF6α and ABCD3**

CRISPR interference was performed in U2OS cells stably expressing dCas9-KRAB. Cells were transduced with lentiviral vectors expressing two sgRNAs targeting either ATF6 or ABCD3, or with non-targeting control sgRNAs. The sgRNA sequences were selected from a previously published CRISPRi library<sup>1</sup> and are listed below. Following transduction, cells were maintained under puromycin selection for 5 days to enrich for sgRNA-expressing cells before downstream experiments. Depletion efficiency was confirmed by immunoblotting.

ATF6 sgRNA 1: 5' GCCACGAGTGGGCGGAAGTA 3'

ATF6 sgRNA2: 5' TGGAGTCACCTTTTAGCCCC 3'

ABCD3 sgRNA1: 5' GGTACCAGCGAGCCGGCGAG 3'

ABCD3 sgRNA2: 5' TGGCGGCCTTCAGCAAGTAC 3'

#### **Immunoprecipitation and immunoblotting**

For GFP-ATF6α, Halo-ATF6α immunoprecipitation experiments, cells were treated with vehicle Ceapin A7 or cellular stressors, washed with ice-cold phosphate-buffered saline, and lysed in lysis buffer (50 mM Tris (pH 7.5), 150 mM NaCl, 1% TritonX-100, protease inhibitor cocktail). Cell lysates were clarified by centrifugation at 13000g, 10min and the supernatant was used as input samples for the western blot and immunoprecipitations. Protein content in the samples were quantified by BCA assay (Thermo Scientific, PI23227) following manufacturer's protocol. For co-immunoprecipitation, 1mg total protein was incubated with Chromatek GFP beads (for GFP-ATF6α) or HaloTag affinity resin (for Halo-ATF6α) under gentle rotation at 4 °C for 1 h. Reactions were washed with wash buffer (50mM Tris (pH 7.5), 300mM NaCl, 0.1% TritonX-100, protease inhibitor cocktail), bound proteins were eluted by boiling in SDS sample buffer and separated by SDS-PAGE before transfer onto nitrocellulose membranes. Membranes were blocked with 5% milk in TBS containing 0.1% Tween-20, and incubated with primary antibodies against HA tag (to detect Halo-ATF6α), ABCD3, and other indicated proteins overnight at 4 °C followed by incubation with horseradish peroxidase-conjugated secondary antibodies. Protein bands were visualized using enhanced chemiluminescence and quantified by the Odyssey LiCor Imager. Representative immunoblots from at least three independent biological replicates are shown.

#### 3D correlative cryo-focused ion beam milling and electron tomograph

Au/SiO<sub>2</sub> Quantifoil grids (200 mesh, R2/2) and a 3.5 cm glass-bottom imaging dish were coated with 50 µg/mL collagen in DPBS for 10 minutes at room temperature. After removing the collagen solution from the imaging dish, the coated grids were rinsed by dipping sequentially through six tubes of DPBS and transferred directly into the 3.5 cm dish. U2OS cells stably expressing GFP-ATF6α were seeded into the dish at a density of $2 \times 10^5$ cells/mL (2 mL total volume) and incubated overnight. The following day, cells were treated with 10 µM Ceapin A7 or DMSO for 1 hour. For the final 30 minutes of treatment, cells were incubated with 200 nM Lipid-Blue. Prior to plunge freezing, grids were washed with DPBS to remove excess dye and blotted from the back for 3, 5, or 7 seconds using a Leica EM GP2 plunge freezer before vitrification in liquid ethane.

Sample grids were clipped into Autogrids and loaded into an Arctis cryo-plasma FIB (Thermo Fisher Scientific) for automated lamella preparation. The workflow was executed using the Arctis WebUI (Thermo Fisher Scientific). Grids were coated with an ion-sputtered metallic Pt layer followed by deposition of an organometallic Pt layer via the GIS. Targeted GFP-ATF6α foci were identified using the integrated fluorescence light microscope and subsequently subjected to automated lamella preparation using Xe<sup>+</sup> ions. The final lamella thickness was set to 200 nm for the polishing step. The Lipid Blue channel was imaged for later correlative light and electron microscopy (CLEM) registration within the TEM search map, enabling accurate alignment of tilt series acquisition with the defined region of interest.

Tilt series were acquired on a Krios G4 TEM equipped with a Selectris X energy filter and a Falcon 4i direct electron detector (Thermo Fisher Scientific). Data collection was performed using a dose-symmetric tilt scheme with Tomography 5-v5.22 (Thermo Fisher Scientific). A tilt range of $\pm 60^\circ$ was used with $2^\circ$ increments, starting at $12^\circ$ to compensate for the lamella milling angle. Frames were written in TIFF format at a nominal pixel size of 3.04 Å and an electron dose of approximately $3.4 \text{ e}^-/\text{Å}^2$ per tilt. A 100 µm objective lens aperture and 10 eV energy filter slit was used throughout data collection. CLEM was performed on the low-magnification search maps to target GFP-ATF6 foci while peroxisomes could be readily identified in search maps without the need for a fluorescence marker.

Segmentation was performed on SIRT-filtered and bin 4x tomograms output by TomoLive/CryoFlow-v1.27.0 (Thermo Fisher Scientific). Membranous features were initially segmented using membrain2-v0.0.10<sup>2</sup> using the pre-trained model MemBrain\_seg\_v10\_beta.ckpt and further refined manually using Amira/Avizo-v2024.2 (Thermo Fisher Scientific).

#### *In vitro* ATF6α pulldown

Relating to **Figure 1E**, 4 µM ABCD3-10XHis-eGFP and 2 µM purified 3XFlag-ATF6α (2-90)-MBP-6XHis were incubated with or without 5 mM sodium orthovanadate (NEB), 5 mM ATP (Thermo Scientific #R0441), and 5 mM MgCl<sub>2</sub> or 10 µM Ceapin A7 (Integral Biosciences, solubilized in DMSO) in Reaction Buffer (25 mM HEPES pH 7.5, 100 mM NaCl, 0.02% GDN, 0.5 mM TCEP) for 1 hour at 25°C. Relating to **Figure 3E**, 2 µM ABCD3-10XHis-eGFP and 2 µM purified 3XFlag-ATF6α were incubated in Reaction Buffer with or without 10 µM Ceapin A7 with varying concentrations of phytanoyl-coenzyme A (Avanti #870742P) for 1 hour at 25°C. Amylose resin (NEB) was equilibrated with Reaction Buffer, added to each reaction at a 1:1 ratio, and incubated with each reaction for 1 hour at 4°C with rotation. Resin was washed 3 times with Wash

Buffer (25 mM HEPES pH 7.5, 300 mM NaCl, 0.02% GDN, 0.5 mM TCEP) and eluted with Elution Buffer (25 mM HEPES pH 7.5, 100 mM NaCl, 0.02% GDN, 0.5 mM TCEP, 10 mM maltose). The elution was analyzed by SDS-PAGE on an AnykDa mini-protean gel (BioRad) and stained with Coomassie Instant Blue (Abcam #ISB1L).

##### **ABCD3•Ceapin A7•ATF6α and ABCD3•Ceapin A7 complex preparation for cryo-EM**

For the ABCD3•Ceapin A7•ATF6α complex, 13 μM of 3XFlag-ATF6α (2-90) was incubated with 10 μM 1XFlag-ABCD3 with or without 100 μM Ceapin A7 (Integral Biosciences, solubilized in DMSO) in Reaction Buffer (25 mM HEPES pH 7.5, 150 mM NaCl, 0.02% GDN). Samples were incubated for 30 minutes at 25°C and then concentrated at 4°C in a spin concentrator (ViVaSpin, 100 MWCO) to an approximate final concentration of ~6 mg/mL. 2.5 μL of sample was applied to Quantifoil glow-discharged R1.2/1.3, 300 mesh, Au, 25nm thick holey carbon grid (Quantifoil Micro Tools GmbH) and plunge-frozen in liquid ethane using a Vitrobot Mark IV (FEI) with blot time 4 seconds, blotting force of 0, at 10°C and 100% humidity.

For the ABCD3•Ceapin A7 complex, 7.45 μM 1XFlag-ABCD3 was incubated with 10 μM Ceapin A7 (Integral Biosciences, solubilized in DMSO) for 30 minutes at 25°C and then spin concentrated at 4°C in a spin concentrator (ViVaSpin, 100 MWCO) to an approximate final concentration of ~8 mg/mL. 3 μL of sample was applied to Quantifoil glow-discharged R1.2/1.3, 300 mesh, Au, 25nm thick holey carbon grid (Quantifoil Micro Tools GmbH) and plunge-frozen in liquid ethane using a Leica GP2.

##### **Cryo-EM data acquisition**

For the ABCD3•Ceapin A7•ATF6α complex, images were collected on a Titan Krios G4 transmission electron microscope (Thermo Fisher Scientific) operating at 300 keV. For the ABCD3•Ceapin A7 complex, images were collected on a Glacios transmission electron microscope (Thermo Fisher Scientific) operating at 200 keV. Microscopes were equipped with a Falcon 4i detector and SelectrisX (Krios) or SelectrisEF (Glacios) energy filter with a slit width of 10 eV. Data was acquired using Thermo Scientific EPU software (3.6.0.6389). The microscope and data collection parameters for each dataset are provided in **Table 1**.

##### **Cryo-EM data processing**

Specific details of data analysis are presented in **Extended Data Figure 2B and 3A** for the ABCD3•Ceapin A7•ATF6α and ABCD3•Ceapin A7 datasets, respectively. For both datasets, patch-based global and local motion correction, and patch CTF estimation were performed within CryoSPARC live (Structura Biotechnology) using default settings. Image analysis was carried out using CryoSPARC v4.7.1. For ABCD3•Ceapin A7•ATF6α dataset, three separate datasets were collected and combined for 18,200 total micrographs. For ABCD3•Ceapin A7, one dataset with 5,500 micrographs was collected. Micrographs were curated based on CTF fit resolution and particles were picked based on template ABCD3 structure (PDB ID:8Z9X)<sup>3</sup>. Particles were extracted at box size of 360 pixels and Fourier cropped for initial processing. 2D class averages were performed and particles from the best classes were selected for iterative rounds of Non-uniform 3D refinement, heterogeneous 3D refinement, and 3D classification at C2, selecting classes with the best particles for further refinement throughout. Reference based motion correction was performed and local and global CTF parameters were refined. A final round of 3D classification was performed to select the final particle stack and these particles were subjected to a final local refinement. For the ABCD3•Ceapin A7•ATF6α dataset, iterative rounds of 3D refinement and 3D classification were performed at C1 to resolve the apparent asymmetry of the ATF6α density. The nominal resolution for final reconstructions of ABCD3•Ceapin A7•ATF6α and ABCD3•Ceapin A7 were determined by the gold standard Fourier shell correlation (FSC) using the 0.143 criterion. Maps were sharpened and filtered automatically as determined by CryoSparc according to an estimated overall map B-factor. CryoSparc was used to estimate the local resolution of each map and displayed in ChimeraX.

#### Model building and refinement

The initial ABCD3•Ceapin A7•ATF6α model was generated by AlphaFold2 prediction using the full-length human ABCD3 amino acid sequence and the human ATF6α sequence from amino acids 2-90. The top model was selected based on predicted aligned error score and visual inspection of map-to-model fit. The model was globally relaxed into the map using ISOLDE within UCSF ChimeraX. The Ceapin A7 SMILES file based on the chemical structure was used to generate restraints using Acedrg within the CCP4 program suite (v9.0.12), and fit manually into the observed cryo-EM density using ISOLDE. Each residue was manually refined to fit the experimental map. The ABCD3•Ceapin A7•ATF6α model was then used as a starting point for ABCD3•Ceapin A7 model for subsequent global relaxation and per-residue refinement in ISOLDE. A final refinement of all models was carried out against half-maps using Servalcat. All figures including cryo-EM maps and models were created using UCSF ChimeraX v.1.8.

#### Assignment of ATF6 amino acids 61-76 to the density observed in the ATF6(2-90)•ABCD3•Ceapin A7 cryo-EM structure.

To model the ATF6 sequence into the density of the ABCD3•Ceapin A7•ATF6(2-90) cryo-EM structure and validate the resulting model, we combined information resulting from multiple lines of evidence and approaches:

1. By visual inspection, we see a strand of density within the ABCD3 chamber for ~7 residues at high resolution, including well-resolved side chain density (**Extended Data Fig. 4A**). We made an initial assignment of the register of this 7 amino acid stretch with AlphaFold using rigid-body docking (**Extended Data Fig. 4B**). The model predicted a fit of ATF6α(64-70) into the density.
2. Phylogenetic analysis revealed that the ATF6α(62-69) region is highly conserved, indicating the region's potential to participate in a functionally important interaction (**Extended Data Fig. 4C**).
3. We performed *in vitro* fluorescence polarization anisotropy binding studies with fluorescent peptides tiled across the ATF6α 2-90 region (**Extended Data Fig. 4D**). To this end, we mixed each peptide with purified ABCD3 and measured equilibrium binding. The assay identified two fragments that bound ABCD3 with highest affinity (ATF6α(22-41) and ATF6α(51-70)) (**Extended Data Fig. 4E**).
4. Binding of various ATF6α truncation mutants *in cellula* using the ATF6α puncta assay (**Extended Data Fig. 4F**). We found i) that the ATF6α (37-90) region was sufficient for binding to ABCD3 and ii) that the ATF6α (37-76) region was necessary for binding. By contrast, the ATF6α (2-36) region was not sufficient, and the ATF6α (11-36) region was not necessary. Thus, the necessary and sufficient binding site lies within the ATF6α (37-76) region, matching the AlphaFold model.

5. Point mutation of ATF6 amino acids F62, L66, and W69 with side chains that in our model are predicted to interact with ABCD3 abolish binding *in cellula*, as shown by the ATF6α puncta and nanoBret assays. By contrast, point mutations of ATF6α mutant of amino acids D72 and I76 that in our model are predicted not to interact with ABCD3 do not abolish binding to ABCD3 *in cellula* (**Extended Data Fig. 4G**).

#### Fluorescence polarization assays

Relating to **Figure 3A, 3B** and **Extended Data Figure 5H, 5I**, fluorescence polarization was performed using fluorescent Ceapin A7 synthesized with a 6-carboxyfluorescein moiety (FAM-Ceapin A7, Integral Biosciences) in a 384-well, non-binding, low volume black plate (Corning #3575). Polarization anisotropy was measured using a CLARIOstar Plus plate reader (BMG Labtech) with excitation/emission wavelengths of 482 nm/520 nm and temperature control set at 25°C. For **Figure 3A**, various concentrations of purified 1xFlag-ABCD3 were mixed with 10 nM FAM-Ceapin A7 in ABCD3 Reaction Buffer (25 mM HEPES pH 7.5, 100 mM NaCl, 0.02% GDN, 0.5 mM TCEP). For **Figure 3B**, 1 μM purified 1xFlag-ABCD3 was mixed with 10 nM FAM-Ceapin A7 and various concentrations of phytanoyl-coenzyme A (Avanti) in ABCD3 Reaction Buffer. For **Extended Data Figure 5H**, various concentrations of 3XFlag-ATF6α (2-90)-MBP were mixed with 10 nM FAM-Ceapin A7 in ATF6 Reaction Buffer (50 mM Hepes pH 7.5, 100 mM KCl, 5% glycerol, 0.5 mM TCEP). For **Extended Data Figure 5I**, 100 nM purified 1XFlag-ABCD3 was mixed with 10 nM FAM-Ceapin A7 with or without 5 mM sodium orthovanadate and 5 mM ATP-MgCl<sub>2</sub> in ABCD3 Reaction Buffer.

Relating to **Figure 2F, 3G**, and **Extended Data Figure 4E**, fluorescent peptides corresponding to specific short regions of ATF6α were synthesized by GenScript with an N-terminal fluorophore (either 5-carboxyfluorescein or fluorescein). Fluorescence polarization assays were performed as described above. For **Figure 2F**, various concentrations of purified 1xFlag-ABCD3 were mixed with 0.1 nM FAM-ATF6α (61-76) and 25 μM Ceapin A7 (Integral Biosciences) or DMSO in ABCD3 Reaction Buffer. Notably, the Hill coefficient of FAM-ATF6α (61-76) binding to ABCD3 is $> 2$, indicating ATF6α may promote ABCD3 dimerization. For **Figure 3G**, 4 nM purified 1xFlag-ABCD3 was mixed with 0.1 nM FAM-ATF6α (61-76) and 25 μM Ceapin A7, phytanoyl-CoA (Avanti), CoA-hybrid 1, CoA-hybrid 2, or DMSO control in ABCD3 Reaction Buffer. For **Extended Data Figure 4E**, 10 nM ATF6α peptide 1-8 were mixed with 50 nM purified 1xFlag-ABCD3 and 0.2 mg/mL BSA (to reduce non-specific binding).

FAM-Ceapin A7 or fluorescent ATF6α peptide binding was measured until reaching equilibrium. The plotted equilibrium values represent the mean of 3 replicates with error bars representing standard deviation. Values were plotted as the change in milli-polarization units [ΔmP] by subtracting the measured baseline polarization value of the fluorescent molecule (FAM-Ceapin A7 or fluorescent ATF6α peptide). Data was analyzed and plotted using Prism v.10.5.0 (GraphPad).

#### ATPase assays

Relating to **Figure 3C** and **3H**, ATPase activity was measured using the malachite green ATPase/GTPase Activity Assay Kit (Sigma). 1.5 μM 1xFlag-ABCD3 was pre-incubated with or without 4 μM phytanoyl-CoA (Avanti) and indicated concentrations of Ceapin A7, Ceapin A5, coenzyme A (Sigma #C3019), CoA-hybrid 1, or CoA-hybrid 2 for 30 minutes at 25°C in a 40 μL volume in ATPase Assay Buffer (25 mM HEPES pH 7.5, 100 mM NaCl, 0.02% GDN, 0.5 mM

TCEP). 1 mM ATP/MgCl<sub>2</sub> was added for 30 minutes at 37°C and the reaction was stopped by adding quenching reagent from the assay kit and incubated for 30 minutes at 25°C. Free phosphate was measured with absorbance at 620 nm using the CLARIOstar Plus plate reader (BMG LabTech). The plotted values represent the mean of 3 replicates with error bars representing standard deviation. Data was analyzed and plotted using Prism v.10.5.0 (GraphPad).

#### **Treatment of cells with stressors**

Cells were treated with the indicated pharmacological compounds at the concentrations and durations listed below unless otherwise specified in the figure legends. Cells were treated with Ceapin A7 (10 µM, 4 h); the ER stressors tunicamycin (2.5 µg ml<sup>-1</sup>, 4 h), thapsigargin (500 nM, 4 h), or dithiothreitol (DTT; 1 mM, 1 h); the lipid metabolic stressors palmitic acid (400 µM, 4 h) or oleic acid (200 µM, 4 h); the peroxisomal lipid substrates trihydroxycholestanoic acid (THCA; 10 µM, 4 h), dihydroxycholestanoic acid (DHCA; 10 µM, 4 h), or phytanic acid (10 µM, 4 h); the mitochondrial inhibitors FCCP (1 µM, 4 h) or oligomycin (1 µM, 4 h); or hydrogen peroxide (1 mM, 1 h) to induce oxidative stress. THCA, DHCA, phytanic acid, palmitic acid, and oleic acid were conjugated to 1% fatty acid-free bovine serum albumin (BSA) in DMEM for 30 min before addition to cells. Where indicated, actin polymerization was inhibited with CK-666 (100 µM, 2 h), lysosomal degradation was inhibited with chloroquine (50 µM, 4 h), and cells were incubated with the fluorescent fatty acid analogue BODIPY-C12 (1 µM, 4 h). Vehicle-treated cells served as controls in all experiments.

#### **Enrichment of ubiquitylated proteins**

To assess ABCD3 ubiquitylation, control and ATF6A-depleted U2OS cells were treated with vehicle or Ceapin A7 (10 µM) for 6 h. Cells were washed with ice-cold PBS and lysed in 50 mM Tris-HCl (pH 7.5), 150 mM NaCl, and 1% Triton X-100 supplemented with protease inhibitors and 10 mM N-ethylmaleimide (NEM) to inhibit deubiquitylating enzymes and preserve ubiquitin conjugates. Lysates were clarified by centrifugation, and protein concentrations were determined using a BCA assay. One milligram of total protein from each sample was incubated with TUBE2 agarose (LifeSensors, UM402) for 2 h at 4 °C with end-over-end rotation. Following incubation, the resin was washed extensively with lysis buffer, and bound proteins were eluted by boiling in SDS sample buffer. Eluted proteins together with corresponding input lysates were resolved by SDS-PAGE and analyzed by immunoblotting using antibodies against ABCD3, and Ubiquitin.

#### **Lysosome immunoprecipitation (Lyso-IP)**

Lysosomes were isolated from GFP-ATF6α U2OS cells with stable expression of 3×HA-TMEM192 using lysosome immunoprecipitation<sup>4</sup>. Cells were treated with vehicle, Ceapin A7, THCA for 16 h, washed with ice-cold PBS, and harvested in KPBS (136 mM KCl, 10 mM KH<sub>2</sub>PO<sub>4</sub>, pH 7.5) buffer supplemented with protease inhibitors. Cells were mechanically disrupted using a dounce homogenizer under conditions preserving organelle integrity, and post-nuclear supernatants were obtained by low-speed centrifugation (800g, 5 min). The lysosome-containing supernatant was incubated with anti-HA magnetic beads (Thermo Fischer Scientific, #88836) at 4 °C with gentle rotation for 20min. Beads were rapidly washed with ice-cold isolation buffer, and lysosome-enriched material was eluted for immunoblotting or proteomic analysis. Lysosome enrichment and purity were assessed by immunoblotting for the lysosomal membrane protein LAMP1 and the lysosomal luminal enzyme cathepsin C. Lysosomal fractions were further analyzed for ATF6α, ABCD3, PEX14, and other indicated proteins. Bead-associated material was also subject to mass spectrometric analysis.

##### *Resuspension*

De-salted peptide samples were centrifuged under vacuum to dryness, then resuspended with 3% ACN, 0.1% Trifluoroacetic acid (TFA) in water at a concentration of 100 ng/uL.

##### *LC-MS/MS method*

Liquid chromatography–tandem mass spectrometry (LC–MS/MS) was performed with a Thermo Fisher Vanquish Neo UHPLC coupled to a Thermo Fisher Orbitrap Ascend Tribrid mass spectrometer. Peptides were separated on an IonOpticks Aurora Ultimate column (25 cm × 75 μM, 1.7 μM C18), heated to 50°C inside a Sonation oven. The LC mobile phases consisted of 0.1% formic acid in water (solvent A) and 80% ACN, 0.1% formic acid in water (solvent B), with all solvents being LC-MS grade. A two-step linear gradient was delivered at 400 nl/min for 3–35% B in 60 min, followed by 35–45% B in 6 min. The column was washed by increasing solvent B to 95% in 1 minute, maintaining for 3 minutes at 400 nl/min, then increasing the flowrate to 800 nl/min in 1 minute, and washing for an additional 4 minutes. The Orbitrap Ascend mass spectrometer was operated in the positive mode using a nanospray ionization source (NSI), with a spray voltage of 1,800 V, and ion transfer tube temperature of 275°C. Full MS scans were acquired in the Orbitrap at a resolution of 120,000, normalized automatic gain control (AGC) target at 250%, maximum inject time (IT) of 251 ms, RF-lens setting of 60, and a full scan range of 350 — 1,050 m/z. Data-independent acquisition (DIA) scans were acquired with higher-energy collisional dissociation (31% normalized collision energy), at an Orbitrap resolution of 15,000, over a scan range of 200 - 1,800 m/z. Seventy overlapping 9.5 m/z windows covering the mass range of 389.25 – 1010.25 m/z were used, with an AGC target of 1,000%, with a maximum IT of 27 ms, and RF-lens setting of 60.

##### **HaloTag labeling and peroxisome staining with Peroxi-SPY555**

Halo-HA-ATF6α knock-in cells were seeded onto glass-bottom imaging dishes 24 h before imaging. Prior to image acquisition, cells were incubated with 100nM HaloTag ligand (Janelia Fluor-650, Promega HT1110) for 30min to fluorescently label endogenous Halo-ATF6α. Excess ligand was removed by washing with PBS and adding complete medium before imaging.

Peroxisomes were labeled using the fluorescent probe 1μM Peroxi-SPY555 (CY-SC207) for 15 min. For experiments involving Ceapin A7 or other pharmacological treatments, cells were incubated with compounds for the indicated durations before HaloTag and peroxisome labeling. Live-cell imaging was performed immediately following staining.

##### **Split-Venus fluorescence complementation assay**

To monitor the interaction between ATF6α and ABCD3 in living cells, a bimolecular fluorescence complementation (Split-Venus) assay was employed. Cells expressing ABCD3-Venus (1-211), and Venus (212–239)-ATF6α were treated with 10μM Ceapin A7 for 4 h or different stressors followed by fixation in 4% paraformaldehyde and imaging. Venus fluorescence was acquired using identical microscope settings across all experimental conditions. Because reconstitution of the Venus fluorophore stabilizes transient protein interactions, fluorescence intensity and puncta abundance represent cumulative ATF6α–ABCD3 interaction events rather than native interaction kinetics. The number of Venus-positive puncta per cell was quantified using Fiji/ImageJ following automated thresholding and particle analysis. At least three independent biological replicates taken from different experiments were analyzed for each condition.

#### **Split-Venus ATF6α-ABCD3 interactome**

To identify proteins associated with the ATF6α-ABCD3 complex, U2OS cells with stable expression of Split-Venus tagged ATF6α and ABCD3 were treated with vehicle or Ceapin A7 or other drugs. Cells were washed twice with ice-cold phosphate-buffered saline and lysed in ice-cold non-denaturing lysis buffer containing 50 mM Tris-HCl (pH 7.5), 150 mM NaCl, 1% TritonX-100, and protease and phosphatase inhibitor cocktails. Lysates were clarified by centrifugation at $16,000 \times g$ for 15 min at 4 °C.

1mg protein was incubated with Chromo Tek GFP-Trap agarose beads which efficiently recognize the reconstituted Venus fluorophore, for 2 h at 4 °C with end-over-end rotation. Following immunoprecipitation, beads were washed five times with the wash buffer (50 mM Tris-HCl (pH 7.5), 300 mM NaCl, 0.1% TritonX-100) to remove nonspecific interactors. Bound proteins were either eluted in SDS sample buffer for immunoblot analysis or processed directly on-bead for liquid chromatography-tandem mass spectrometry (LC-MS/MS).

#### **Sample preparation for mass spectrometry**

For proteomic analysis, bead-bound proteins were eluted from GFP beads (for the Split-Venus ATF6α-ABCD3 interactome) and from HA magnetic beads (for the lyso-IP) by incubation with 4 M urea for 30 min at 37 °C. Samples were subsequently reduced using 5 mM dithiothreitol for 30 min at 56 °C and alkylated using 40 mM chloroacetamide at room temperature in the dark for 30 min. Following dilution of the urea concentration to 0.8 M with 50 mM Tris-HCl (pH 8.0), proteins were digested overnight at 37 °C with sequencing-grade modified trypsin and Lys-C. Resulting peptides were acidified with trifluoroacetic acid, desalted using C18 StageTips, and analyzed by nanoLC-MS/MS. Raw spectra were searched against the UniProt human reference proteome using MaxQuant with label-free quantification enabled. Proteins identified as contaminants, reverse hits, or identified only by site were excluded from downstream analysis. Differential protein enrichment between vehicle- and Ceapin A7-treated samples was determined using label-free intensity values. Functional enrichment analysis of significantly enriched proteins was performed using Gene Ontology (GO) pathway analysis.

#### *Resuspension*

De-salted peptide samples were centrifuged under vacuum to dryness, then resuspended with 3% ACN, 0.1% Trifluoroacetic acid (TFA) in water at a concentration of 10 ng/uL.

#### *LC-MS/MS method*

Liquid chromatography–tandem mass spectrometry (LC-MS/MS) was performed with a Thermo Fisher Vanquish Neo UHPLC coupled to a Bruker timsTOF SCP mass spectrometer via CaptiveSpray nanoelectric spray ion source (Bruker) operated in positive mode. Peptides were separated on an Ionopticks Aurora Rapid CSI column (5 cm x 75 μm 1.7 μm C18), heated to 50°C inside a Bruker Column Toaster. The LC mobile phases consisted of 0.1% formic acid in water (solvent A) and 80% ACN, 0.1% formic acid in water (solvent B), with all solvents being LC-MS grade. A two-step linear gradient was delivered at 200 nl/min for 0–35% B in 25 min, followed by

35–45% B in 3 min. The column was washed by increasing solvent B to 95% in 1 minute and holding it for 1 minute.

The timsTOF SCP mass spectrometer was operated in dia-PASEF mode. MS1 scans were acquired from 100 - 1700 m/z from 1.3 - 0.7 1/K0[V-s/cm-2]. Seven diaPASEF scans covering from 300-1200 m/z and 1.3 - 0.7 1/K0[V-s/cm-2] consisting of 3 or 4 windows ranging from 31 to 61 m/z in width (complete table below). Accumulation and ramp times were fixed to 75 ms.

| Id | Type | Cycle | OneOverK0Start | OneOverK0End | IsolationMz | IsolationWidth | CollisionEnergy |
| --- | --- | --- | --- | --- | --- | --- | --- |
| 1 | 0 | 0 |  |  |  |  |  |
| 2 | 1 | 1 | 1.215 | 1.3 | 1170 | 61 |  |
| 3 | 1 | 1 | 1.055 | 1.215 | 912.5 | 36 |  |
| 4 | 1 | 1 | 0.865 | 1.055 | 667.5 | 36 |  |
| 5 | 1 | 1 | 0.7 | 0.865 | 422.5 | 36 |  |
| 6 | 1 | 2 | 1.189 | 1.3 | 1120 | 41 |  |
| 7 | 1 | 2 | 1.029 | 1.189 | 877.5 | 36 |  |
| 8 | 1 | 2 | 0.836 | 1.029 | 632.5 | 36 |  |
| 9 | 1 | 2 | 0.7 | 0.836 | 387.5 | 36 |  |
| 10 | 1 | 3 | 1.17 | 1.3 | 1085 | 31 |  |
| 11 | 1 | 3 | 1.005 | 1.17 | 842.5 | 36 |  |
| 12 | 1 | 3 | 0.808 | 1.005 | 597.5 | 36 |  |
| 13 | 1 | 3 | 0.7 | 0.808 | 352.5 | 36 |  |
| 14 | 1 | 4 | 1.148 | 1.3 | 1055 | 31 |  |
| 15 | 1 | 4 | 0.964 | 1.148 | 807.5 | 36 |  |
| 16 | 1 | 4 | 0.782 | 0.964 | 562.5 | 36 |  |
| 17 | 1 | 4 | 0.7 | 0.782 | 317.5 | 36 |  |
| 18 | 1 | 5 | 1.126 | 1.3 | 1020 | 41 |  |

|  |  |  |  |  |  |  |
| --- | --- | --- | --- | --- | --- | --- |
| 19 | 1 | 5 | 0.938 | 1.126 | 772.5 | 36 |
| 20 | 1 | 5 | 0.7 | 0.938 | 527.5 | 36 |
| 21 | 1 | 6 | 1.101 | 1.3 | 982.5 | 36 |
| 22 | 1 | 6 | 0.913 | 1.101 | 737.5 | 36 |
| 23 | 1 | 6 | 0.7 | 0.913 | 492.5 | 36 |
| 24 | 1 | 7 | 1.078 | 1.3 | 947.5 | 36 |
| 25 | 1 | 7 | 0.889 | 1.078 | 702.5 | 36 |
| 26 | 1 | 7 | 0.7 | 0.889 | 457.5 | 36 |

#### Fluorescence imaging and image quantification

For microscopy, cells grown on Cellvis 24-well glass-bottom plates featuring high-performance #1.5 cover glass bottoms. Cells were fixed with 4% paraformaldehyde for 10min , permeabilized using blocking buffer (0.1% saponin in 10% FBS in PBS), and incubated with primary antibodies overnight at 4 °C. Following washing, cells were incubated with species-appropriate fluorescent secondary antibodies and imaged in PBS.

Fluorescence imaging was performed using a Nikon Ti2 inverted microscope equipped with a Yokogawa CSU-W1 spinning-disk confocal scanner, controlled using Micro-Manager (v2.0), and maintained at 37 °C and 5% CO<sub>2</sub> using a stage-top environmental chamber. Images were acquired using either a Plan Apo 100×/1.45 NA oil-immersion objective for high-resolution visualization of ER-peroxisome contact sites and autophagy-associated structures or a Plan Apo 40×/1.4 NA objective for quantitative analysis of GFP-ATF6α or Split-Venus puncta. All images were acquired using identical laser power, exposure time, camera gain, and acquisition settings for all experimental conditions within each experiment.. Z-stacks with 0.3μm z-intervals were acquired where indicated. Images shown are single slices at the center of the cell (where the nucleus is at best focus) unless specified otherwise.

Colocalization between Halo-ATF6α puncta and peroxisomes labeled with either Peroxi-SPY555 or PEX14 immunostaining was quantified using the Manders overlap coefficient (M1) following background subtraction with the Coloc2 plugin. At least three independent biological replicates were analyzed, and multiple fields containing randomly selected cells were quantified for each condition.

Image analysis was performed using FIJI/ImageJ (v2.x). Following background subtraction, GFP-ATF6α or Split-Venus puncta were segmented by applying a fixed intensity threshold and quantified using the Analyze Particles function (Analyze → Analyze Particles). Objects with an area between 0.1 and 2.0 $\mu\text{m}^2$ and a circularity of 0.2–1.0 were included for analysis. The total number of Venus-positive puncta was recorded for each field of view. To normalize puncta abundance, the display threshold of the fluorescence image was adjusted to visualize diffuse cellular fluorescence and delineate individual cell boundaries. The number of cells within each field was then determined manually, and puncta counts were normalized to the total number of cells and expressed as Venus puncta per cell. Cells located at the edge of the field of view or exhibiting abnormal morphology were excluded from analysis. Identical image processing, thresholding, and particle analysis parameters were applied to all images within an experiment. Quantification was performed from at least three biological replicates taken from 3 independent experiments.

For quantification of peroxisome abundance, cells were fixed and immunostained for PEX14 as described above. Images were acquired using identical acquisition settings for all experimental conditions. Image analysis was performed using FIJI/ImageJ (v2.x) following uniform background subtraction. The PEX14 channel was segmented using a fixed intensity threshold that was applied uniformly across all samples within each experiment. Individual peroxisomes were identified using the Analyze Particles function with an area filter of 0.05–4.0 $\mu\text{m}^2$ and a circularity threshold of 0.20–1.00. The total number of PEX14-positive puncta was quantified for each field of view and normalized to the number of cells in the corresponding field, yielding peroxisomes per cell. Quantification was performed on at least 50 cells from three independent biological replicates for each condition.

#### **Peroxisomal HyPer3 imaging**

Peroxisomal hydrogen peroxide levels were monitored using U2OS cells stably expressing the genetically encoded ratiometric fluorescent biosensor HyPer3-SKL, in which HyPer3 is targeted to the peroxisomal lumen via a C-terminal SKL targeting sequence. Cells were treated with vehicle, Ceapin A7, THCA, or the indicated compounds and imaged live using identical acquisition settings for all experimental conditions. HyPer3 was sequentially excited at 420 nm and 500 nm, and fluorescence emission was collected using identical laser power, exposure time, and detector settings. Images were acquired sequentially to minimize photobleaching and channel misalignment.

Ratiometric image analysis was performed using FIJI/ImageJ (v2.x). Images acquired at both excitation wavelengths were first subjected to identical background subtraction. Pixel-wise ratio images ($F_{500}/F_{420}$) were generated using the Image Calculator function (Process → Image Calculator; Divide) and displayed using the Rainbow RGB lookup table (LUT) for visualization. Individual peroxisomes were identified as discrete fluorescent puncta, and the fluorescence intensity of each punctum was measured in both excitation channels. The HyPer3 red/oxidized ratio ($F_{500}/F_{420}$) was calculated for each peroxisome and used as a quantitative measure of peroxisomal hydrogen peroxide levels. At least 30 cells from three independent biological replicates were analyzed for each condition.

#### **Immunofluorescence analysis of p62 recruitment to peroxisomes**

Halo-ATF6α U2OS cells were seeded onto glass coverslips and treated with vehicle, Ceapin A7, THCA, or the indicated compounds for 16 h. Unless otherwise specified, Ceapin A7 and THCA were used at 10 $\mu\text{M}$. Cells were washed with PBS, fixed with 4% paraformaldehyde, and permeabilized and blocked for 1 h at room temperature in PBS containing 0.1% saponin and 10% fetal bovine serum. Cells were incubated overnight at 4 °C with primary antibodies against SQSTM1/p62 and PEX14 diluted in the blocking buffer. Samples were washed with PBS containing 10% fetal bovine serum and incubated for 1 h at room temperature with species-appropriate Alexa Fluor-conjugated secondary antibodies. Coverslips were washed and mounted in an antifade mounting medium before imaging.

Images were acquired using identical acquisition settings for all conditions within each experiment. Image analysis was performed in FIJI/ImageJ following uniform background subtraction. The PEX14 and p62 channels were independently thresholded using fixed parameters and converted to binary masks. Colocalized structures were identified by generating the intersection of the PEX14 and p62 masks using the Image Calculator AND operation. The resulting overlap objects were quantified using the Analyze Particles function with identical size and circularity criteria across all samples. A pexophagy-associated event was defined as one discrete overlap object containing both PEX14 and p62 signal. Events were normalized to the number of cells in each field. At least 30 cells from three independent biological replicates were analyzed for each condition.

#### **Tandem fluorescent pexophagy reporter assay**

Pexophagy-associated lysosomal delivery was monitored using a tandem fluorescent reporter consisting of GFP and mCherry fused to the C-terminal peroxisomal targeting sequence SKL. U2OS cells with stable expression of GFP–mCherry–SKL were treated with vehicle, Ceapin A7, THCA for 16 h. In intact peroxisomes and other neutral compartments, both fluorophores remain fluorescent and appear as GFP-positive/mCherry-positive puncta. Following delivery to acidic lysosomal compartments, GFP fluorescence is quenched whereas mCherry fluorescence persists, generating mCherry-only puncta. Images were acquired using identical laser power, exposure time, camera gain, and acquisition settings for all conditions within each experiment. Image analysis was performed in FIJI/ImageJ following uniform background subtraction. The GFP and mCherry channels were independently thresholded using fixed parameters and converted to binary masks. GFP-positive/mCherry-positive puncta were identified from the intersection of the two masks using the Image Calculator AND operation. To identify mCherry-only structures, the GFP mask was subtracted from the mCherry mask using the Image Calculator Subtract operation. The resulting mCherry-only objects were quantified using the Analyze Particles function with identical size and circularity criteria across all samples. A pexophagy event was defined as one discrete mCherry-positive/GFP-negative punctum. The number of pexophagy events was normalized to the number of cells in each field and expressed as mCherry-only puncta per cell. At least 30 cells from three independent biological replicates were analyzed for each condition.

#### **Statistical analysis of fluorescence imaging**

All imaging experiments were performed using a minimum of three biological replicates taken from 3 independent experiments. Quantitative data are presented as mean ± s.e.m. for bar graphs. Box Plots were generated in Box Plot R (http://shiny.chemgrid.org/boxplotr/). For box plots, Center lines indicate medians; box limits represent the 25th and 75th percentiles; whiskers extend 1.5× the interquartile range; outliers are shown as individual points. Statistical analyses were performed using Microsoft Excel. Comparisons between two groups were analyzed using two-tailed unpaired Student's t-tests. Sample sizes, statistical tests, and P values are provided in the corresponding figure legends. Differences were considered statistically significant at $P < 0.05$.

#### **NanoBRET assay**

8 x 10<sup>5</sup> 293T cells (ATCC CRL-3216) were seeded per well in 6-well plates (Costar, Cat. 3516) and incubated at 37°C with 5% CO<sub>2</sub> for 6 hours. ATF6α and ABCD3 proteins were tagged with Halo or NanoLuciferase (NLuc) tags and transfected as Halo:NLuc pairs at a ratio of 1:10. Specifically, a mixture containing 1 µg NLuc plasmid, 10 µg Halo plasmid, 15 µl FuGENE HD Transfection Reagent (Promega, Cat. E2311), and 100 µl Opti-MEM I reduced serum medium without phenol red (Gibco, Cat. 31985062) per well was incubated at room temperature for 20 minutes before being added dropwise to the cells in the 10 cm dishes. After 24 hours, cells from the 10 cm dishes were trypsinized, washed once with medium (5 min, 600 g, room temperature), then resuspended in Opti-MEM I reduced serum medium, without phenol red and reseeded into white solid-bottom, TC-treated 96-well plates (Costar, Cat. 3917), with or without the addition of HaloTag NanoBRET 618 Ligand (Promega, Cat. G9801). After 18 hours of incubation, a 12-point compound titration of Ceapin A7 (dilution in Opti-MEM I, 0 to 10 µM) was added to the cells. Following 1 hour of treatment, NanoBRET Nano-Glo Substrate (Promega, Cat. N1571) was added. The 96-well plates were imaged using the GloMax Discover Microplate Reader (Promega, Cat. GM3000) with the default “BRET: NanoBRET 618” protocol (Donor: 450 BP; Acceptor: 600 LP; Integration: 0.3 sec). Data analysis was performed according to the user manual.

#### **HDX experiment**

##### *General procedure*

Samples containing ABCD3 and ATF6α at a 1:1 molar ratio, with or without ligand as indicated, were diluted 1:20 into D<sub>2</sub>O labeling buffer (pD 7.1, 10 mM HEPES, 150 mM NaCl) to initiate hydrogen–deuterium exchange; triplicate measurements were collected for each condition measured at labeling times (seconds): 60.0, 370.0, 2276.0, 14200.0, 86400.0. Labeling was quenched at the indicated times by 1:1 dilution with a pH 2.50 solution containing 1 M glycine and 2.7 M guanidine hydrochloride.

Labeling, quenching, and online sample preparation were performed using a Trajan HDX system equipped with an extended-rail LEAP PAL HDX autosampler (Trajan Scientific). Quenched samples (95 µL) were digested at 150 µL/min using an immobilized Nepenthesin II protease column (AffiPro) and trapped on a BEH Vanguard C18 column (Waters). Peptides were washed isocratically for 3 min and separated on a 1 × 50 mm BEH C18 analytical column (Waters) using a binary gradient of water (A) and acetonitrile (B), each containing 0.1% formic acid and 0.04% TFA, as previously recommended<sup>5,6</sup>.

Eluate was introduced by electrospray at 35 µL/min into an Orbitrap Exploris 480 mass spectrometer (Thermo Fisher Scientific) operated in MS-only mode with the following settings: ion-transfer-tube temperature, 120 °C; HESI temperature, 75 °C; spray voltage, 4.0 kV; and RF lens, 50%. Automatic gain control and injection times were determined automatically at a resolving power of 120,000. Data-dependent acquisition for peptide identification was performed using top-5 DDA at resolving powers of 60,000 for MS1 and 7,500 for MS2.

Extracted ion chromatograms (XICs) for each peptide and time point were generated by the Mass-Spectrometry Engine (MSE) v1.0. Tandem MS identifications guiding MSE analysis were generated with Protein Metrics Byonic and Byologic using a 5 ppm tolerance and a nonspecific C-

terminal protease setting. Peptides with an FDR2D score below 0.01 were excluded. Default MSE XICs were summed across charge states before centroid calculation. EPFs were computed as described. To bound each peptide's data, an endpoint immediately before the first measured time point was assigned the minimum D-uptake across all conditions, and one immediately after the last measured time point was assigned the maximum D-uptake. This produces an EPF less than or equal to the true peptide-averaged PF and endpoints permit EPF calculation when traces do not overlap. Only peptides detected in every condition were included. Peptide-averaged EPFs from all peptides overlapping each residue were used to calculate WEPF. For fully exchanged or unexchanged peptides, EPF was set to 1 and uncertainty to 0.

Residue-level WEPFs were calculated from all peptides reporting on a residue in $\log_{10}(\text{PF})$ space. The subscript $i$ denotes a peptide. At each time point, the maximum, median, and minimum triplicate values defined three uptake traces per condition per peptide. Computing the EPF between two conditions thus yielded an indexed peptide-level EPF comparison ensemble. Treating it as an ensemble rather than a set preserves duplicate values in the mean. Here $k$ indexes the comparisons and $K_i$ is their number:

$$\mathcal{E}_i = (\text{EPF}_{i,k})_{k=1}^{K_i}$$

$$\mu_i = \log_{10}(\text{mean}(\mathcal{E}_i)), L_i = \log_{10}(\min(\mathcal{E}_i)), H_i = \log_{10}(\max(\mathcal{E}_i))$$

A deliberately conservative proxy for uncertainty was defined as the largest deviation:

$$\sigma_i = \max\{\mu_i - L_i, H_i - \mu_i, 0\}$$

For residue weighting, peptide $i$ was assigned $w_i = 1/\ell_i$, where the number of exchange sites was $\ell_i$.

Let $I_r$ denote the set of indices for peptides that report on residue $r$. The normalized peptide weight and the residue-level weighted mean are defined as follows:

$$\alpha_i = \frac{w_i}{\sum_{j \in I_r} w_j}$$

$$\mu_r = \sum_{i \in I_r} \alpha_i \mu_i$$

Uncertainty was propagated using within- and between-peptide variance terms:

$$V_{\text{within},r} = \sum_{i \in I_r} \alpha_i^2 \sigma_i^2, \quad V_{\text{between},r} = \sum_{i \in I_r} \alpha_i (\mu_i - \mu_r)^2$$

The total variance and residue-level standard deviation were:

$$V_{\text{total},r} = V_{\text{within},r} + V_{\text{between},r}, \quad \sigma_r = \sqrt{V_{\text{total},r}}$$

Thus, the final quantity is $\text{WEPF}_r = 10^{\mu_r}$, with uncertainty interval $10^{\mu_r \pm \sigma_r}$. If $|\mu_r| - \sigma_r < 0$, the result is considered insignificant and reported as $\mu_r = 0$ (*i. e.*, $\text{WEPF}_r = 1$). The value was also set to unity when the shortest peptide covering a residue was fully exchanged or showed no exchange (flat uptake traces). The associated Excel file provides replicate-level uptake, peptide EPF ensembles, limit-state labels, and WEPF calculations for all peptides and conditions.

Raw mzXML files, peptide ID's and retention times, the ABCD3 protein sequence used for HX, and MSE output files (.deut or .allH) are deposited in the MassIVE database under accession code MSV000102763;. The "Coverage" tab of the supplemental HX Excel file (260807\_HX\_SI\_ATFABCDmanuscript) reports recommended experimental metrics<sup>7</sup>, including coverage, redundancy, and a coverage map. This file also contains peptide uptake values, EPF-calculation components, and imputed endpoints. The SI\_HX PDF (260801\_HXplots\_SI\_R3) lists every peptide, condition-specific uptake, and a theoretical unprotected exchange trace. Because the HX sequence contains additional residues, manuscript numbering uses a -34 offset to match canonical residue numbers; HX numbering may remain in the supplemental files. For each residue, this file lists the contributing peptides, shortest-peptide substitutions, and assigned limits. Limits that disable EPF calculations and set the value to 1 are labeled "fully exchanged" or "no exchange" at the first and last time points, respectively. Other limits reflect trace divergence at the earliest or latest time points.

##### Software versions used:

MSE v1.0 (see GitHub repository).

Protein Metrics Byonic Version 4.5.2

Protein Metrics Byologic Version 4.5-53 x64

Chronos Version 5.1.10

MSConvertGUI Version 3.0.22027-235693e

#### Synthesis of Ceapin analogs

##### CoA-A7-1

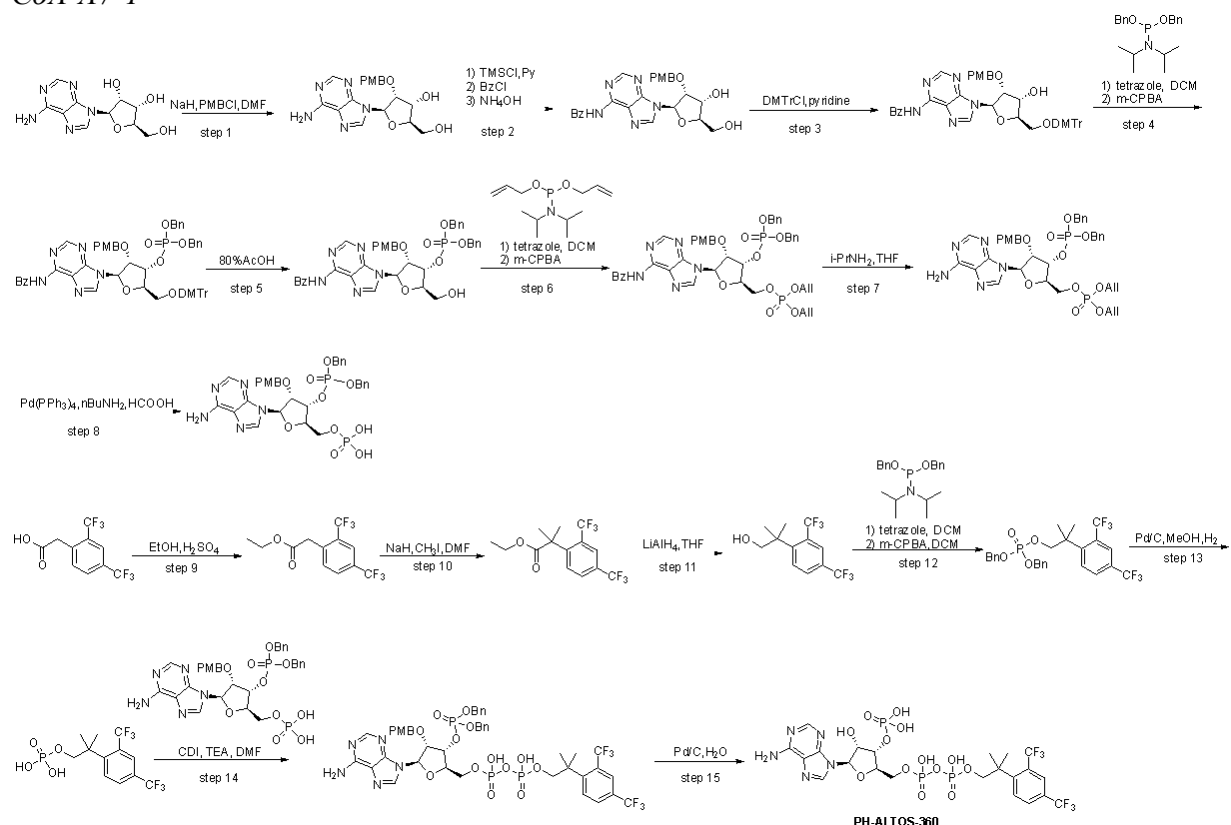

See Supplementary Methods below for detailed protocol.

### CoA-A7-2

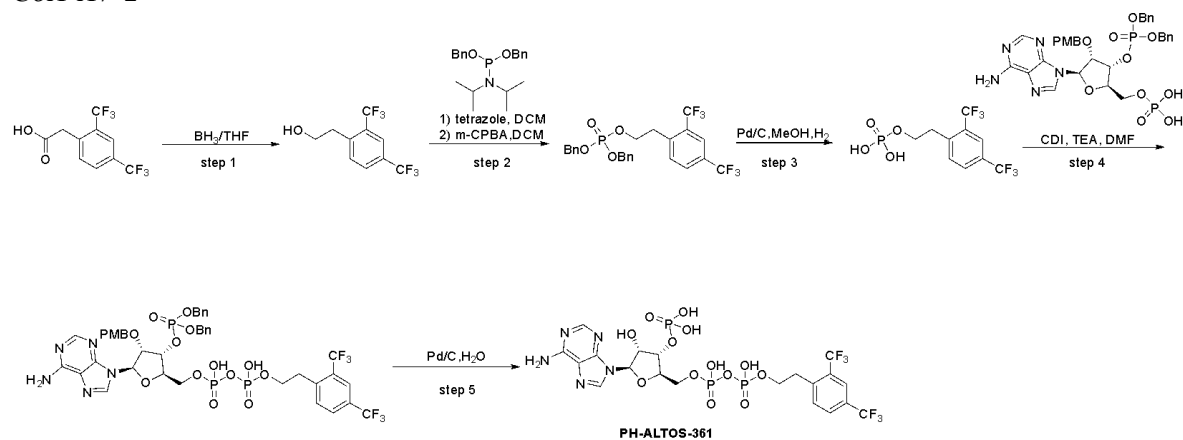

See **Supplementary Methods** below for detailed protocol.

### Sequence alignments

Relating to **Extended Data Figure 4C**, ATF6α homologous sequences were aligned in ClustalOmega and visualized in JalView v.2.11.5.1. Uniprot accession codes for sequences in the alignment: Q20435 (*C. elegans*), G3V909 (*R. norvegicus*), F6VAN0 (*M. musculus*), P18850 (*H. sapiens*), Q7K2V1 (*D. melanogaster*), H2Q0H3 (*P. troglodytes*), H0VGM7 (*C. porcellus*).

### Quantification and statistical analysis for *in vitro* assays

All data were tested for statistical significance with Prism 10.5.0 (GraphPad). No randomization or blinding was performed. For all statistical analyses, variances across groups were not significantly different (as evaluated by Brown-Forsythe test or F test) and therefore no corrections were used. Relating to **Figure 3C**, the one-way ANOVA with *post-hoc* Bartlett's test *P* values are as follows: ABCD3 vs. ABCD3 + 2 μM Ceapin A7 = 0.0323, ABCD3 vs. ABCD3 + 10 μM Ceapin A7 = 0.0002, ABCD3 vs. ABCD3 + 50 μM Ceapin A7 < 0.0001, ABCD3 + 4 μM phytanoyl-CoA vs. ABCD3 + 4 μM phytanoyl-CoA + 25 μM Ceapin A7 < 0.0001, ABCD3 + 4 μM phytanoyl-CoA vs. ABCD3 + 4 μM phytanoyl-CoA + 25 μM Ceapin A5 = 0.5703. Relating to **Figure 3G**, the one-way ANOVA with *post-hoc* Dunnett's test *P* values are as follows: ABCD3 vs. ABCD3 + 25 μM Ceapin A7 = 0.6753, ABCD3 vs. ABCD3 + 25 μM phytanoyl-CoA < 0.0001, ABCD3 vs. ABCD3 + 25 μM CoA < 0.0001, ABCD3 vs. ABCD3 + 25 μM CoA-A7-1 < 0.0001, ABCD3 vs. ABCD3 + 25 μM CoA-A7-2 < 0.0001. Relating to **Figure 3H**, the one-way ANOVA with *post-hoc* Sidak's test *P* values are as follows: ABCD3 vs. ABCD3 + 25 μM CoA < 0.0001, ABCD3 + 25 μM CoA vs. ABCD3 + CoA-A7-1 < 0.0001, ABCD3 + 25 μM CoA vs. ABCD3 + CoA-A7-2 < 0.0001. Relating to **Extended Data Figure 5I**, the unpaired two-tailed *t*-test *P* values are as follows: ABCD3 vs ABCD3 + 5 mM ATP/Vanadate = 0.0024.

### Supplementary Methods

#### Synthesis of CoA-A7-1

**[(2R,3S,4R,5R)-5-(6-aminopurin-9-yl)-2-{{{[2-[2,4-bis(trifluoromethyl)phenyl]-2-methylpropoxy(hydroxy)phosphoryl]oxy(hydroxy)phosphoryl}oxy]methyl}-4-hydroxyoxolan-3-yl]oxyphosphonic acid (PH-ALTOS-360-0)**

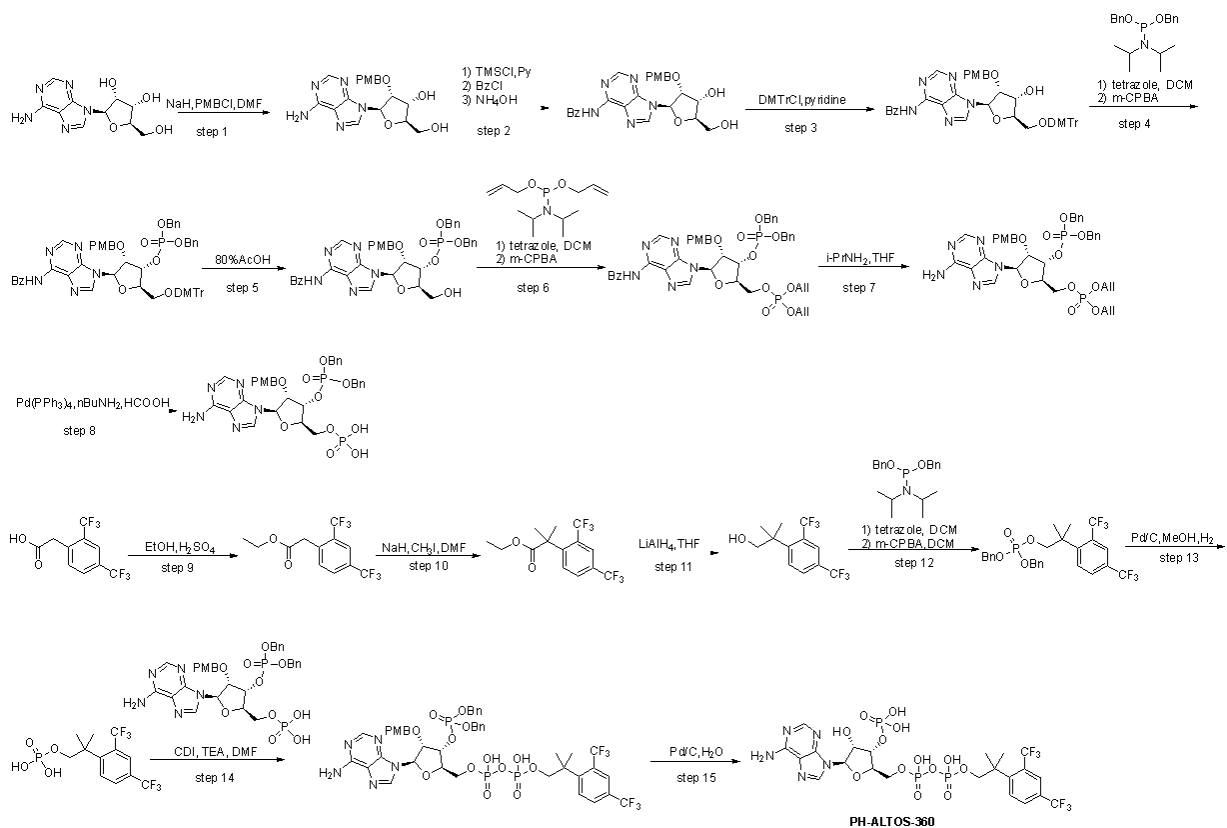

#### Step 1. (2R,3R,4R,5R)-5-(6-aminopurin-9-yl)-2-(hydroxymethyl)-4-[(4-methoxyphenyl)methoxy]oxolan-3-ol

To a stirred solution of adenosine (75.0 g, 281 mmol, 1.00 equiv) in DMF (2.5 L) was added NaH (60%) (13.5 g, 337 mmol, 1.20 equiv) in portions at -5 °C. The reaction was quenched with water (50 mL) at 0 °C. The mixture was concentrated and purified by silica gel column (eluting with DCM / MeOH (50:1)) and re-purified by reverse flash chromatography (column: C18, 330 g, 20-35 μm; mobile phase, A: water (10 mM $\text{NH}_4\text{HCO}_3$) and B: ACN (10% to 30% in 20 min); detector: UV 254/220 nm). The collected fraction was concentrated to afford (2R,3R,4R,5R)-5-(6-aminopurin-9-yl)-2-(hydroxymethyl)-4-[(4-methoxyphenyl)methoxy]oxolan-3-ol (25.0 g, 23% yield) as a white solid. LCMS (ES, $m/z$): 387.85 $[\text{M}+\text{H}]^+$.

#### Step 2. N-{9-[(2R,3R,4R,5R)-4-hydroxy-5-(hydroxymethyl)-3-[(4-methoxyphenyl)methoxy]oxolan-2-yl]purin-6-yl}benzamide

To a stirred solution of (2R,3R,4R,5R)-5-(6-aminopurin-9-yl)-2-(hydroxymethyl)-4-[(4-methoxyphenyl)methoxy]oxolan-3-ol (25.0 g, 64.5 mmol, 1.00 equiv) in pyridine (350 mL) was added chlorotrimethylsilane (35.1 g, 323 mmol, 5.00 equiv) dropwise at 0 °C. The reaction was stirred for 2 h at 25 °C. The reaction was re-cooled to 0 °C. Benzoyl chloride (18.1 g, 129 mmol, 2.00 equiv) was added and the reaction was stirred for 16 h at 25 °C. Next, water (100 mL) and $\text{NH}_3\cdot\text{H}_2\text{O}$ (150 mL) were carefully added at 0 °C. The reaction was stirred for 0.5 h at 25 °C. The reaction was diluted with brine (300 mL) and extracted with DCM (2 x 300 mL). The combined organic phase was dried with anhydrous $\text{Na}_2\text{SO}_4$, filtered and concentrated. The residue was triturated with a mixture of EA (200 mL) and MeOH (4 mL) to afford N-{9-[(2R,3R,4R,5R)-4-hydroxy-5-(hydroxymethyl)-3-[(4-methoxyphenyl)methoxy]oxolan-2-yl]purin-6-yl}benzamide (24.0 g, 77% purity) as a white solid. LCMS (ES, $m/z$): 492.20 $[\text{M}+\text{H}]^+$.

**Step 3. N-{9-[(2R,3R,4R,5R)-5-{[bis(4-methoxyphenyl)(phenyl)methoxy]methyl}-4-hydroxy-3-[(4-methoxyphenyl)methoxy]oxolan-2-yl]purin-6-yl}benzamide**

To a stirred solution of N-{9-[(2R,3R,4R,5R)-4-hydroxy-5-(hydroxymethyl)-3-[(4-methoxyphenyl)methoxy]oxolan-2-yl]purin-6-yl}benzamide (24.0 g, 48.8 mmol, 1.00 equiv) in pyridine (350 mL) was added 1-[chloro(4-methoxyphenyl)benzyl]-4-methoxybenzene (24.8 g, 73.2 mmol, 1.50 equiv) in portions at 0 °C. The reaction was stirred for 16 h at 25 °C. The reaction was quenched with NaHCO<sub>3</sub> (sat., aq.) (300 mL). The resulting mixture was extracted with EtOAc (3 x 300 mL). The combined organic layers were washed with brine (500 mL), dried over anhydrous Na<sub>2</sub>SO<sub>4</sub>. After filtration, the filtrate was concentrated under reduced pressure. The residue was purified by silica gel column, eluting with EA (100%) to afford N-{9-[(2R,3R,4R,5R)-5-{[bis(4-methoxyphenyl)(phenyl)methoxy]methyl}-4-hydroxy-3-[(4-methoxyphenyl)methoxy]oxolan-2-yl]purin-6-yl}benzamide (27.0 g, 69% yield) as a white solid. LCMS (ES, m/z): 794.40 [M+H]<sup>+</sup>.

**Step 4. (2R,3R,4R,5R)-5-(6-benzamidopurin-9-yl)-2-{[bis(4-methoxyphenyl)(phenyl)methoxy]methyl}-4-[(4-methoxyphenyl)methoxy]oxolan-3-yl dibenzyl phosphate**

To a stirred solution of N-{9-[(2R,3R,4R,5R)-5-{[bis(4-methoxyphenyl)(phenyl)methoxy]methyl}-4-hydroxy-3-[(4-methoxyphenyl)methoxy]oxolan-2-yl]purin-6-yl}benzamide (27.0 g, 34.0 mmol, 1.00 equiv) and tetrazole (0.45 M in acetonitrile) (296 mL, 136 mmol, 4.00 equiv) in DCM (200 mL) was added [bis(benzyloxy)phosphanyl]diisopropylamine (47.0 g, 136 mmol, 4.00 equiv) dropwise at 0 °C under nitrogen atmosphere. The reaction was stirred for 1 h at 0 °C. Then m-CPBA (23.5 g, 136 mmol, 4.00 equiv) was added. The resulting solution was stirred for 2 h at 0 °C. The reaction was quenched with NaHCO<sub>3</sub> (sat., aq.) (200 mL). The resulting mixture was extracted with DCM (3 x 200 mL). The combined organic layers were washed with brine (200 mL), dried over anhydrous Na<sub>2</sub>SO<sub>4</sub>. After filtration, the filtrate was concentrated under reduced pressure. The residue was purified by silica gel column, eluting with EA (100%) to afford (2R,3R,4R,5R)-5-(6-benzamidopurin-9-yl)-2-{[bis(4-methoxyphenyl)(phenyl)methoxy]methyl}-4-[(4-methoxyphenyl)methoxy]oxolan-3-yl dibenzyl phosphate (27.0 g, 75% yield) as a light yellow solid. LCMS (ES, m/z): 1054.55 [M+H]<sup>+</sup>.

**Step 5. (2R,3R,4R,5R)-5-(6-benzamidopurin-9-yl)-2-(hydroxymethyl)-4-[(4-methoxyphenyl)methoxy]oxolan-3-yl dibenzyl phosphate**

A solution of (2R,3R,4R,5R)-5-(6-benzamidopurin-9-yl)-2-{[bis(4-methoxyphenyl)(phenyl)methoxy]methyl}-4-[(4-methoxyphenyl)methoxy]oxolan-3-yl dibenzyl phosphate (27.0 g, 25.6 mmol, 1.00 equiv) in 80% AcOH (1 L) was stirred for 2 h at 25 °C. The reaction was quenched with ice water (800 mL). The resulting mixture was extracted with DCM (2 x 2 L). The combined organic layers were washed with brine (1 L), dried over anhydrous Na<sub>2</sub>SO<sub>4</sub>. After filtration, the filtrate was concentrated under reduced pressure. The residue was purified by silica gel column, eluting with DCM/MeOH (10:1) to afford (2R,3R,4R,5R)-5-(6-benzamidopurin-9-yl)-2-(hydroxymethyl)-4-[(4-methoxyphenyl)methoxy]oxolan-3-yl dibenzyl phosphate (14.0 g, 72.7% yield) as a light yellow oil. LCMS (ES, m/z): 752.30 [M+H]<sup>+</sup>.

**Step 6. diallyl (((2R,3R,4R,5R)-5-(6-benzamido-9H-purin-9-yl)-3-((bis(benzyloxy)phosphoryl)oxy)-4-((4-methoxybenzyl)oxy)tetrahydrofuran-2-yl)methyl)phosphate**

To a stirred solution of (2R,3R,4R,5R)-5-(6-benzamidopurin-9-yl)-2-(hydroxymethyl)-4-[(4-methoxyphenyl)methoxy]oxolan-3-yl dibenzyl phosphate (14.0 g, 18.6 mmol, 1.00 equiv) and tetrazole (0.45 M in acetonitrile) (166 mL, 74.6 mmol, 4.00 equiv) in DCM (100 mL) was added [bis(prop-2-en-1-yloxy)phosphanyl]diisopropylamine (18.3 g, 74.6 mmol, 4.00 equiv) dropwise at 0 °C under nitrogen atmosphere. The reaction was stirred for 2 h at 0 °C. Then m-CPBA (12.8 g, 74.6 mmol, 4.00 equiv) was added. The resulting solution was stirred for 2 h at 0 °C. The reaction was quenched with NaHCO<sub>3</sub> (sat., aq.) (100 mL). The resulting mixture was extracted with DCM (2 x 200 mL). The combined organic layers were washed with brine (200 mL), dried over anhydrous Na<sub>2</sub>SO<sub>4</sub>. After filtration, the filtrate was concentrated under reduced pressure. The residue was purified by silica gel column, eluting with DCM/MeOH (10:1) to afford diallyl (((2R,3R,4R,5R)-5-(6-benzamido-9H-purin-9-yl)-3-((bis(benzyloxy)phosphoryl)oxy)-4-((4-methoxybenzyl)oxy)tetrahydrofuran-2-yl)methyl) phosphate (15.0 g, 88% yield) as a light yellow oil. LCMS (ES, m/z): 912.35 [M+H]<sup>+</sup>.

**Step 7. diallyl (((2R,3R,4R,5R)-5-(6-amino-9H-purin-9-yl)-3-((bis(benzyloxy)phosphoryl)oxy)-4-((4-methoxybenzyl)oxy)tetrahydrofuran-2-yl)methyl)phosphate**

A mixture of diallyl (((2R,3R,4R,5R)-5-(6-benzamido-9H-purin-9-yl)-3-((bis(benzyloxy)phosphoryl)oxy)-4-((4-methoxybenzyl)oxy)tetrahydrofuran-2-yl)methyl) phosphate (15.0 g, 16.5 mmol, 1.00 equiv) and propan-1-amine (9.72 g, 165 mmol, 10.0 equiv) in THF (100 mL) was stirred for 3 h at 25 °C under nitrogen atmosphere. The mixture was concentrated under reduced pressure and purified by silica gel column, eluting with DCM/MeOH (20:1) to afford diallyl (((2R,3R,4R,5R)-5-(6-amino-9H-purin-9-yl)-3-((bis(benzyloxy)phosphoryl)oxy)-4-((4-methoxybenzyl)oxy)tetrahydrofuran-2-yl)methyl) phosphate (7.00 g, 53% yield) as a colorless oil. LCMS (ES, m/z): 808.25 [M+H]<sup>+</sup>.

**Step 8. [(2R,3R,4R,5R)-5-(6-aminopurin-9-yl)-3-[(dibenzyloxyphosphoryl)oxy]-4-[(4-methoxyphenyl)methoxy]oxolan-2-yl]methoxyphosphonic acid**

To a stirred mixture of diallyl (((2R,3R,4R,5R)-5-(6-amino-9H-purin-9-yl)-3-((bis(benzyloxy)phosphoryl)oxy)-4-((4-methoxybenzyl)oxy)tetrahydrofuran-2-yl)methyl) phosphate (4.00 g, 4.95 mmol, 1.00 equiv), butan-1-amine (2.17 g, 29.7 mmol, 6.00 equiv) and PPh<sub>3</sub> (780 mg, 2.97 mmol, 0.60 equiv) in THF (25 mL) was added formic acid (5.47 g, 119 mmol, 24.0 equiv) and Pd(PPh<sub>3</sub>)<sub>4</sub> (572 mg, 0.495 mmol, 0.10 equiv). The resulting mixture was stirred for 4 h at 40 °C under nitrogen atmosphere. The mixture was cooled to room temperature, concentrated under reduced pressure and purified by silica gel column, eluting with EA/MeOH/H<sub>2</sub>O (10:2:1) to afford [(2R,3R,4R,5R)-5-(6-aminopurin-9-yl)-3-[(dibenzyloxyphosphoryl)oxy]-4-[(4-methoxyphenyl)methoxy]oxolan-2-yl]methoxyphosphonic acid (3.00 g, 83% yield) as a colorless oil. LCMS (ES, m/z): 726.20 [M-H]<sup>-</sup>.

**Step 9. ethyl 2-[2,4-bis(trifluoromethyl)phenyl]acetate**

To a stirred mixture of [2,4-bis(trifluoromethyl)phenyl]acetic acid (10.0 g, 36.7 mmol, 1 equiv) in EtOH (100 mL) was added con.H<sub>2</sub>SO<sub>4</sub> (1.80 g, 18.3 mmol, 0.50 equiv) dropwise at 0 °C. The mixture was stirred for 16 h at 50 °C. The reaction mixture was concentrated under reduced pressure. The residue was poured into ice/water (100 mL) and extracted with EtOAc (3 x 100 mL). The combined organic layers were washed with brine (100 mL), dried over anhydrous sodium sulfate and concentrated under reduced pressure. The crude product was purified by silica gel column, eluting with PE/THF (2:1) to afford ethyl 2-[2,4-bis(trifluoromethyl)phenyl]acetate (7.6 g, 69% yield) as a yellow oil. GCMS (ES, m/z): 300.06 [M].

##### **Step 10. ethyl 2-[2,4-bis(trifluoromethyl)phenyl]-2-methylpropanoate**

To a stirred mixture of ethyl 2-[2,4-bis(trifluoromethyl)phenyl]acetate (7.50 g, 25.0 mmol, 1.00 equiv) in DMF (70 mL) was added NaH (60%) (4.00 g, 100 mmol, 4.00 equiv) at 0 °C. The resulting mixture was stirred for 0.5 h at 0 °C. Then MeI (14.2 g, 100 mmol, 4.00 equiv) was added at 0 °C. The resulting mixture was stirred for 1 h at 25 °C. The mixture was slowly poured into ice/water (100 mL) and extracted with EtOAc (3 x 100 mL). The combined organic layers were washed with brine (100 mL), dried over anhydrous sodium sulfate and concentrated under reduced pressure. The crude product was purified by silica gel column, eluting with PE/EtOAc (10:1) to afford ethyl 2-[2,4-bis(trifluoromethyl)phenyl]-2-methylpropanoate (5.50 g, 67% yield) as a colorless oil. LCMS (ES, m/z): 329.00 [M+H]<sup>+</sup>.

##### **Step 11. 2-[2,4-bis(trifluoromethyl)phenyl]-2-methylpropan-1-ol**

To a stirred mixture of ethyl 2-[2,4-bis(trifluoromethyl)phenyl]-2-methylpropanoate (2.20 g, 6.70 mmol, 1.00 equiv) in THF (20 mL) was added LiAlH<sub>4</sub> (305 mg, 8.04 mmol, 1.20 equiv) at 0 °C. The resulting mixture was stirred for 2 h at 0 °C. The mixture was slowly poured into NH<sub>4</sub>Cl (sat., aq.) (40 mL) and extracted with EtOAc (3 x 70 mL). The combined organic layers were washed with brine (70 mL), dried over anhydrous sodium sulfate and concentrated under reduced pressure. The crude product was purified by silica gel column, eluting with PE/EA (3:1) to afford 2-[2,4-bis(trifluoromethyl)phenyl]-2-methylpropan-1-ol (1.0 g, 52% yield) as a colorless oil. GCMS (ES, m/z): 286.08 [M].

##### **Step 12. dibenzyl 2-[2,4-bis(trifluoromethyl)phenyl]-2-methylpropyl phosphate**

To a stirred solution of 2-[2,4-bis(trifluoromethyl)phenyl]-2-methylpropan-1-ol (900 mg, 3.14 mmol, 1.00 equiv) in DCM (10 mL) was added tetrazole (0.45 M in acetonitrile) (28.0 mL, 12.6 mmol, 4.00 equiv) and [bis(benzyloxy)phosphanyl]diisopropylamine (4.34 g, 12.6 mmol, 4.00 equiv) dropwise at 0 °C. The mixture was stirred for 1 h. Then m-CPBA (2.17 g, 12.6 mmol, 4.00 equiv) was added at 0 °C. Then the mixture was stirred for 1 h. The reaction was quenched with NaHCO<sub>3</sub> (sat., aq.) (30 mL). The resulting mixture was extracted with DCM (2 x 30 mL). The combined organic layers were washed with brine (30 mL), dried over anhydrous Na<sub>2</sub>SO<sub>4</sub>. After filtration, the filtrate was concentrated under reduced pressure. The crude product was purified by silica gel column, eluting with PE/EA (3:1) to afford dibenzyl 2-[2,4-bis(trifluoromethyl)phenyl]-2-methylpropyl phosphate (900 mg, 52% yield) as a colorless oil. LCMS (ES, m/z): 546.90 [M+H]<sup>+</sup>.

##### **Step 13. 2-[2,4-bis(trifluoromethyl)phenyl]-2-methylpropoxyphosphonic acid**

A mixture of dibenzyl 2-[2,4-bis(trifluoromethyl)phenyl]-2-methylpropyl phosphate (900 mg, 1.65 mmol, 1.00 equiv) and Pd/C (10%) (900 mg) in MeOH (8 mL) was stirred for 2 h at 25 °C under hydrogen atmosphere (~3 atm). The solids were filtered out and the filtrate was concentrated under reduced pressure to afford 2-[2,4-bis(trifluoromethyl)phenyl]-2-methylpropoxyphosphonic acid (600 mg, 99% yield) as a white solid. LCMS (ES, m/z): 365.05 [M-H]<sup>-</sup>.

**Step 14. [(2R,3R,4R,5R)-5-(6-aminopurin-9-yl)-3-[(dibenzyloxyphosphoryl)oxy]-4-[(4-methoxyphenyl)methoxy]oxolan-2-yl)methoxy({2-[2,4-bis(trifluoromethyl)phenyl]-2-methylpropoxy(hydroxy)phosphoryl}oxy)phosphinic acid**

To a stirred mixture of 2-[2,4-bis(trifluoromethyl)phenyl]-2-methylpropoxyphosphonic acid (604 mg, 1.65 mmol, 1.50 equiv) in DMF (5 mL) was added TEA (122 mg, 1.21 mmol, 1.10 equiv) and CDI (571 mg, 3.52 mmol, 3.20 equiv). The resulting mixture was stirred for 3 h at 25 °C under nitrogen atmosphere. Then a few drops of methanol were added. After 5 min, the reaction was concentrated under reduced pressure. A solution of [(2R,3R,4R,5R)-5-(6-aminopurin-9-yl)-3-[(dibenzyloxyphosphoryl)oxy]-4-[(4-methoxyphenyl)methoxy]oxolan-2-yl)methoxyphosphonic acid (800 mg, 1.10 mmol, 1.00 equiv) in DMF (5 mL) was added. The resulting mixture was stirred for 2 days at 25 °C under nitrogen atmosphere. The mixture was purified by reverse flash chromatography (column: C18, 120 g, 20-35 μm; mobile phase, A: water (25 mM NH<sub>4</sub>OAc) and B: ACN (0% to 80% in 50 min); detector: UV 254/ 220 nm). The collected fraction was concentrated to afford [(2R,3R,4R,5R)-5-(6-aminopurin-9-yl)-3-[(dibenzyloxyphosphoryl)oxy]-4-[(4-methoxyphenyl)methoxy]oxolan-2-yl)methoxy({2-[2,4-bis(trifluoromethyl)phenyl]-2-methylpropoxy(hydroxy)phosphoryl}oxy)phosphinic acid (250 mg, 21% yield) as a white solid. LCMS (ES, m/z): 1076.25 [M+H]<sup>+</sup>.

**Step 15. [(2R,3S,4R,5R)-5-(6-aminopurin-9-yl)-2-{{{2-[2,4-bis(trifluoromethyl)phenyl]-2-methylpropoxy(hydroxy)phosphoryl}oxy(hydroxy)phosphoryl}oxy)methyl}-4-hydroxyoxolan-3-yl]oxyphosphonic acid**

A mixture of [(2R,3R,4R,5R)-5-(6-aminopurin-9-yl)-3-[(dibenzyloxyphosphoryl)oxy]-4-[(4-methoxyphenyl)methoxy]oxolan-2-yl)methoxy({2-[2,4-bis(trifluoromethyl)phenyl]-2-methylpropoxy(hydroxy)phosphoryl}oxy)phosphinic acid (200 mg, 0.186 mmol, 1.00 equiv) and Pd/C (10%) (200 mg) in H<sub>2</sub>O (5 mL) was stirred for 6 h at 25 °C under hydrogen atmosphere (~ 3 atm). The solids were filtered out and the filtrate was concentrated under reduced pressure. The residue was purified by Prep-HPLC (column: XBridge Prep Phenyl OBD C18 Column, 19 x 250 mm, 5μm; mobile Phase, A: water (10 mmol/L NH<sub>4</sub>HCO<sub>3</sub>) and B: ACN (1% B to 30% B in 10min); flow rate: 25 mL/min; detector: 254/220 nm). The collected fraction was lyophilized to afford [(2R,3S,4R,5R)-5-(6-aminopurin-9-yl)-2-{{{2-[2,4-bis(trifluoromethyl)phenyl]-2-methylpropoxy(hydroxy)phosphoryl}oxy(hydroxy)phosphoryl}oxy)methyl}-4-hydroxyoxolan-3-yl]oxyphosphonic acid (11.1 mg, 7.7% yield) as a white solid.

**PH-ALTOS-360-0:** <sup>1</sup>H NMR (400 MHz, DMSO-d<sub>6</sub>) $\delta$ 8.33 (s, 1H), 8.06 (d, J = 2.1 Hz, 1H), 7.82 (s, 1H), 7.52 (s, 2H), 5.97 (d, J = 6.1 Hz, 1H), 4.70 – 4.50 (m, 2H), 4.44 (s, 1H), 4.30 – 4.06 (m, 2H), 3.94 (d, J = 4.5 Hz, 2H), 1.24 (d, J = 13.2 Hz, 6H).

**LCMS** (ES, m/z): 775.90 [M+H]<sup>+</sup>, **HRMS** Calcd for C<sub>22</sub>H<sub>26</sub>F<sub>6</sub>N<sub>5</sub>O<sub>13</sub>P<sub>3</sub> [M+H]<sup>+</sup> m/z 776.0644, Found 776.0725.

*Synthesis of CoA-A7-2*

**[(2R,3S,4R,5R)-5-(6-aminopurin-9-yl)-2-{{{2-[2,4-bis(trifluoromethyl)phenyl]ethoxy(hydroxy)phosphoryl}oxy(hydroxy)phosphoryl}oxy)methyl}-4-hydroxyoxolan-3-yl]oxyphosphonic acid (PH-ALTOS-361-0)**

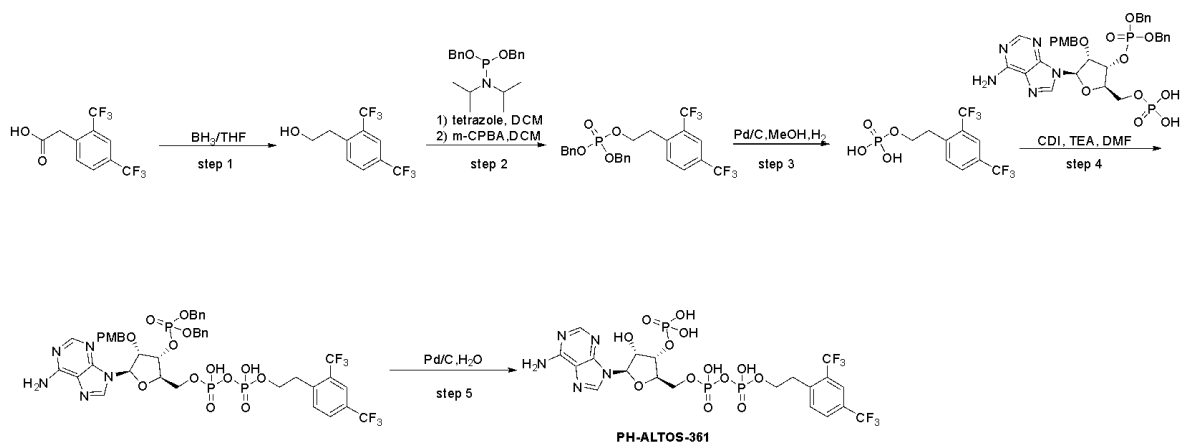

#### Step 1. 2-[2,4-bis(trifluoromethyl)phenyl]ethanol

A solution of 2-[2,4-bis(trifluoromethyl)phenyl]acetic acid (5.00 g, 18.4 mmol, 1.00 equiv) in THF (60 mL) was added $\text{BH}_3$ (1.0 M in THF) (73.6 mL, 73.6 mmol, 4.00 equiv) dropwise at 0 °C. Then the reaction was stirred for 3 h at 60 °C. The mixture was cooled to room temperature, quenched with MeOH (100 mL) at 0 °C. The combined organic layers were concentrated under reduced pressure. The residue was purified by silica gel column, eluting with PE/EA (2:1) to afford 2-[2,4-bis(trifluoromethyl)phenyl]ethanol (4.00 g, 84% yield) as a colorless oil. LCMS (ES, m/z): 291.30 $[\text{M}+\text{H}]^+$.

#### Step 2. dibenzyl 2-[2,4-bis(trifluoromethyl)phenyl]ethyl phosphate

To a stirred solution of 2-[2,4-bis(trifluoromethyl)phenyl]ethanol (3.00 g, 11.6 mmol, 1.00 equiv) in DCM (35 mL) was added tetrazole (0.45 M in acetonitrile) (103 mL, 46.4 mmol, 4.00 equiv) and [bis(benzyloxy)phosphanyl]diisopropylamine (16.1 g, 46.4 mmol, 4.00 equiv) dropwise at 0 °C. The mixture was stirred for 1 h at 0 °C. Then m-CPBA (8.02 g, 46.4 mmol, 4.00 equiv) was added. Then the mixture was stirred for 1 h at 0 °C. The reaction was quenched with $\text{NaHCO}_3$ (sat., aq.) (40 mL). The resulting mixture was extracted with DCM (2 x 70 mL). The combined organic layers were washed with brine (70 mL), dried over anhydrous $\text{Na}_2\text{SO}_4$. After filtration, the filtrate was concentrated under reduced pressure. The combined organic layers were concentrated under reduced pressure and purified by silica gel column, eluting with PE/EA (1:1) to afford dibenzyl 2-[2,4-bis(trifluoromethyl)phenyl]ethyl phosphate (4.50 g, 75% yield) as a colorless oil. LCMS (ES, m/z): 519.10 $[\text{M}+\text{H}]^+$.

#### Step 3. 2-[2,4-bis(trifluoromethyl)phenyl]ethoxyphosphonic acid

A mixture of dibenzyl 2-[2,4-bis(trifluoromethyl)phenyl]ethyl phosphate (4.50 g, 8.68 mmol, 1.00 equiv) and Pd/C (10%) (2.30 g) in MeOH (25 mL) was stirred for 16 h at 25 °C under hydrogen atmosphere (~ 3 atm). The solids were filtered out and the filtrate was concentrated under reduced pressure to afford 2-[2,4-bis(trifluoromethyl)phenyl]ethoxyphosphonic acid (2.00 g, 68% yield) as a colorless oil. LCMS (ES, m/z): 338.95 $[\text{M}+\text{H}]^+$.

#### Step 4. $\{[(2\text{R},3\text{R},4\text{R},5\text{R})\text{-}5\text{-(6-aminopurin-9-yl)-3-}[(\text{dibenzyl oxyphosphoryl})\text{oxy}]\text{-4-}[(4\text{-methoxyphenyl})\text{methoxy}]\text{oxolan-2-yl)]\text{methoxy(hydroxy)phosphoryl}\text{oxy}2\text{-[2,4-bis(trifluoromethyl)phenyl]ethoxy}\}\text{phosphinic acid}$

To a stirred mixture of 2-[2,4-bis(trifluoromethyl)phenyl]ethoxyphosphonic acid (349 mg, 1.03 mmol, 1.50 equiv) in DMF (4 mL) was added TEA (76.5 mg, 0.757 mmol, 1.10 equiv) and CDI

(571 mg, 3.52 mmol, 3.20 equiv). The resulting mixture was stirred for 3 h at 25 °C under nitrogen atmosphere. Then a few drops of methanol were added. After 5 min, the reaction was concentrated under reduced pressure. The solution of [(2R,3R,4R,5R)-5-(6-aminopurin-9-yl)-3-[(dibenzyloxyphosphoryl)oxy]-4-[(4-methoxyphenyl)methoxy]oxolan-2-yl]methoxyphosphonic acid (600 mg, 0.688 mmol, 1.00 equiv) in DMF (4 mL) was added. The resulting mixture was stirred for 2 days at 25 °C under nitrogen atmosphere. The mixture was purified by reverse flash chromatography (column: C18, 90 g, 20-35 μm; mobile phase, A, water (25 mM $\text{NH}_4\text{OAc}$) and B: ACN (0% to 80% in 50 min); detector: UV 254/220 nm). The collected fraction was concentrated to afford {[(2R,3R,4R,5R)-5-(6-aminopurin-9-yl)-3-[(dibenzyloxyphosphoryl)oxy]-4-[(4-methoxyphenyl)methoxy]oxolan-2-yl]methoxy(hydroxy)phosphoryl}oxy(2-[2,4-bis(trifluoromethyl)phenyl]ethoxy)phosphonic acid (400 mg, 56% yield) as a white solid. LCMS (ES, m/z): 1048.20 $[\text{M}+\text{H}]^+$.

**Step 5. [(2R,3S,4R,5R)-5-(6-aminopurin-9-yl)-2-{{{2-[2,4-bis(trifluoromethyl)phenyl]ethoxy(hydroxy)phosphoryl}oxy(hydroxy)phosphoryl}oxy)methyl}-4-hydroxyoxolan-3-yl]oxyphosphonic acid**

A mixture of {[(2R,3R,4R,5R)-5-(6-aminopurin-9-yl)-3-[(dibenzyloxyphosphoryl)oxy]-4-[(4-methoxyphenyl)methoxy]oxolan-2-yl]methoxy(hydroxy)phosphoryl}oxy(2-[2,4-bis(trifluoromethyl)phenyl]ethoxy)phosphonic acid (200 mg, 0.191 mmol, 1.00 equiv) and Pd/C (10%) (200 mg) in $\text{H}_2\text{O}$ (3 mL) was stirred for 6 h at 25 °C under hydrogen atmosphere (~ 3 atm). The solids were filtered out and the filtrate was concentrated under reduced pressure. The crude product was purified by Prep-HPLC (column: XBridge Prep Phenyl OBD, 19 x 250 mm, 5μm; mobile Phase, A: water (10 mmol/L $\text{NH}_4\text{HCO}_3$) and B: ACN (1% to 30% in 8 min); flow rate: 25 mL/min; detector: 254/220 nm). The collected fraction was lyophilized to afford [(2R,3S,4R,5R)-5-(6-aminopurin-9-yl)-2-{{{2-[2,4-bis(trifluoromethyl)phenyl]ethoxy(hydroxy)phosphoryl}oxy(hydroxy)phosphoryl}oxy)methyl}-4-hydroxyoxolan-3-yl]oxyphosphonic acid (16.5 mg, 11.6% yield) as a white solid.

**PH-ALTOS-361-0:** $^1\text{H}$ NMR (400 MHz, $\text{DMSO}-d_6$) $\delta$ 8.34 (s, 1H), 7.99 (s, 1H), 7.70 (s, 1H), 7.48 (d, $J$ = 8.2 Hz, 1H), 7.21 (d, $J$ = 8.1 Hz, 1H), 6.02 – 5.89 (m, 1H), 4.85 – 4.82 (m, 2H), 4.50 (s, 1H), 4.31 – 4.12 (m, 2H), 4.05 – 3.83 (m, 2H), 3.00 – 2.80 (m, 2H).

$^{13}\text{C}$ NMR (101 MHz, $\text{D}_2\text{O}+\text{DMSO}$): $\delta$ 152.69, 150.30, 146.59, 138.11, 137.31, 129.88, 126.16, 125.62, 122.30, 120.57, 120.28, 115.72, 84.04, 80.99, 71.50, 71.33, 63.21, 63.07, 30.27.

**LCMS** (ES, m/z): 747.90. $[\text{M}+\text{H}]^+$, **HRMS** Calcd for $\text{C}_{20}\text{H}_{22}\text{F}_6\text{N}_5\text{O}_{13}\text{P}_3$ $[\text{M}+\text{H}]^+$ m/z 748.0331, Found 748.0402.

### References

1. Replogle, J. M. *et al.* Maximizing CRISPRi efficacy and accessibility with dual-sgRNA libraries and optimal effectors. *Elife* **11**, e81856 (2022).

2. Lamm, L. *et al.* MemBrain: A deep learning-aided pipeline for detection of membrane proteins in Cryo-electron tomograms. *Comput Methods Programs Biomed* **224**, 106990 (2022).
3. Li, Y. *et al.* Structural insights into human ABCD3-mediated peroxisomal acyl-CoA translocation. *Cell Discov.* **10**, 92 (2024).
4. Abu-Remaileh, M. *et al.* Lysosomal metabolomics reveals V-ATPase- and mTOR-dependent regulation of amino acid efflux from lysosomes. *Science* **358**, 807–813 (2017).
5. Walters, B. T., Ricciuti, A., Mayne, L. & Englander, S. W. Minimizing back exchange in the hydrogen exchange-mass spectrometry experiment. *J. Am. Soc. Mass Spectrom.* **23**, 2132–2139 (2012).
6. Mayne, L. *et al.* Many overlapping peptides for protein hydrogen exchange experiments by the fragment separation-mass spectrometry method. *J. Am. Soc. Mass Spectrom.* **22**, 1898–1905 (2011).
7. Masson, G. R. *et al.* Recommendations for performing, interpreting and reporting hydrogen deuterium exchange mass spectrometry (HDX-MS) experiments. *Nat. Methods* **16**, 595–602 (2019).
